# CaMKII activity spreads by inter-holoenzyme phosphorylation

**DOI:** 10.1101/2022.08.03.502606

**Authors:** Iva Lučić, Léonie Héluin, Pin-Lian Jiang, Alejandro G. Castro Scalise, Cong Wang, Andreas Franz, Markus Wahl, Fan Liu, Andrew J.R. Plested

## Abstract

The dodecameric protein kinase CaMKII is expressed throughout the body. The alpha isoform is responsible for synaptic plasticity and participates in memory through its phosphorylation of synaptic proteins. Its elaborate subunit organization and propensity for autophosphorylation allow it to preserve neuronal plasticity across space and time. The prevailing hypothesis for the spread of CaMKII activity, involving shuffling of subunits between activated and naïve holoenzymes, is broadly termed subunit exchange. In contrast to the expectations of previous work, we found little evidence for subunit exchange upon activation, and no effect of restraining subunits to their parent holoenzymes. Rather, mass photometry, crosslinking mass spectrometry, single molecule TIRF microscopy and biochemical assays identify inter-holoenzyme phosphorylation (IHP) as the mechanism for spreading phosphorylation. The transient, activity-dependent formation of groups of holoenzymes is well suited to the speed of neuronal activity. Our results place fundamental limits on the activation mechanism of this kinase.

## Introduction

The Calcium-calmodulin-dependent protein kinase 2 (CaMKII) is encoded by 4 genes CaMKII *A*, *B*, *G*, and *D* (Tombes, Faison and Turbeville, 2003). Whilst the γ and δ protein isoforms are expressed throughout the body, CaMKIIα and β show brain-specific expression where they are amongst the most abundant proteins in the neuronal cytosol (Cheng *et al.*, 2006; Sheng and Kim, 2011). CaMKII is an oligomeric structure of mostly 12 or 14 subunits (Rosenberg *et al.*, 2006; Chao *et al.*, 2011; Bhattacharyya *et al.*, 2016; Buonarati *et al.*, 2021). The individual subunits associate into holoenzymes through avid hub domain interactions. The hub connects to the N-terminal kinase domains through a variable-length linker and regulatory domain. The peripheral location of the kinase domains presumably facilitates interaction with each other and substrates (Chao *et al.*, 2011; Myers *et al.*, 2017; Buonarati *et al.*, 2021) (Figure 1A).

**Figure 1.**
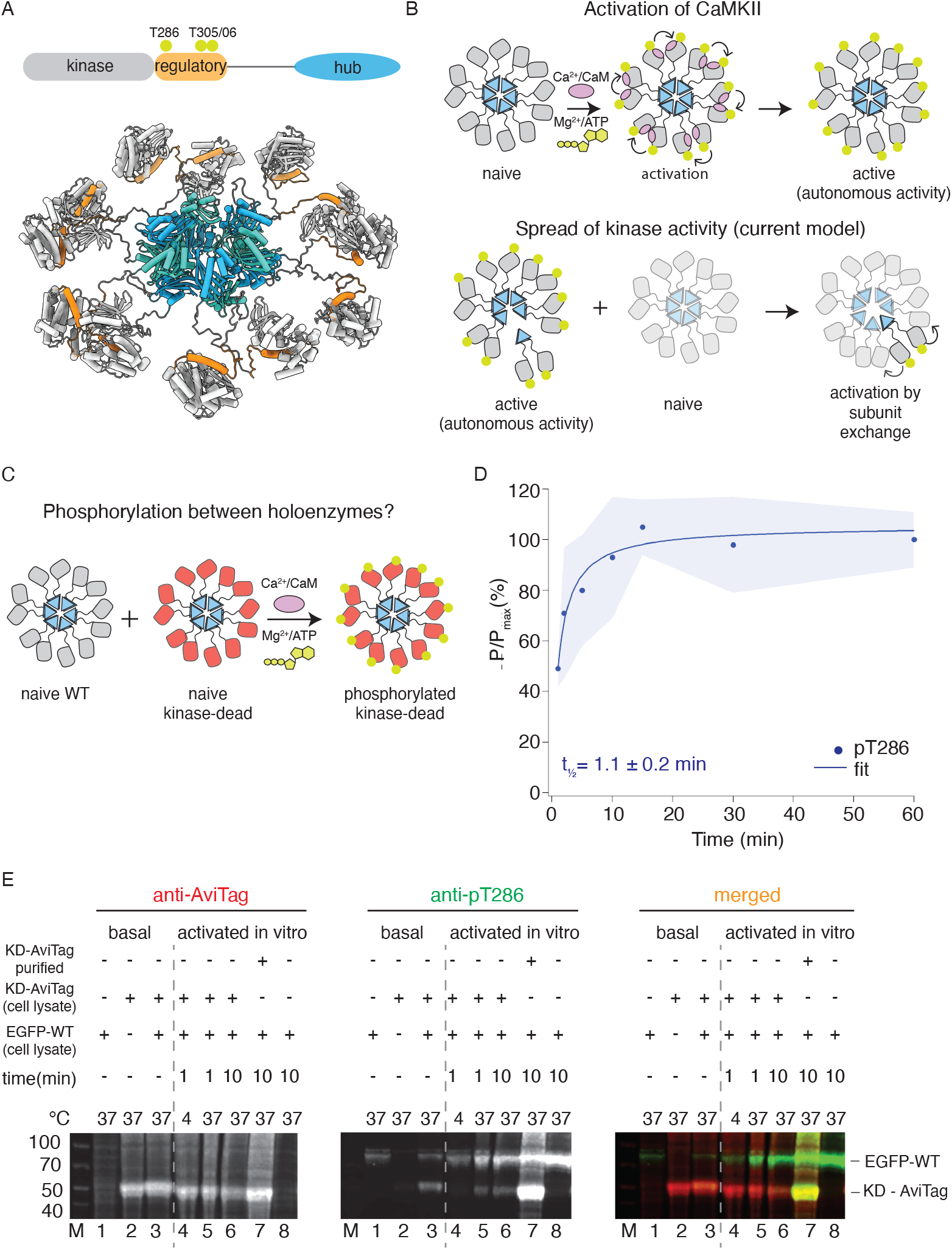
CaMKIIWT phosphorylates CaMKII^KD^. (A) Schematic representation of CaMKIIα domain arrangement (top). Yellow circles indicate phosphorylation sites. CaMKIIα holoenzyme structure (bottom, PDB:5u6y). (B) Cartoon representation of activation of CaMKII by calcium:calmodulin (top) and proposed mechanism for spread of kinase activity (bottom). Yellow circles indicate phosphorylation sites. C) Schematic representation of experiment performed in panel (D). D) Kinase activity of CaMKII^WT^ (10 nM) against CaMKII^KD^ (4 μM). Half-time of maximum phosphorylation (t½ = 1.1 ± 0.2 min) determined by western blot, using an antibody against pT286. (E) Kinase activity of EGFP-CaMKII^WT^ and CaMKII^KD^-AviTag in HEK cells and HEK cell lysates. “Basal”: activity in cells; “activated in vitro”: activity in cell lysates supplemented with 1μM purified Ca^2+^:CaM and 0.5mM ATP:Mg^2+^. Lane 1 – autophosphorylation of EGFP-CaMKII^WT^ in singly transfected cells, lane 2 – autophosphorylation of CaMKII^KD^-AviTag in singly transfected cells, lane 3 – autophosphorylation of EGFP-CaMKII^WT^ and CaMKII^KD^-AviTag in co-transfected cells, lane 4 – stimulated kinase activity of EGFP-CaMKII^WT^ from lysate in lane 1 and CaMKII^KD^-AviTag from lysate in lane 2, incubated together at 4°C for 1 min, lane 5 - stimulated kinase activity of EGFP-CaMKII^WT^ from lysate in lane 1 and CaMKII^KD^-AviTag from lysate in lane 2, incubated together at 37°C for 1 min, lane 6 - stimulated kinase activity of EGFP-CaMKII^WT^ from lysate in lane 1 and CaMKII^KD^-AviTag from lysate in lane 2, incubated together at 37°C for 10 min, lane 7 - stimulated kinase activity of EGFP-CaMKII^WT^ from lysate in lane 1 and 8 μM purified CaMKII^KD^-AviTag, incubated together at 37°C for 10 min, lane 8 - stimulated kinase activity of EGFP-CaMKII^WT^ from lysate in lane 1 37°C for 10 min. The ability of CaMKII^WT^ to phosphorylate CaMKII^KD^ is in contrast to previous reports, which failed to detect any inter-holoenzyme phosphorylation of CaMKII^WT^ or CaMKII^KD^ (Miller, S. G. & Kennedy, 1986; Hanson *et al.*, 1994; Rich and Schulman, 1998). Most previous work used either short incubation time (30 - 60 s) or low incubation temperature (4°C) or both; these likely prevented CaMKII activity. Another difficulty could be the lower expression of CaMKII^KD^ in cell lines, compared to dendritic spines where CaMKII concentration is estimated to be between 20 and 100 μM (Otmakhov and Lisman, 2012). This discrepancy means that *in vitro* experiments, which allowed us to use 4-8 μM CaMKII^KD^ protein, are more representative of the crowded environment of neurons.

In basal conditions, CaMKII is kept silent because the regulatory domain blocks access to the substrate binding cleft (Chao *et al.*, 2011; Myers *et al.*, 2017). A canonical form of synaptic plasticity relates to the recruitment of active CaMKII to synapses following calcium (Ca^2+^) influx via the N-methyl-D-aspartate (NMDA)-type glutamate receptors (Lisman, Schulman and Cline, 2002; Lisman, Yasuda and Raghavachari, 2012). Binding of Ca^2+^-bound calmodulin to the regulatory domain activates CaMKII by liberating its substrate binding site. Auto-phosphorylation at T286 (CaMKIIα numbering) in the regulatory domain prevents rebinding of the regulatory domain back to the kinase and allows substrate engagement. At this stage CaMKII no longer needs Ca^2+^:CaM in order to be active. This mode of operation, which allows CaMKII to remain active after the Ca^2+^ signal (and the associated Calmodulin binding) has finished, is referred to as autonomous activity (Lou, Lloyd and Schulman, 1986; Miller, S. G. & Kennedy, 1986; Lou and Schulman, 1989; Waldmann, Hanson and Schulman, 1990; Chang *et al.*, 2019) (Figure 1B). Auto-phosphorylation at two other sites in the regulatory domain (T305 and T306) prevents further binding of Ca^2+^:CaM (Lou and Schulman, 1989; Elgersma *et al.*, 2002). Autonomous activity, was first envisaged as a desirable general mechanism for memory storage by Crick before CaMKII was identified (Crick, 1984). The oligomeric structure of CaMKII might aid autonomous activity. In particular, the intra-holoenzyme autophosphorylation might provide a reservoir against phosphatase attack, because any dephosphorylated subunits can be “revitalized” by their neighbours. CaMKIIα organizes a range of functions related to the plasticity of neuronal structure and function. CaMKII is concentrated in dendritic spines, where it bundles actin and phosphorylates its targets such as the α-amino-3-hydroxy-5-methyl-4-isoxazolepropionic acid (AMPA) receptor auxiliary proteins (Coultrap and Bayer, 2012; Herring and Nicoll, 2016; Bayer and Schulman, 2019)

A further key concept, that CaMKII phosphorylation is spread throughout a naïve population of holoenzymes by physical exchange of subunits (that is, active subunits are swapped into naïve holoenzymes, which then auto-phosphorylate around the ring) is grounded on several observations. First, early reports suggested that naïve holoenzymes do not get phosphorylated when mixed with activated holoenzymes (Miller, S. G. & Kennedy, 1986; Hanson *et al.*, 1994; Rich and Schulman, 1998; Chao *et al.*, 2011). However, these experiments were carried out at low temperature and with short mixing times and at low concentrations that do not mimic the physiological realm of neurons. Second, labelled subunits accumulate into clusters upon activation, as observed in total internal reflection fluorescence (TIRF) microscopy, but the limited resolution of optical microscopy means that clusters could contain multiple holoenzymes (Stratton *et al.*, 2014; Bhattacharyya *et al.*, 2016). Third, peptides derived from the regulatory domain can selectively break up hubs during electrospray mass spectrometry, but not in solution (Karandur *et al.*, 2020).

In this work, we provide data that support a new model in which inter-holoenzyme phosphorylation (IHP) is the predominant mechanism by which CaMKII activity propagates. Subunit mixing experiments, using a panel of approaches and labels, confirm that restraining subunits through the hub domain has no effect on the accumulation of phosphorylation, that holoenzymes reversibly accumulate into clusters upon activation, and that whilst kinase domains interact promiscuously during autophosphorylation, the hub domains do not mix.

## Results

### CaMKII subunits from different holoenzymes can trans-autophosphorylate

Seeking to validate the subunit exchange as a mechanism for spread of kinase activity in CaMKII, we first tested the ability of wild-type CaMKII (CaMKII^WT^) to phosphorylate kinase-dead CaMKII (K42R and D135N double mutant; CaMKII^KD^) *in vitro* using purified proteins (Figure 1 C and D; Figure S1 and S2). We used CaMKII^KD^ double mutant, because in our hands, the single mutations did not completely abolish kinase activity.

We readily detected phosphorylation of CaMKIIKD by either immunoblotting (Figure 1D and Figure S1A, B) against phosphorylated T286, or incorporation of radioactive ^32^P (Figure S2). In these experiments we used 10 nM CaMKII^WT^ and 4μM CaMKII^KD^, (400-fold excess of KD) in the presence of Ca^2+^:CaM and ATP:Mg^2+^. The WT and KD proteins are the same size (monomer Mw = 55 kDa), which forbids their separation on SDS-polyacrylamide gel electrophoresis (PAGE). However, at 10 nM CaMKII^WT^ phosphorylation was undetectable on a gel (Figure S1D), meaning that the phosphorylation signal we detected is overwhelmingly from the 400-fold more abundant kinase-dead protein. The half-times of maximal phosphorylation are different, being 1.1 min for the immunoblotting (Figures 1D and S1E) and 4 mins for the P^32^ incorporation (Figure S2C), perhaps because T286 is phosphorylated early compared to the entire cohort of phosphosites incorporating ^32^P.

Next, we tested phosphorylation of CaMKII in HEK cells and in HEK cell lysates (Figure 1E). We expressed EGFP-tagged CaMKII^WT^ alone, CaMKII^KD^-AviTag alone, or both proteins together, and sought to detect phosphorylation on T286 by western blotting. When expressed separately, we could observe phosphorylation on CaMKII^WT^ but not on CaMKII^KD^ (Figure 1E, lanes 1 and 2). We could, however, detect phosphorylation on CaMKII^KD^ when it was co-expressed with EGFP- CaMKII^WT^ (Figure 1E, lane 3). A confounding explanation for this is that WT and KD subunits might assemble into common holoenzymes during co-expression in cells, allowing KD subunits to be auto-phosphorylated in an intra-holoenzyme context (Hanson *et al.*, 1994). Next, we incubated HEK cell lysate expressing EGFP-tagged CaMKII^WT^ alone with HEK cell lysate expressing CaMKII^KD^-AviTag alone, in the presence of 0.5 μM purified Calmodulin as well as 0.5mM ATP: Mg^2+^ (Figure 1E, lanes 4-6), over different incubation temperatures and times. By incubating lysates with separately expressed WT and KD proteins, we wanted to exclude the possibility of the two proteins assembling into the same holoenzymes during translation, which is to be expected when they are co-expressed. Similar to previous reports (Hanson *et al.*, 1994; Rich and Schulman, 1998) we failed to detect phosphorylation on CaMKII^KD^-AviTag after 1 min incubation at 4°C (Figure 1E, lane 4). In contrast, the phospho-signal was readily detected on CaMKII^KD^-AviTag at 37°C (Figure 1E, lanes 5 and 6), indicating that temperature and incubation time play an important role in enzymatic activity of CaMKII. Finally, when we incubated HEK cell lysate expressing EGFP-tagged CaMKII^WT^ with 4μM CaMKII^KD^-AviTag purified from *E. coli*, we could detect robust phosphorylation on CaMKII^KD^ protein (Figure 1E, lane 7), indicating the importance of CaMKII concentration in this process. Having in mind high dendritic CaMKII concentrations, this experiment indicates that crowding of CaMKII holoenzymes, rather than subunit exchange *per se*, might be important for spreading phosphorylation.

### Holoenzyme constituents do not rearrange into other holoenzymes during activation

Having established that active CaMKII readily phosphorylates subunits from naïve holoenzymes, we next used a genetically-encoded photocrosslinker to test whether the phosphorylation of CaMKII^KD^ could still be detected if active CaMKII is restricted to its respective holoenzymes. If phosphorylation spreads by subunit exchange between holoenzymes, then restraining subunits is expected to abolish, or at least slow down, the phosphorylation of CaMKIIKD. We designed several CaMKII mutants in which we placed a BzF residue on different positions in the hub domain, using genetic code expansion technology (Figure 2A) (Chin *et al.*, 2002; Ye *et al.*, 2008; Klippenstein *et al.*, 2014). BzF is an unnatural amino acid which, when exposed to UV light, covalently crosslinks to residues at ~ 5 Å distant (Kauer *et al.*, 1986; Young *et al.*, 2010). Exposing hub domain BzF mutants to UV light of 365nm wavelength generated higher order oligomeric bands on an SDS-PAGE gel (Figure 2B). We chose the F394BzF mutation for downstream phosphorylation assays, because this mutant behaved like the WT and KD protein in terms of dodecameric assembly and had similar activity to WT (Figure S3).

**Figure 2.**
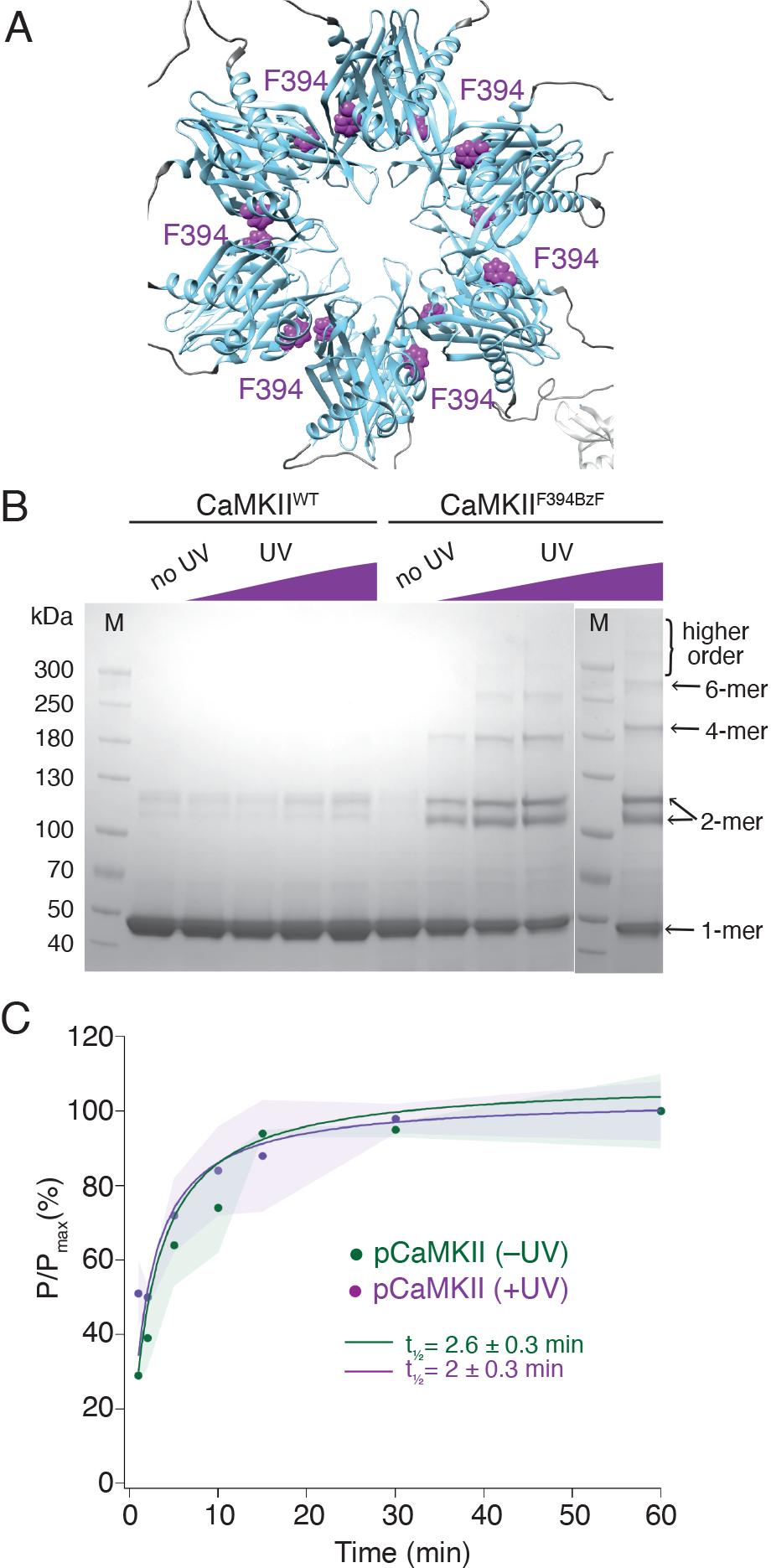
Restricting CaMKII mobility does not slow down kinase reaction. (A) Position of F394 residue (purple) in the hub domain of CaMKIIα (PDB: 5u6y), showing orientation towards the interface between adjacent hub domains within one hub ring. (B) Coomassie stained gel of UV-induced crosslinking of CaMKII^F394BzF^ (right) to higher order oligomers, and CaMKII^WT^ (left) as a control. Purple ramp indicates increasing UV exposure. (C) Kinase activity of UV treated (+UV) and untreated (−UV) CaMKII^F394BzF^ against CaMKII^KD^-AviTag. Phosphorylation was measured as incorporation of radioactive AT^32^P at phosphorylation sites on CaMKII^KD^-AviTag (pCaMKII). Half-times of maximum phosphorylation are determined to t½ = 2.6 ± 0.3 min for untreated and t½ = 2 ± 0.3 min for UV treated CaMKII^F394BzF^.

We performed two sets of experiments using CaMKII^F394BzF^ and CaMKII^KD^. First, we monitored the phosphorylation kinetics of CaMKIIKD when incubated with CaMKII^F394BzF^, which was either previously treated with UV light or not (Figure 2C). We assume that UV treatment crosslinks monomers, imprisoning them within their original holoenzymes and forcing them to phosphorylate CaMKII^KD^ in separate holoenzymes in trans, in order to spread activity. Figure 2C shows that the UV treatment of CaMKII^F394BzF^ before the incubation with CaMKII^KD^ and activation stimuli did not slow down the phosphorylation kinetics of CaMKII^KD^. Regardless of whether CaMKII^F394BzF^ was crosslinked or not, phosphorylation of CaMKII^KD^ proceeded with a half time of 2 min. We performed crosslinking of CaMKII^F394BzF^ at concentration of 8 μM, and then diluted it to 10 nM for the phosphorylation assay, whereas we kept the CaMKII^KD^ concentration at 4 μM in the phosphorylation assay, again assuring that the overwhelming majority of the signal from the autoradiography gel is from the kinase-dead protein (Figure S3). The same lack of effect of UV induced crosslinking was also observed for another BzF hub domain mutant, CaMKII^H418BzF^ (Figure S4). Finally, restricting the ability of CaMKII^F394BzF^ to dissociate from holoenzymes also failed to alter phosphorylation of the well-known Syntide-2 substrate (Hashimoto and Soderling, 1987) (Figure S5).

Since active forms of CaMKII can efficiently phosphorylate CaMKII^KD^ *in vitro* (Figure 1D, 2C and S4), we used the F394BzF mutant to investigate whether we could capture CaMKII^KD^ subunits with BzF crosslinks during phosphorylation (Figure 3A). For this purpose, we incubated 4 μM CaMKII^F394BzF^ with 4 μM CaMKII^KD^-AviTag under activating conditions at 37°C for 1h. After this substantial mixing period, and at which point the phosphorylation reaction had already plateaued, we treated the samples with UV light to promote BzF crosslinking. We reasoned that if the subunit exchange were to have occurred during phosphorylation of CaMKII^KD^ by CaMKII^F394BzF^, some fraction of CaMKII^KD^ protein should be captured by UV-induced crosslinking, due to mixing of CaMKII^F394BzF^ and CaMKII^KD^ subunits. The kinase-dead protein had an AviTag on the C-terminus, right after the hub domain. We could not detect the AviTag signal in higher order oligomeric bands by western blot (Figure 3B), whereas higher bands were visible on the Coomassie stained gel, coming exclusively from CaMKII^F394BzF^ crosslinking (Figure S6A). Despite evident covalent crosslinking by BzF, no AviTagged CaMKII^KD^ protein was captured by the crosslinks. This result indicated that although CaMKII^KD^ can be phosphorylated by CaMKII^F394BzF^, these proteins do not mix within the same holoenzymes. To further ensure that this failure to detect subunit exchange was not an artifact of the KD mutant, we repeated the same experiment using CaMKII^WT^-AviTag and CaMKII^F394BzF^. Again, no AviTag labelled protein was enriched in higher order oligomeric bands on the blot after incubation with CaMKII^F394BzF^ and subsequent UV-treatment (Figure S6B).

**Figure 3.**
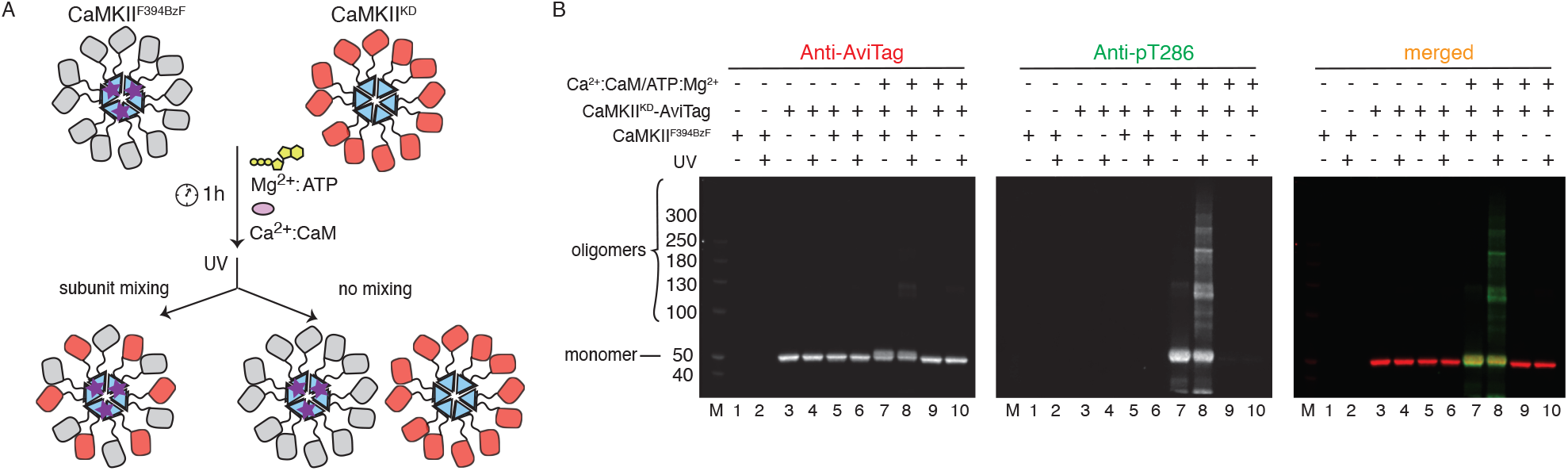
CaMKII holoenzymes do not mix during activation. (A) Schematic representation of the experiment performed in panel (B) and possible outcomes. (B) Western blot detection of possible CaMKII^KD^-AviTag incorporation in CaMKII^F394BzF^ holoenzymes. Lane 1 – CaMKII^F394BzF^, lane 2 - CaMKII^F394BzF^ treated with UV, lane 3 - CaMKII^KD^-AviTag, lane 4 - CaMKII^KD^-AviTag treated with UV, lane 5 - CaMKII^F394BzF^ incubated with CaMKII^KD^-AviTag, lane 6 - CaMKII^F394BzF^ incubated with CaMKII^KD^-AviTag, then UV treated, lane 7 - CaMKII^F394BzF^ incubated with CaMKII^KD^-AviTag and activation stimuli (Ca^2+^:CaM and Mg^2+^:ATP), lane 8 - CaMKII^F394BzF^ incubated with CaMKII^KD^-AviTag and activation stimuli (Ca^2+^:CaM and Mg^2+^:ATP), then UV treated, lane 9 - CaMKII^KD^-AviTag incubated with activation stimuli (Ca^2+^:CaM and Mg^2+^:ATP), lane 10 - CaMKII^KD^-AviTag incubated with activation stimuli (Ca^2+^:CaM and Mg^2+^:ATP), then UV treated.

### Inter-holoenzyme contacts between kinase domains but not hub domains are detected during activation

Although CaMKII can perform phosphorylation of subunits coming from different holoenzymes (Figures 1D, 2C, S4), we failed to detect mixing of subunits during this process, using UV-induced crosslinking of BzF CaMKII mutants (Figure 3B and S6B). Since the prevailing model for spread of kinase activity of CaMKII argues for subunit exchange between holoenzymes (Stratton *et al.*, 2014; Bhattacharyya *et al.*, 2016), we inspected the mixing of holoenzymes using crosslinking detected by mass-spectrometry. Here we used CaMKII^WT^ from *E. coli* grown under regular conditions, and mixed it with CaMKII^WT^ from *E.coli* grown in ^15^N supplemented media. In the latter case, we obtained about 97% ^15^N incorporation in CaMKII. After incubation of ^14^N-and ^15^N-incorporated CaMKII^WT^ with activation stimuli, we added Disuccinimidyl suberate (DSS) in order to crosslink the proteins in activated states. We also performed this reaction on mixed holoenzymes in basal conditions. At two time points, 30 minutes and 150 minutes, the samples were digested and subjected to liquid chromatography mass spectrometry (LC-MS, Fig. 4A, *n* = 2 replicates). Crosslinked peptides detected from mass spectra were classified according to the ratio (*R*) between the intensities of heterotypic (mixed isotopic, that is I_14N:15N_ and I_15N:14N_) and homotypic (uni-isotopic I_14N:14N_ and I_15N:15N_) peaks (see Methods) (Figure S7). While some crosslinks were only detected between homo-isotypes, others were found (generally with a lower frequency) between both hetero-isotypes and homo-isotypes. The low ratio for heterotypic crosslinks detected is consistent with intra-holoenzyme contacts between subunits being more frequent than contacts between holoenzymes, as expected from local concentration effects.

**Figure 4.**
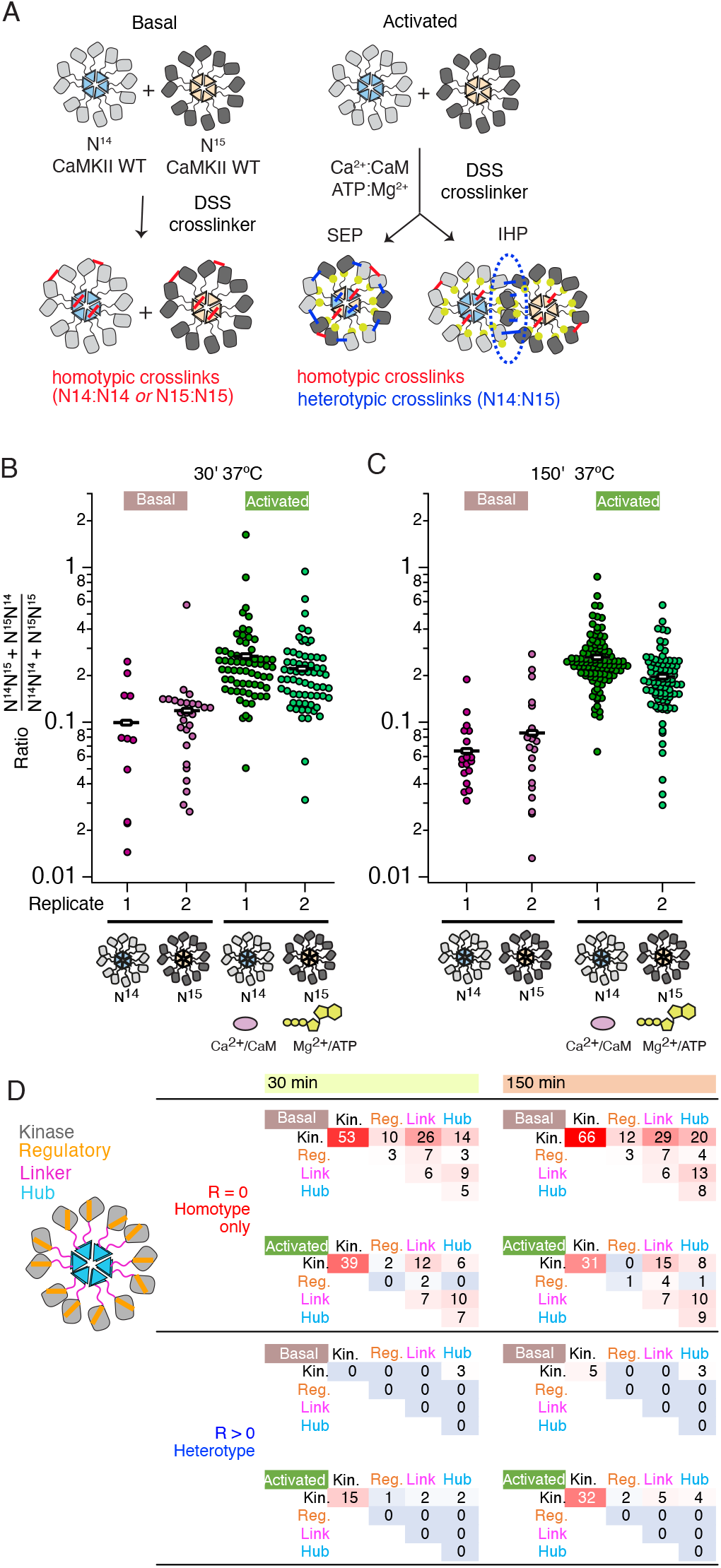
Crosslinking mass spectrometry reveals inter-holoenzyme kinase domain contacts during activation. (A) Schematic representation of crosslinking experiments and expected outcomes. In the case of subunit exchange, a flat profile of intersubunit crosslinks is expected, whereas for interholoenzyme phosphorylation, a bias towards kinase domain crosslinks is expected. (B) Heterotypic interactions plotted as R values of 2 independent replicates in basal and activating conditions. Incubation time 30 min prior to addition of DSS crosslink. (C) As for panel (B) but with incubation time of 150 min prior to addition of DSS crosslink. (D) Heat map indicating the number of homotypic (upper) and heterotypic (lower) crosslinks by domain under basal and activating conditions, after 30 and 150 min of incubation. Only peptides that were identified in both samples for each condition were counted in the heat map.

Across the four conditions, we classified more heterotypic crosslinks after longer incubations (Fig. 4B and C). Activating conditions increased the number and average ratio of heterotypic crosslinks (Fig. 4B and C, Supplementary Tables 1 and 2) for both 30’ and 150’ intervals. Next, we counted the number of crosslinks and constructed heatmaps of the crosslinking profiles by domain including only crosslinks found in both replicates (Figure 4D). The differential pattern of crosslinks between homo- and hetero-isotypes gives insight into the arrangement of subunits. Heterotypic crosslinks were restricted to the peripheral kinase domains, and were practically absent in basal conditions (Figure 4D and Supplementary Tables 3 and 4). In contrast, homotypic crosslinks preferred the kinase domain but otherwise showed a relatively flat profile (Figure 4D). Each subdomain (kinase, regulatory segment, linker and hub) readily linked with the others, consistent with a dynamic holoenzyme (Figure 4D). In activating conditions, we detected fewer homotypic crosslinks involving the regulatory domain than in basal conditions, either because CaM binding shields residues or because heterogeneous phosphorylation hindered peptide identification. It is notable that the regulatory segment was readily crosslinked in basal conditions (23 out of 136 crosslinks involved the regulatory segment), perhaps inconsistent with the canonical view that it is stably docked to the kinase domain in an autoinhibitory conformation.

Plotting interactions directly onto a model of the CaMKII holoenzyme (Myers *et al.*, 2017) gives insight into the phosphorylation pattern (Figure 5). Homotypic interactions are relatively uniform and spatially consistent across time points and basal or activated conditions (Figure 5B and C). In contrast, heterotypic interactions, modelled onto subunits in a pair of adjacent holoenzymes, are limited to sparse contacts. Without any calculation or docking, a “close” mode of interaction in which kinase domains from distinct holoenzymes are interdigitated satisfies the distance constraints of crosslinking much better (Figure S8) without introducing extensive clashes (~100 close atomic clashes mostly in flexible loops, compare with ~90k atoms per holoenzyme).

**Figure 5.**
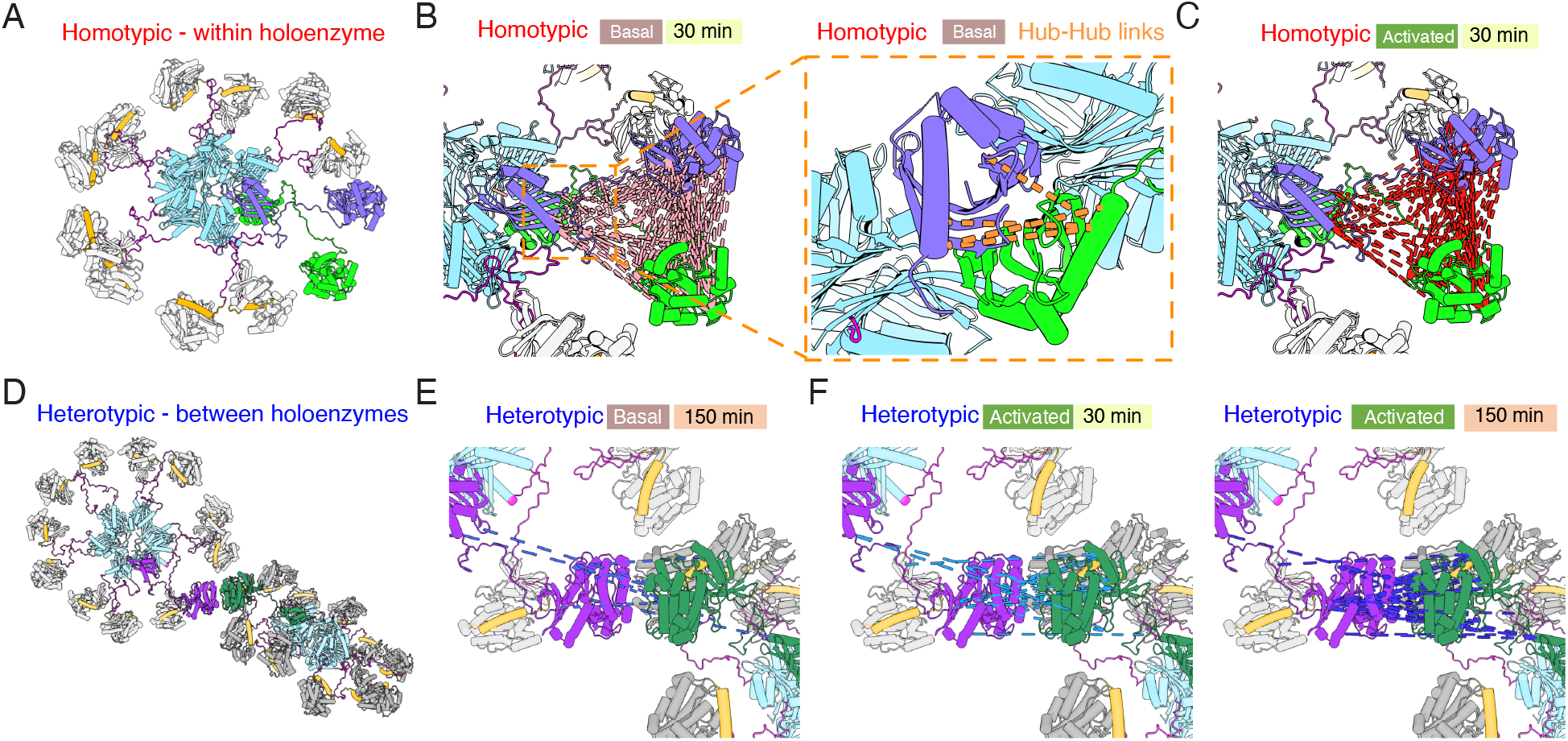
Mapping crosslinks onto holoenzyme structure. (A) Holoenzyme structure with two neighbouring subunits (green and purple) indicated (PDB: 5u6y). Hub domain is in light blue, regulatory segment (docked) in orange. (B) Basal crosslinks between homo-isotypes (30 minutes incubation, 136 crosslinks) plotted onto the holoenzyme as intersubunit interactions. Inset shows the 5 homo-isotopic crosslinks found for interactions between neighbouring hub domains. (C) Crosslinks between homo-isotypes in activated conditions (30 minutes, 85 crosslinks) showed a similar pattern to the basal condition. (D) Two holoenzymes arranged to allow kinase-kinase contacts to form (between green and purple subunits). (E) Sparse heterotypic crosslinks (8 in total) in the basal condition after 150 minutes. All heterotypic crosslinks involve the kinase domain. (F) In activating conditions, 15 heterotypic kinase-kinase interactions were detected after 30 minutes (out of 20 total). Over 150 minutes activation, 32 heterotypic kinase-kinase interactions were found (out of 43 total).

Across each of the four conditions, between 5 to 9 homotypic crosslinks were found between hub domains (Figure 5B). A frequently-observed crosslink involved lysines at 344 and 347 (Supplementary Tables 1,2,5,6). Steric considerations favour inter-subunit linkage in this case. On the other hand, heterotypic hub-hub interactions were not detected in any condition, contrary to expectations had subunit exchange between holoenzymes formed of 15N and 14N subunits occurred (Supplementary Tables 1-8). The dearth of heterotypic hub-hub crosslinks is in good agreement with absence of UV-induced hub domain crosslinking between subunits from different holoenzymes. In all activated samples, from both homo- and hetero-isotypes, we could detect phosphorylation on T286 (Figure S8, Supplementary Table 9 and 10), consistent with robust autophosphorylation within holoenzymes.

These types of interactions differ from information obtained by both the crystal structure of the linker-less inhibited CaMKII holoenzyme (Chao *et al.*, 2011), which is compact, as well as to the cryo-EM structure of the CaMKII holoenzyme with 30 residue linkers (Myers *et al.*, 2017), in which the interaction between the kinase and the hub domain is almost never observed. Upon activation, the interaction of the hub with the kinase domains is reduced to favour more kinase:kinase interactions presumably corresponding to trans-autophosphorylation both within and between holoenzymes (compare Supplementary Tables 5 & 6 with 7 & 8).

### Colocalization of CaMKII is activity-dependent

Much impetus for the subunit exchange theory comes from visualization of fluorescently labeled CaMKII molecules by TIRF microscopy (Stratton *et al.*, 2014) or fluorescence resonance energy transfer (FRET) experiments (Bhattacharyya *et al.*, 2016). Colocalization of dye-labelled subunits was interpreted to signify that CaMKII can undergo subunit exchange between activated and basal holoenzymes in order to spread its activity, and that this process was activity-dependent.

We performed TIRF experiments with purified CaMKII^WT^ and/or CaMKII^KD^, which were labelled at their N-termini *in vitro* using Sortase with Atto-488 or Atto-594 fluorophores (see scheme in Figure 6A). In order to monitor colocalization of CaMKII, we first incubated Atto-488-CaMKII^WT^ and Atto-594-CaMKII^WT^, under resting (basal) and activating (activated) conditions (Figure 6B). Subunits visualized in the microscope are bound to the coverslip, providing a “snapshot” of their association at the point of mounting. We used 55 nM CaMKII, a concentration at which dodecamers are around 98% of the population (see below). Limited colocalization between labeled CaMKII^WT^ subunits under resting conditions contrasted with around 50% colocalization of CaMKII^WT^ molecules after incubation in activating conditions (with Ca^2+^:CaM and ATP:Mg^2+^) (Figure 6B and C). This result is in good agreement with the previous literature (Stratton *et al.*, 2014). When we incubated CaMKII^WT^ with ATP:Mg^2+^ but without Ca^2+^:CaM, we could not detect colocalization (Figure 6C, Figure S10C). The same effect was observed when we stripped the Ca^2+^ ions by incubating Ca^2+^ /CaM with BAPTA, a robust Ca^2+^ -ion chelator (Collatz, Riidel and Brinkmeier, 1997), prior to adding it to the incubation reaction with CaMKII and ATP/Mg^2+^ (Figure 6C, Figure S10B). Furthermore, incubation of CaMKII^WT^ with Ca^2+^:CaM without ATP:Mg^2+^ again failed to drive CaMKII^WT^ colocalization and confirmed that both activating stimuli are necessary to drive colocalization of CaMKII (Figure 6C, Figure S10A).

**Figure 6.**
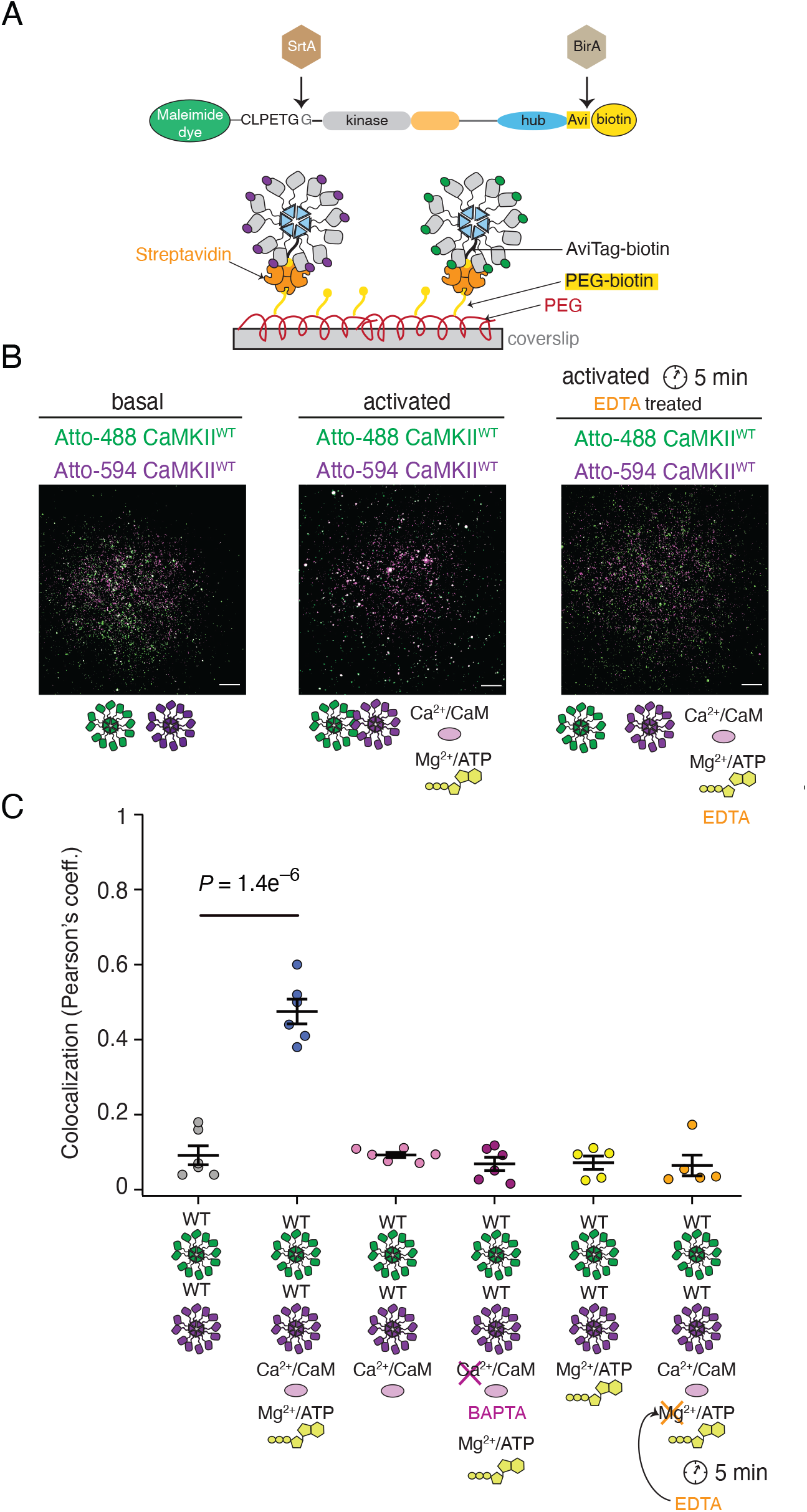
Reversible activity-dependent colocalization of CaMKII holoenzymes. (A) Schematic representation of CaMKII *in vitro* enzymatic labelling with maleimide dyes and biotin, and TIRF experimental set-up. (B) Representative TIRF images of unactivated (basal) CaMKII^WT^ sample (left), activated CaMKII^WT^ sample (middle), and CaMKII^WT^ sample first activated and then quenched with EDTA (right). (C) Summary of colocalization analysis (Pearson coefficient) for TIRF images for CaMKII under different conditions. Statistical significance was calculated using a multi-comparison test (Dunnett’s test) with α=0.05. Mean and standard deviation of the mean are indicated.

Finally, we looked for a way to initiate the kinase reaction, and putative subunit exchange, and to then quench the kinase reaction and trans-autophosphorylation. We envisaged that if the subunit exchange is indeed taking place during activation, at least partial colocalization of differently labelled CaMKII^WT^ molecules would be observed even after stopping the kinase reaction, thanks to subunit exchange before the reaction has been stopped. To test this possibility, we incubated differentially-labeled CaMKII^WT^ with activating stimuli (both Ca^2+^:CaM and ATP:Mg^2+^), allowed autophosphorylation to occur, and then added

EDTA to chelate Mg^2+^ and prevent further phospho-transfer during trans-autophosphorylation (Rudolf *et al.*, 2014). In this condition, we surprisingly found a total absence of subunit colocalization between differentially labeled CaMKII^WT^. This critical result indicates that colocalization is activity dependent but transient. It is likely not a consequence of the subunits being exchanged between holoenzymes. Rather, what we observe as a colocalization under TIRF comes from different holoenzymes coming into close proximity to trans-autophosphorylate each other in an inter-holoenzyme manner. When ATP was absent, the holoenzymes did not colocalize because they were not trans-autophosphorylating each other. Finally, when we incubated CaMKII^WT^ with CaMKII^KD^ we could still detect colocalization of CaMKII molecules under activating conditions, whereas colocalization of differentially labeled CaMKII^KD^ was completely absent in both conditions (Figure S11). The colocalization observed between CaMKII^WT^ and CaMKII^KD^ was less pronounced than for mixed WT subunits (Figure S11B), presumably because half of the sample in this case is inactive, and therefore the activity-driven colocalization occurs to a lesser extent.

### Higher order clusters of holoenzymes detected during activation

If subunits do not exchange, then holoenzymes must come in close proximity in order to trans-autophosphorylate each other. In order to test this hypothesis, we used mass photometry to compare the molecular weight (Mw) of CaMKII^WT^ and CaMKII^KD^ particles under basal and activating conditions. Because mass photometry is a surface-based microscopy method, the micromolar concentrations of CaMKII we used in other assays gave too many particles in the field of view for a valid measurement. The highest CaMKII concentration which we could measure was 400 nM. The high sensitivity of mass photometry gave the substantial advantage that we could work at concentrations of CaMKII particles as low as 10 nM. Figure 7A shows measured mass distribution of CaMKII^WT^ particles under resting conditions. The only peak which could be fitted with a Gaussian distribution corresponds to particles with Mw corresponding to a dodecamer (592 kDa). This peak was asymmetric on the high molecular weight side, indicating we worked at the high concentration limit of the experiment (Figure S12 A). Upon activation of CaMKII^WT^, we detected two additional sets of particles with molecular weights corresponding to dimer-tetramers and 24-30-mers, in addition to the main dodecameric peak (Figure 7B, Figure S12 B). The measured molecular weights in each peak correspond to CaMKII with bound calmodulin. Peak 0 has a Mw of 157 kDa, which corresponds to CaMKII dimer with two bound calmodulin. Peak 1 corresponds to particles with Mw of 721 kDa, close to CaMKII dodecamer with bound calmodulins. Finally, Peak 2 corresponds to particles with Mw of 1442 kDa, which is roughly CaMKII 24-mer with bound calmodulin. We presume that the 24-mer peak corresponds to 2 interacting holoenzymes. The abundance of CaMKII monomer in each peak was 0.7% for CaMKII dimer, 81.5% for CaMKII dodecamer, and 17.8% for CaMKII 24-mer (see “Methods” section for calculation of

**Figure 7.**
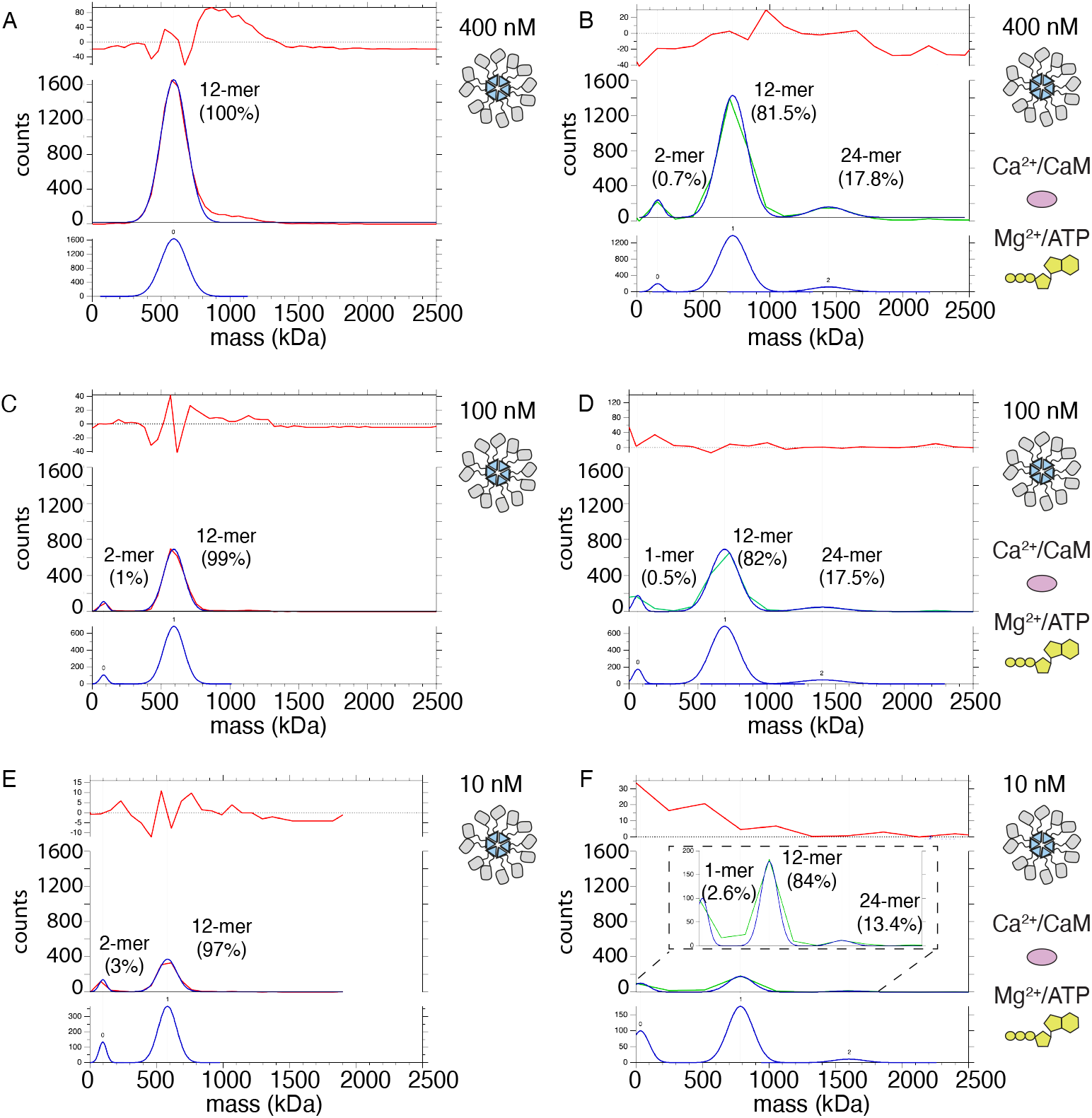
Mass photometry detects clusters of holoenzymes forming upon activation. (A) Mass distribution of 400nM CaMKII^WT^ under basal conditions (red curve). Blue curve is multi-Gaussian fit (shown separately on lower graph). Red curve in upper graph is the fit residual. (B) Mass distribution of 400nM CaMKII^WT^ under activating conditions (green curve), with multi-Gaussian fit in blue (peaks numbered in lower graph). (C) Mass distribution of 100nM CaMKII^WT^ under basal conditions (red curve). (D) Mass distribution of 100nM CaMKII^WT^ under activating conditions (green curve). (E) Mass distribution of 10nM CaMKII^WT^ under basal conditions (red curve). (F) Mass distribution of 10nM CaMKII^WT^ under activating conditions (green curve). Inset shows fit on expanded scale.

CaMKII monomer fractions in each peak). Under activating conditions, the majority of CaMKII forms dodecamers, implying that most of trans-autophosphorylation still occurs intra-holoenzyme. However, around 18% of CaMKII forms 24-mers, indicating that the spread of kinase activity might occur between two holoenzymes (inter-holoenzyme). Mass photometry detected less than 1% of CaMKII subunit dimers at 400 nM concentration upon activation, which could be interpreted as either subunit dimers leaving holoenzymes and exchanging with other holoenzymes, or as a general destabilization of a sub-set of holoenzymes brought about by activation. The distribution of Mw peaks of CaMKII at 100 nM concentration showed a monomer-dimer peak (Peak 0 = 81.5 kDa) already appears in resting conditions (Fig. 7C; Figure S12 C), albeit at low intensity (representing around 1% of the sample), indicating that lowering CaMKII concentration destabilizes holoenzyme integrity. The main peak still corresponded to dodecameric CaMKII (Peak 1 Mw = 591 kDa). Incubation with activating stimuli at 100 nM concentration generated a similar peak distribution as during activation at 400 nM CaMKII. The first peak corresponded to a monomer population (Peak 0 Mw = 62 kDa) participating with around 0.5% of CaMKII monomers, the main peak is still dodecameric (Peak 1 Mw = 693kDa) with about 82% of CaMKII monomers participating in this peak, and finally the 24-mer peak was detected (Peak 2 Mw = 1404 kDa) consisting of 17.5% of all CaMKII monomers. (Figure 7D, Figure S12D). Lowering CaMKII concentration to 10 nM further destabilized the holoenzyme, as can be seen in the appearance of higher percentage (3%) of particles corresponding to Mw of CaMKII dimer (Peak 0 Mw = 95 kDa), but still the majority of particles formed dodecamers (97%, Peak 1 Mw = 581 kDa) (Figure 7E). Remarkably, activation of CaMKII at 10 nM concentration still led to formation of 24-mers (Peak 2 Mw = 1600 kDa with 13.4%), dodecameric fraction at 84% (Peak 1 Mw = 784 kDa), and monomeric at 2.6% (Peak 0 Mw = 33.2kDa). The peak distribution of activated samples is similar over the range of concentrations, further emphasizing the stability of CaMKII holoenzyme and propensity to form higher order oligomers during activation. CaMKIIKD particle distributions at 400 nM concentration were similar to the WT protein under basal condition, but the maximum Mw detected was corresponding to decameric holoenzymes (Figure S13A). Activation of KD protein gave no higher-order peak but generated a small peak of tetrameric CaMKII (12%, Fig. S13B). The absence of a higher order oligomeric peak from the CaMKIIKD sample in activated conditions further emphasizes that transient CaMKII clustering is driven by trans-autophosphorylation between holoenzymes.

## Discussion

The importance of CaMKII activity, and its idiosyncratic subunit arrangement in dodecameric holoenzymes, have stimulated great interest in its mechanisms of action. Particularly beguiling is the idea that CaMKII can convert brief Ca^2+^ signals corresponding to decisions, calculations and comparisons in neurons into sustained phosphorylation of targets in order to store information in neural circuits (Lisman, Schulman and Cline, 2002; Coultrap and Bayer, 2012; Tao *et al.*, 2021). The prevailing model for the requisite spatial and temporal spread of CaMKII autonomous activity argues that CaMKII must exchange subunits between activated and naïve (unactivated) holoenzymes (Stratton *et al.*, 2014; Bhattacharyya *et al.*, 2016), principally because previous work suggests autophosphorylation between holoenzymes does not occur (Miller, S. G. & Kennedy, 1986; Hanson *et al.*, 1994; Rich and Schulman, 1998). The data we present here, collected over 5 separate, complementary techniques, convincingly demonstrate that subunit exchange, if it does occur, is a rare event. We also provide support for a new model of efficient spread of kinase activity, through inter-holoenzyme trans-autophosphorylation. In this model, CaMKII subunits remain in their respective stable holoenzymes, but phosphorylate neighboring holoenzymes, exploiting the long, flexible linkers which connect the kinase domains to the hub. Contacts between kinase domains are driven by activity, presumably corresponding to multivalent, synergistic autophosphorylation events. The high local concentration of CaMKII in dendrites would be expected to facilitate this interaction between holoenzymes.

Why did previous studies fail to detect inter-holoenzyme phosphorylation? We think that three factors contributed: low concentrations of CaMKII substrate (kinase-dead CaMKII), short incubation times (30-60s) and low temperature (4°C) in previous experiments (Hanson *et al.*, 1994; Rich and Schulman, 1998). We showed that CaMKII^WT^ can phosphorylate CaMKII^KD^ subunits from distinct holoenzymes *in vitro*, at 37°C, and also in mammalian cell extracts. We used 4 μM kinase-dead CaMKII, in order to approach the physiological CaMKII concentration, which is estimated to be 20-100 μM in dendritic spines (Otmakhov and Lisman, 2012). We further confirmed that CaMKII^KD^ holoenzymes can associate with CaMKII^WT^ holoenzymes in a single molecule TIRF assay.

Evidence indicating that CaMKII spreads its kinase activity via subunit exchange between activated and unactivated holoenzymes came from single molecule fluorescence experiments, which monitored colocalization of differently-labelled CaMKII under TIRF or by FRET (Stratton *et al.*, 2014; Bhattacharyya *et al.*, 2016). These studies identified activation-dependent colocalization of CaMKII subunits. We observed the same, but critically, if we triggered activation normally but then quenched it, before TIRF observation, the colocalization was entirely lost. At least two possibilities exist for obtaining this result. However improbable, subunits that exchanged during the activation period might have re-equilibrated to a low level of colocalization within their respective holoenzymes upon inhibition of the kinase reaction. Such selective reassembly seems far-fetched because our CaMKII^WT^ subunits differed only by the identity of the fluorophore at the N-terminus. More likely, subunits did not exchange at all during the kinase reaction, but instead what we (and others, (Stratton *et al.*, 2014)) have observed is the transient colocalization of holoenzymes during inter-holoenzyme phosphorylation.

The colocalization of kinase-dead and WT CaMKII subunits was weaker, which is expected if association requires kinase activity, because in this case only half of the sample was competent to perform phosphorylation. Importantly, this interpretation presents little conflict with existing data, which failed to detect colocalization of CaMKII^WT^ and CaMKII^WT^ incubated with the inhibitor bosutinib, used to mimic kinase-dead CaMKII (Stratton *et al.*, 2014). According to our model, the absence of colocalization in CaMKII^WT^ inhibited by bosutinib is in the nature of the inhibitor binding, which blocks the ATP-binding site (Pellicena and Schulman, 2014) and stabilizes compact inactive conformation of CaMKII (Chao *et al.*, 2011). In such a conformation, CaMKII is not able to initiate trans-autophosphorylation, which manifests as a lack of colocalization, like in our TIRF experiments when ATP was omitted.

We directly probed subunit mixing with a photoactive crosslinker in the hub domain, but could not detect any mixing, nor did the restraint have any effect on kinase activity. The genetically-encoded BzF crosslinker is monofunctional, and can capture unlabeled subunits at interfaces within ~5-10 Å (Wittelsberger, Mierke and Rosenblatt, 2008). Even though crosslinking of the hub domain by BzF was not saturated, we observed bands of apparent high molecular weight above 0.3 MDa, yet the kinase activity (between untreated and crosslinked enzymes) was identical. In this context, we note that if subunit exchange were a prominent mechanism for spreading activity, then small molecules that stabilized the hub domain should reduce activity. However, binding of the putative CaMKII inhibitor GHB, which was shown to stabilize the hub domain, left substrate phosphorylation and trans-autophosphorylation untouched (Leurs *et al.*, 2021). We could detect phosphorylation of CaMKII^KD^ by CaMKII^F394BzF^, but we could not detect kinase-dead protein in higher order oligomeric bands on western blot, which were exclusively from CaMKII^F394BzF^. This result matches well with the much more promiscuous crosslinking of lysines by the bifunctional DSS crosslinker, which showed less crosslinking between subunit isotypes from different preparations (N14 vs N15), but abundant crosslinking within isotypes (N14:N14 and N15:N15).

How do holoenzymes interact during IHP? Crosslinks between heterotypic peptides detected by mass spectrometry were much more prevalent in activating conditions, reporting activity-dependent holoenzyme contacts with a spacing of 10Å or less. We detected direct capture of substrate-active site peptide interactions (including phosphorylation of T286, Figure S8 and Supplementary Tables 9 & 10). Almost all peptide pairs were detected with an intensity ratio below one, meaning links between homo-isotypes (likely within holoenzymes) were more frequent than links between hetero-isotypes (between holoenzymes). This observation could in principle indicate that heterotypic holoenzymes (composed of mixed N14 and N15 subunits) were present, yet in the minority. However, the profile of crosslinks, being absent in all but the kinase domains for heterotypic links, speaks against the idea of subunit exchange. In particular, almost all crosslinks cover parts of the substrate binding cleft and the catalytic machinery. Heterotypic crosslinks were not detected in the hub domain. The mean ratio for kinase-kinase crosslinks was 25 ± 1% (32 crosslinks in activated conditions, 150 min incubation). If we assume that all heterotypic links come from inter-holoenzyme contacts, IHP might occur at about ¼ the rate of intra-holoenzyme autophosphorylation (in other words, with surprising speed). The flexible linkers might allow inter-holoenzyme contacts involving multiple kinase domains to act in a “cooperative manner”. We illustrate this possibility in our speculative “close mode” involving interdigitated holoenzymes, which reduces crosslink distances substantially (Figure S9). These observations further cast doubt on the rule that inter-holoenzyme autophosphorylation cannot occur. If a CaMKII dodecamer with mobile kinase domains were to abundantly phosphorylate its targets and auto-phosphorylate its own domains, but never auto-phosphorylate a neighbour despite being one of the most abundant cytosolic proteins, it should require as yet unknown additional factors which have been absent in all *in vitro* experiments to date.

Attempts to analyze the number of CaMKII molecules colocalized under TIRF by photobleaching of well-labelled CaMKII (Theile *et al.*, 2013; Elsner *et al.*, 2019) proved to be unreliable. The large number of subunits present means that shot noise overwhelms the stepwise fluorescence loss during single subunit bleaching. Instead, we used mass photometry to obtain the distribution of molecular weights of CaMKII particles in basal and activating conditions. We detected activation-dependent appearance of particles corresponding to pairs of holoenzymes, supporting the idea that phosphorylation can occur in an inter-holoenzyme fashion (as well as the established intra-holoenzyme reaction). Similarly, self-association of CaMKII has previously been described *in vitro* (Hudmon *et al.*, 1996, 2001), as well as in cells (Hudmon *et al.*, 2005), where it was linked to ischemia-like conditions of lower pH. CaMKII clustering has also been detected on negative-stain EM grids (Buonarati *et al.*, 2021), albeit no link to activation was asserted. Although peptides derived from the regulatory segment of CaMKII can break isolated hubs to lower molecular weight subunits in electrospray mass-spectrometry, the effect was absent in solution (Karandur *et al.*, 2020). Consistent with this, we observed only a very minor fraction of monomer-dimer particles in activating conditions. Keeping in mind that the affinity for holoenzyme assembly is in low nM range (Torres-Ocampo *et al.*, 2020), an observation that we could replicate (Figure 7), and that in the packed environment in dendritic spines, the concentration of CaMKII is in μM range (Otmakhov and Lisman, 2012), it is likely that CaMKII forms clusters of holoenzymes during activation, where more than 2 holoenzymes participate, but we did not detect a long tail in the mass distribution corresponding to larger supercomplexes.

What are the advantages or disadvantages of inter-holoenzyme phosphorylation? First of all, we should consider the dodecameric form of CaMKII. Under persistent phosphatase attack, dodecamers can rescue their fully-phosphorylated state and autonomous activity, through intra-holoenzyme trans-autophosphorylation. In contrast, isolated CaMKII subunits in lower order forms (which should exist in the subunit exchange model) would be vulnerable to terminal dephosphorylation, destroying the signal. Gains in reliability may be offset against speed or scope. In principle, subunit exchange in tandem with rapid intra-holoenzyme autophosphorylation could grow CaMKII activity exponentially, albeit with a slower initial rate, because of the time to break and reform dodecamers. Inter-holoenzyme phosphorylation should proceed with a faster initial rate but be less prone to saturate local stocks of holoenzymes. Indeed, measurements in dendrites suggest a steady summation of CaMKII activity (Chang *et al.*, 2019; Tao *et al.*, 2021), not a runaway exponential wave. Capture of peptides including phosphorylated T286 were only about 8x less abundant in heterotypes (see Supplementary Tables (& 10 and Figure S9). The crosslinks obtained from mass-spectrometry suggest that CaMKII kinase domains are highly dynamic. Therefore, the loss of reaction rate from a lack of proximity might be compensated by the larger degree of freedom for inter-holoenzyme kinase-kinase domain interactions. Cooperative phosphorylation, whereby contacts made during inter-holoenzyme phosphorylation may be augmented by additional interactions, is naturally facilitated by the dodecameric ring architecture. Fixed dodecameric holoenzymes should have a larger fraction of their time to perform downstream signaling, unlike holoenzymes that must disassemble to spread activity. In our model of inter-holoenzyme phosphorylation, active dodecamers are a finished product, whose activity depends only on the rate constants of phosphorylation and stochastic attack of phosphatases. Notably, this model predicts that there should be some dependence of phosphorylation spread on linker length. Isoforms with short linkers should propagate activity more slowly due to steric restrictions.

## Materials and methods

### Expression constructs

Human *wild-type* CaMKII lacking the first 5 residues, and with a 30-residue linker (CaMKII^WT^) was codon optimized for expression in *E. coli*, and gene synthesized (GenScript) in pET28b+ plasmid, with flanking restriction sites NcoI and HindIII. Mutations K42R, D135N, F394TAG, and H418TAG were introduced by site-directed mutagenesis. AviTag was added to the C-terminus of CaMKII^WT^ and CaMKII^K42RD135N^ by splice overlap extension PCR. In order to express CaMKII in mammalian cells, human CaMKII^WT^, and human CaMKII^K42RD135N^ with a C-terminal AviTag were cloned into pEGFP-N1 (Clontech) vector, with an IRES sequence inserted between CaMKII gene and EGFP. Rat CaMKII^WT^ with an N-terminal EGFP pCAG-mEGFP-CaMKIIa was a gift from Ryohei Yasuda (Addgene plasmid # 127389; http://n2t.net/addgene:127389; RRID:Addgene_127389; Chang *et al.*, 2018). The sortase A pentamutant (eSrtA) in pET29 was a gift from David Liu (Addgene plasmid # 75144; http://n2t.net/addgene:75144; RRID:Addgene_75144, Chen et al., 2011) and pET21a-BirA (Biotin Ligase) was a gift from Alice Ting (Addgene plasmid # 20857; http://n2t.net/addgene:20857; RRID:Addgene_20857, Howarth et al., 2005).

The vector encoding Lambda Phosphatase (LPP) was a gift from John Chodera & Nicholas Levinson & Markus Seeliger (Addgene plasmid # 79748; http://n2t.net/addgene:79748; RRID:Addgene_79748; Albanese et al., 2018). His-tagged Syntide 2 (PLARTLSVAGLPGKK) - GST was cloned into pETM11 vector.

### Protein expression

All CaMKII constructs (Kanamycin resistance) were co-expressed with Lambda protein phosphatase (LPP) (Spectinomycin resistance) (Chao *et al.*, 2011) in *E. coli* BL21 cells, in LB medium, supplemented with 1 mM MnCl_2_, which is a co-factor of LPP. Cultures were grown at 37°C, until OD_600_ = 0.8-1, and then protein expression was induced by adding 0.4 mM IPTG. Expression was continued over-night at 20°C.

For expression of ^15^N-incorporated CaMKII, BL21 *E. coli* co-transformed with CaMKII^WT^ and LPP were grown at 37°C in minimal media (2mM EDTA, 3.5 mM FeSO_4_, 0.4 mM ZnCl_2_, 0.06 mM CuSO_4_, 1mM MgSO_4_, 0.3mM CaCl_2_, 50 mM Na_2_HPO_4_, 25 mM KH_2_PO_4_, 10mM NaCl, 0.5% glucose, 1.5 μg/mL Thiamine, 0.15 μg/mL Biotin) supplemented with 0.75 mg/mL of ^15^NH_4_Cl as a source of ^15^N isotope. Expression was induced by adding 0.5 mM IPTG at OD_600_ = 0.8-1, and the expression was continued over night at 20°C/200rpm.

Expression of BzF constructs was enabled by co-transformation of BL21 cells with individual plasmids containing a CaMKII amber (TAG) mutant, LPP, and orthogonal aminoacyl synthetase (aaRS) and tRNA from *M. jannaschii* (Young *et al.*, 2010), which was a kind gift from Prof. Dr. Thomas Söllner. aaRS and tRNA were in a pEVOL vector under control of the araBAD promoter. In order for CaMKII to incorporate BzF at a desired position, the growth media was supplemented with 1 mM BzF at OD_600_ = 0.6 at 37°C. After 30 minutes, the pEVOL vector expression was induced by addition of 15 mM arabinose. Finally, at OD_600_ = 1.2 – 1.5, CaMKII and LPP protein expressions were induced by addition of 0.4 mM IPTG, and the media was supplemented with 1 mM MnCl2. The cells continued to grow over night at 20°C, 220 rpm.

### Protein purification

Cell pellets were lysed in lysis buffer (50 mM Tris pH 8, 300 mM NaCl, 20 mM imidazole, 1 mM TCEP) supplemented with 0.02 mg/mL DNaseI, 0.5 mg/mL lysozyme and 1 mM PMSF. The lysates were additionally passed through a cell disruptor two times, before the final centrifugation step (16k rpm, 4°C). All CaMKII constructs have an N-terminal His_6_ tag, which was used for affinity purification with 5 mL NiNTA column (GE Healthcare), which was previously equilibrated in Ni_A_ buffer (50 mM Tris pH8, 300 mM NaCl, 20 mM imidazole, 1 mM TCEP). The column was then extensively washed (100 mL of Ni_A_ buffer, 50 mL of Ni_A_ buffer with 50mM imidazole, 50 mL of Ni_A_ buffer with 80 mM imidazole), and finally eluted with a gradient elution from 80 mM imidazole to 1M imidazole (50 mM Tris pH8, 300 mM NaCl, 1 mM TCEP, 1M imidazole) over 10 column volumes. Peak fractions were then pooled, imidazole concentration lowered to less than 100 mM with Ni_A_ and the sample was incubated with 0.01mg/mL final TEV (made-in-house) in order to cleave off the His_6_ tag. On the following day, the cleaved protein was concentrated to 500 μL, using 50 kDa cut off concentrator (4 mL Amicon R Ultra), and injected to Superose 6 10/300 column (GE Healthcare) previously equilibrated in size exclusion chromatography (SEC) buffer (25 mM Tris pH 8, 250 mM NaCl, 1% glycerol, 1 mM TCEP). Elution profile comprised of 2 peaks, one of which corresponded to molecular weight of dodecameric CaMKII (around 650 kDa) and the other one to 40 kDa CaMKII fragment, which is usually present during CaMKII purification from *E. coli*. The peak corresponding to dodecameric CaMKII was pooled and further concentrated to approximately 2 mg/mL. We obtained around 0.5-1 mg of protein per 2L of *E. coli* pellet.

His tagged Syntide 2 – GST was purified from BL2-RIL cell pellets, using a two-step purification process. Frist, the cell pellet was resuspended in lysis buffer (50 mM Tris-HCl pH 7.5, 150 mM NaCl, 20 mM imidazole, 1 mM DTT) containing protease inhibitors (cOmplete, Roche) and further lysed by sonication. Upon centrifugation (16k rpm, 4°), cleared cell lysate was loaded on a HisTrap Crude column (Cytiva) and eluted with a linear gradient of elution buffer (20 mM Tris-HCl pH 7.5, 300 mM NaCl, 500 mM imidazole, 1 mM DTT). Protein containing fractions were pooled, incubated with TEV protease, and dialysed against lysis buffer o/n. Digested sample was re-run on a HisTrap Crude column. The flow through was collected, concentrated and run on a High Load Superdex 75 26/60 size exclusion column (Cytiva), equilibrated in SEC buffer (20 mM PIPES pH 7.5, 50 mM NaCl). Protein containing fractions were pooled, concentrated to >20mg/mL and flash frozen in liquid nitrogen.

Calmodulin from human was codon-optimized for expression in *E.coli* and purified according to published protocols (Putkey and Waxham, 1996).

BirA was expressed and purified according to a protocol from (Fairhead and Howarth, 2015).

SrtA was expressed and purified according to (Popp, 2015).

### Kinase assay with purified proteins

Kinase assays for western blot detection were performed at 37°C, for 1h, with samples taken at 1-, 2-, 5-, 10-, 15-, 30- and 60-minute time points. Each time point was done in triplicate. Each reaction was performed in 10 μL volume with final concentrations as follows: 10 nM CaMKII^WT^, 4 μM CaMKII^K42RD135N^-AviTag, 2 μM CaM (or 100 nM in “low CaM condition”), 2 mM CaCl_2_, 10 mM MgCl_2_, 125 mM NaCl, 25 mM Tris pH 8, 2mM TCEP, 100 μM ATP. First, 5 μL of 2x master mix containing 20 nM CaMKII^WT^, 8 μM CaMKII^K42RD135N^-AviTag and 4 μM CaM, in SEC buffer was distributed in PCR tubes. Then 2 μL of 5x reaction buffer (10 mM CaCl_2_ and 50 mM MgCl_2_) was added to each tube. Finally, 3 μL of 350 μM ATP was added to each tube, using a multi-channel pipette, starting from the longest time point (60 min). In order to quench the reactions, 10 μL of 4x standard sample loading buffer was added using a multichannel pipette, starting with the shortest time point (1 min), followed by heat denaturation (2 min at 95°C). Quenched reactions were kept at 4°C before a western blot was run on the following day. 5 μL of each quenched reaction was loaded on 12% SDS-PAGE gel, giving total of 1-2 ng of CaMKII^WT^ and around 500 ng of CaMKII^K42RD135N^-AviTag per lane.

The kinase assay for radioactivity detection was performed like described above, with several modifications. Briefly, the final reaction volume was 20 μL with final concentrations as follows: 15 nM CaMKII^WT^, 4.4 μM CaMKII^K42RD135N^-AviTag, 2 μM CaM (or 100 nM in “low CaM” condition), 2 mM CaCl_2_, 10 mM MgCl_2_, 0.1% BSA, 125 mM NaCl, 25 mM Tris pH 8, 2mM TCEP, 100 μM ATP. First 11 μL of master mix was distributed in PCR tubes, followed by the addition of 4 μL of 5x reaction buffer (250 mM PIPES pH 7.2, 10 mM CaCl_2_ and 50 mM MgCl_2_, 0.5% BSA), and finally 5 μL of ATP. ATP was made by adding 2 μL of P^32^ radioactive ATP to 400 μM stock of regular P^31^ ATP. Reactions were then sampled like described above, and stopped by adding 10 μL of 4x standard sample loading buffer. 10 μL of each sample was loaded on 12% SDS-PAGE gel, giving total of 5 ng of CaMKII^WT^ and around 1.5 μg of CaMKII^K42RD135N^-AviTag per lane. To probe substrate phosphorylation of CaMKII^F394BzF^, instead of CaMKII^K42RD135N^-AviTag, we used 40 μM of GST-Syntide-2, and sampled the kinase reactions as described above.

### HEK cell transfection and lysis

1.5 million cells were seeded per one well of a 6-well dish, and on the following day the cells were transfected using Polyethylenimine (PEI). For expression of single constructs, 1 ug of plasmid DNA was used with 3 μL of PEI, and mixed in 250 μL of Opti-MEM, vortexed briefly, left for 20-30 minutes for complexes to form and finally added to respective wells in a drop-by-drop manner. For co-expression, 1 ug of each construct was mixed in the same microcentrifuge tube with 6 μL of PEI in 250 μL of Opti-MEM. The medium was exchanged after 5h to fresh 6% FBS DMEM medium, and cells were left expressing the constructs for 48h before harvesting and cell lysis. In order to harvest the cells, the medium was first aspirated, and the cells were resuspended in 1mL of ice cold DPBS per one well, and placed in ice cold microcentrifuge tubes, on ice, until a centrifugation step. After centrifugation at 2000 rpm for 10 min, the cells were snap frozen in liquid nitrogen and stored at −80°C until further use. Cell lysis was performed in 100 μL of lysis buffer (per well): 50 mM Tris pH 8, 150 mM NaCl, 1% Triton X 100, 1 mM TCEP, supplemented with protease inhibitors (1 μM Pepstatin A, 10 μM Leupeptin, 0.3 μM Aprotinin, 1 mM PMSF), 5 mM NaF, 0.02 mg/mL DNaseI, and 5 mM MgSO_4_. After resuspension, the cells were left on ice for 20 min, with brief vortexing every 5 min. The lysates were then cleared by high-speed centrifugation for 15 min at 4°C in a table-top centrifuge. Supernatants, containing CaMKII, were taken for the kinase assay.

### Kinase assay using HEK cell lysates with overexpressed CaMKII

20 μL of each cleared cell lysate (CCL) was combined with 22 μL of lysis buffer and 12 μL of 4x SDS-PAGE loading dye and heated at 95°C for 2 min. 20 μL of this mixture was then loaded per well of 3-8% SDS-PAGE gel, and further subjected to western blotting in order to assess activity of overexpressed CaMKII.

Kinase assay was performed by incubating 20 μL of CCL with either 20 μL of lysis buffer or 20 μL of another CCL, under activating conditions with extra added 0.5 μM purified CaM and 500 μM ATP, at different temperatures (4°C and 37°C) and different incubation times (1 min or 10 min). Additionally, 20 μL of CCL containing overexpressed CaMKII^WT^ was incubated with 8 μM purified CaMKII^K42RD135N^-AviTag, under activating conditions. The reaction was quenched by adding 4x SDS-PAGE loading dye, and heating at 95°C for 2min.

### Western blot detection

Samples were run on a 3-8% pre-casted gels (Thermo Scientific), in 1xTA buffer (Thermo Scientific) on ice for 1h and 10 min at 180V. Transfer to pre-activated PVDF membrane (Millipore) was performed for 2h at 50V at 8°C, using Criterion Blotter (BioRad) in Towbin’s transfer buffer with 20% MeOH. Membranes were blocked in Intercept Blocking Buffer (IBB) (LiCor) for 1 h shaking at room temperature. Incubation with primary antibodies was done over-night at 8°C. Membranes were incubated at the same time with a mixture of primary antibodies. Rabbit anti-pT286 primary antibody (CellSignalling Technologies, product #12716) was diluted 1:1000, while mouse anti-AviTag antibody (antibodiesonline, clone number 1D11D10) was diluted 1:400 in IBB, supplemented with 0.01% tween and 0.02% Na_2_N_3_. On the following day, membranes were washed 3×10 min in TBS-T buffer, and incubated with fluorescently labelled secondary antibodies (IRDye 800CW Goat-anti Rabbit IgG and IRDye 680LT Goat-anti Mouse IgG from LiCor) for 1h at room temperature, shaking. Again, a mixture of two secondary antibodies was diluted 1:10 000 in IBB buffer, supplemented with 0.01% tween and 0.01% SDS. The two secondary antibodies, goat-anti rabbit and goat-anti mouse, are excited at 800nm and 700nm, respectively. After secondary anti-body incubation step, membranes were washed 3×10 min in TBS-T and imaged using LiCor imager. Blots were quantified using ImageJ/FIJI, and measured densitometry was plotted using IgorPro9. The data was fitted using Langmuir function.

### Radioactivity detection

Radioactive kinase reactions were run on 12% SDS-PAGE gels, followed by gel fixation and drying. The radioactivity was visualized using a photostimulable phosphor plate. Gel images were quantified using ImageJ/FIJI. Measured densitometry was plotted using IgorPro9, and fitted with Langmuir function.

### UV-induced crosslinking

In order to induce crosslinking of BzF to neighboring residues in CaMKII^BzF^ mutants, we used a UV lamp which can emit UV light of 365 nm wavelength (Opsytec Dr. Gröbel). The crosslinking was performed in the cold room, using a cold steal metal plate (made-in-house), to avoid excessive heating of the sample. CaMKII concentration during UV illumination was 8 μM in final volume of 20 - 40 μL. Each sample was treated 5 times with fifteen 1 s pulses, with 1s break between pulses. After the illumination, samples were mixed with appropriate amount of SDS-PAGE loading dye, heated at 95°C for 2 min and loaded on 3-8% SDS-PAGE gel. Around 1-2 ug of total protein was loaded per lane. Gels were run on ice for 1h 10 min/180V, and later stained with Coomassie dye or subjected to western blotting.

### Protein labelling for TIRF imaging

Biotinylation of CaMKII-AviTag was performed for 2h on ice in 100 μL final reaction volume, following an established protocol (Stratton *et al.*, 2014) with small modifications. The following final concentrations of components were used: 50 μM CaMKII-AviTag, 2.5 μM BirA, 2 mM ATP, 5 mM MgCl_2_, and 150 μM biotin. After biotinylation, the protein was subjected to fluorescent labelling using SrtA, following already published protocols (Theile *et al.*, 2013).

On the previous day, SrtA recognition peptide (CLPETGG) was fluorescently labelled by incubation with maleimide dyes (ATTO-Tech) in 1.3x excess dye over peptide (4.8 mM peptide and 6.24 mM ATTO-488; 4.22 mM peptide and 5.25 mM ATTO-594) in the dark at room temperature. Peptide labelling reaction was quenched on the following day by adding 1 μL of concentrated BME, and 50 μL of labelling buffer (100 mM Tris, pH 7.5, 150 mM KCl) to the final reaction volume of 120 μL. The sortase labelling reaction was incubated for 3h at room temperature in 400 μL final volume. The reaction contained 100 μL of biotinylation reaction, 12.5 μM SrtA, 40 μL of 10XSrtA buffer (500 mM Tris pH 7.5, 1.5M NaCl, 100 mM CaCl2), 120 μL of labelled peptide, 120 μL of SEC buffer. In order to get rid of BirA and SrtA and excess labelled peptide, the sample was first incubated with cobalt beads (GE Healthcare) to remove BirA and SrtA which both have an N-terminal His-tag. The protein-containing supernatant was then applied to PD-10 Sephadex desalting column (GE Healthcare) to remove access of peptide/dye, and eluted using SEC buffer. Elution fractions were analysed for the presence of labelled CaMKII by running SDS-PAGE and imaging the gel on a BioRad imager, using respective wavelengths. Fractions containing labelled CaMKII were pooled and concentrated using 50 kDa cut-off concentrator tube (4 mL Amicon R Ultra) to approximately 0.5 mg/mL in 150 – 200 μL.

### Preparation of cover slips for TIRF imaging

To reduce background signal and sources of autofluorescence, the coverslips (High Precision 1.5H, 25 mm diameter from Marienfeld GmbH) were cleaned thoroughly, following an established protocol (Paul and Myong, 2022). Briefly, the coverslips were first scrubbed with 5 % alconox using gloved hands. Then, they were sonicated in a sonicator bath (Elmasonic S 30 H Sonicator from Elma) for 20-30 min in MilliQ water. Next, they were etched with 1 M KOH for 30-45 min. Then, they were rinsed with water and dipped in 100 % EtOH before being passed through a butane flame 4-5 times and dried completely. The next step consisted in coating with aminosilane ((3-Aminopropyl)triethoxysilane from Sigma Aldrich) for 30-45 min. For this, 1 mL of aminosilane was diluted with 11 mL of acetone before being put on the coverslips. After rinsing with water, the coverslips were then coated with a mixture of 98% mPEG (5kDa) and 2% biotin-PEG (Silane-PEG-biotin 5kDa from abbexa) in 0.1 M sodium bicarbonate (pH 8.5). Two coverslips were incubated on top of each other, with the PEG-solution in between, overnight. The next day, the coverslips could be separated, rinsed, dried completely and then stored at −20 °C until use. Just before use, the coverslips were taken out of the freezer, left to equilibrate at room temperature for 30 min, and then coated with NeutrAvidin, by incubating each slide with 2mL of 0.1 mg/mL Neutravidin dissolved in PBS (Stratton *et al.*, 2014). The coverslips were then washed 10 x 1mL of PBS prior to sample addition and imaging.

### Preparation of samples for TIRF imaging

2 μL of 8 μM labelled proteins were added to 300 μL of imaging buffer (25 mM Tris pH 8, 150 mM KCl, 1mM TCEP). If needed for the experiment, 150 nM of CaM, 1 mM ATP and 2 mM MgCl_2_ were added and incubated for 2 min. The sample was then added on the coverslip, incubated for 1 min and then rinsed 3 times with imaging buffer. Coverslips were kept in 1 mL of imaging buffer during imaging. In order to chelate Ca^2+^ from CaM, we first incubated Ca^2+^:CaM (where Ca^2+^ is at 5 mM) with 15 mM BAPTA for 5 min at room temperature, and then added this mixture to the rest of the reaction. Final concentration of BAPTA was 0.25mM, prior to putting the sample on the cover slip and washing.

For chelation of Mg^2+^ ions, we first incubated the kinase reaction like usual (see above), and after 5 min. we added 10 mM EDTA (to chelate 2 mM Mg^2+^), incubated for 5 min at room temperature, and then added the sample to the coverslip, washed 3x with imaging buffer and imaged.

### TIRF imaging

Imaging was performed using a Nikon Ti2 Eclipse microscope with a temperature-controlled stage (set to 37°C). Excitation light was provided by alternating 488 nm (diode) and 561 nm (DPSS) lasers housed in an Omicron Lightbox. The laser light was filtered using a Quad band notch filter (400-410, 488, 561, 631-640) and reflected through the 60x Apo TIRF 1.49 NA objective by a quad band dichroic filter (zt405/488/561/640 rpc; from set F73-400). Coverslips were illuminated in TIRF mode and the angle was adjusted in software (Nikon NIS elements) as was the motorised stage and autofocus system. Emission light was filtered with 525/40 (488 nm excitation) or 600/52 filters (561 nm excitation, both Semrock). Images were recorded with a Prime95B 22mm sCMOS camera.

### TIRF image analysis

Colocalization analysis was done with ImageJ/ FIJI (Schindelin et al., 2012) and the colocalization Plug-In “JaCoP” (Bolte and Cordelières, 2006). To remove background, we set the threshold to the top 1% of signal and used it to create an image mask for each channel. These masks were then applied to the original image, which then provided with thresholded images used for colocalization analysis. In order to quantify the amount of colocalization that can be observed, the thresholded images were then analysed using “JaCoP” Plug-In in FIJI and the Pearson’s correlation coefficient was calculated. The Pearson’s correlation coefficient is a measure of linear correlation between two sets of data, the value ranging from −1 for negative correlation to 1 for positive correlation. For each condition, five to six data sets were analyzed, from recordings made on multiple days. The significance was calculated using a multi-comparison test (Dunnett’s test) with α = 0.05 in IgorPro9.

### Mass Photometry

Mass Photometry was performed on Refeyn Two^MP^ mass photometer. The unactivated samples of CaMKII were prepared by diluting 8 μM CaMKII^WT^ in TIRF imaging buffer (25 mM Tris pH 8, 150 mM KCl, 1mM TCEP) to obtain three different CaMKII stock concentrations – 1.6 μM, 0.4 μM and 0.04 μM. The recordings were made by diluting 5 μL of each stock sample in 15 μL of imaging buffer, directly on the cover slide. The final concentration of CaMKII^WT^ in the drop was 400 nM, 100 nM and 10 nM, respectively. Each dilution was measured three times. Activated sample was obtained by incubating 8 μM CaMKII^WT^, 1.5 μM Ca^2+^:CaM, 1 mM ATP, 2 mM MgCl_2_ in imaging buffer. The sample was left incubating for 5 min at room temperature before stock dilutions were made with 1.6 μM, 0.4 μM and 0.04 μM final CaMKII^WT^ by diluting the original incubation reaction in imaging buffer. Each sample dilution was measured three times, by diluting 5 μL of stock sample in 15 μL imaging buffer. The same procedure was repeated for CaMKII^K42R/D135N^-AviTag.

Percentages of monomers participating in each peak were calculated in the following way: first peak area percentages were calculated for each peak. Then, the number of monomers for each peak was calculated with respect to the area percentage. Finally, the percentage of monomers in each peak was calculated based on total particles identified in each measurement. We used IgorPro9 multipeak fitting function to fit the peaks obtained by the instrument. Here is an example of the calculation used for Figure 7B: Peak area % (calculated by the fitting software) for each peak is 4.7% for the 2-mer, 85.9% for the 12-mer, and 9.4% for the 24-mer. These are percentages of particles detected, not monomers. To calculate in terms of monomers, we multiplied each peak area % with the number of subunits and got 4.7×2 = 9.4 for 2-mer, 85.9×12 = 1030.8 for 12-mer and 9.4×24 = 225.6 for 24-mer. We summed these to obtain the total monomer number (1265.8) detected in this measurement. We obtained the fraction of monomers in each of the three peaks with respect to total number of particles, corresponding in this case to 0.7%, 81.5% and 17.8% of the monomers in the sample, respectively.

### Preparation of samples for Mass Spectrometry

First, proteins were buffer exchanged to HEPES containing buffer (25 mM HEPES pH 8, 250 mM NaCl, 1% glycerol, 1 mM TCEP) using PD-10 Sephadex desalting column (GE Healthcare). The following samples (50 μL final reaction volume) were prepared for kinase reaction: 8 μM ^14^N CaMKII^WT^ unactivated, 8 μM ^15^N CaMKII^WT^ unactivated, 4 μM ^14^N CaMKII^WT^ and 4 μM ^15^N CaMKII^WT^ unactivated, 8 μM ^14^N CaMKII^WT^ activated (with 1.5 μM Ca^2+^:CaM, 100 μM ATP, 200 μM MgCl_2_), 8 μM ^15^N CaMKII^WT^ activated (with 1.5 μM Ca^2+^:CaM, 100 μM ATP, 200 μM MgCl_2_), 4 μM N^14^ CaMKII^WT^ and 4 μM ^15^N CaMKII^WT^ activated (with 1.5 μM Ca^2+^:CaM, 100 μM ATP, 200 μM MgCl_2_). The samples were left for 30 min or for 150 min at 37°C. The samples were then moved to room temperature, and incubated with 2 mM DSS final concentration for 30 min, and then quenched with 30 mM Tris pH 8 for 15 min at room temperature. The samples were stored at −80°C until further use.

### Sample digestion for Mass Spectrometry

Samples were denatured with 8 M Urea and reduced with 5 mM dithiothreitol (DTT) at 37°C for 45 min, followed by alkylation with 40 mM chloroacetamide (CAA) for 45 min in the dark. The urea was then diluted to 1 M with 50 mM Triethylammonium bicarbonate buffer (TEAB), pH 8.0. Protein digestion was finished by using Lysyl endopeptidase C (Wako) with an enzyme-to-protein ratio 1:100 (w/w) for 3 h at 37°C and continued with trypsin (Serva) at an enzyme-to-protein ration 1:50 (w/w) for overnight at 37°C. Enzymically digestive samples were cleaned with C8 Sep-Pak (Waters), dried and stored at −20°C for further analysis.

### LC-MS analysis

The LC-MS analysis of crosslinked samples were conducted by Orbitrap Fusion Lumos Tribrid Mass Spectrometer (Thermo Fisher Scientific) coupled with UltiMate 3000 RSLC nano LC system (Thermo Fisher Scientific). The separation of samples was performed by 50 cm reverse-phase column (in-house packed with Poroshell 120 EC-C18, 2.7 μm, Agilent Technologies) with 180 min gradient. High field asymmetric waveform ion mobility spectrometry (FAIMS) was enabled with internal stepping −40/-50/-60 V. Cross-linked samples were acquired with 120,000 resolution at MS1 levels, 30,000 resolution at MS2 levels, charge state 4-8 enabled for MS2, higher-energy collisional dissociation (HCD) at 30% for MS2 fragmentation.

### Data analysis

All raw spectra were searched with pLink2 2.3.9 (Chen et al., 2019), against the protein sequence of CaMKII (Uniprot ID: Q9UQM7. Only ^14^N labeled crosslinks were searched. pLink parameters were as follows: minimum peptide length, 6; maximal peptide length, 60; missed cleavages, 3; Cys carbamidomethyl (57.021 Da) as fixed modification and Met oxidation (15.995 Da) as variable modification; LinkerMass was set to 138.068 Da and MonoMass was set to 156.079 Da; 10 ppm for precursor mass tolerance and 20 ppm for fragment mass tolerance; 1% separate FDR at CSM level. Raw files and search result were imported into in-house developed R-script using the rawrr package (Kockmann et al., 2021) to extract the MS1 intensities of uni-isotope and mixed-isotope crosslinks. For single spectrum of identified cross-linked peptides, the intensity ratio for crosslinks was evaluated with the following equation:

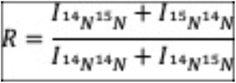

where *I* was the highest centroid peak intensity in centroid envelope from the MS^1^ mass spectrum.

### Structural analysis of crosslinks

Structures were visualized in ChimeraX (1.4, Pettersen et al., 2021). The XMAS plugin was used for a preliminary analysis. Subsequently, crosslinks were imported as pseudobonds between C-alpha atoms. The pseudobond files were generated from mass spectra plink files in Excel. To generate illustrative holoenzyme interactions and “close mode”, the holoenzymes were moved into place by hand, taking care to minimize close residue clashes.

## Acknowledgements

We thank Marcus Wietstruk for technical assistance, Martin Lehmann (FMP Imaging facility) for support with TIRF imaging, the Söllner Group (University of Heidelberg) for the plasmids encoding BzF synthetase and tRNA in *E. coli* and for advice on in vitro LED crosslinking, Sascha Lange (FMP Solid State NMR) for help with expression and purification of ^15^N-labelled CaMKII and Heike Stephanowitz (FMP Structural Interactomics) for intact mass determination. I.L. was recipient of a Marie Curie Incoming International Fellowship (798696) and “*Wiedereinstiegsstipendium*” from the Leibniz FMP. This work was funded by the DFG TRR 186 (Project A07, *Projektnummer* 272140445 to A.J.R.P.) and a DFG Heisenberg Professorship (to A.J.R.P., *Projektnummern* 323514590 & 446182550). Molecular graphics and analyses performed with UCSF ChimeraX, developed by the Resource for Biocomputing, Visualization, and Informatics at the University of California, San Francisco, with support from National Institutes of Health R01-GM129325 and the Office of Cyber Infrastructure and Computational Biology, National Institute of Allergy and Infectious Diseases.

