## Supplementary Figures for "CaMKII activity spreads by inter-holoenzyme phosphorylation"

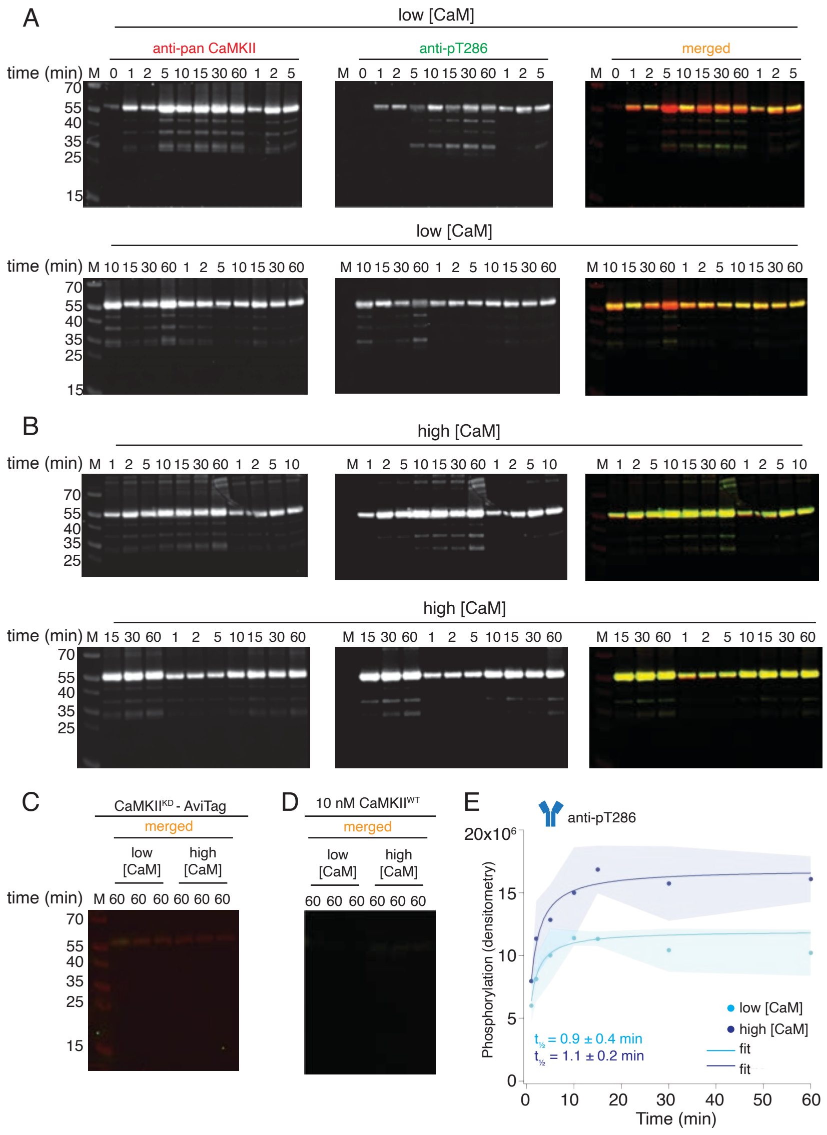


**Supplementary Figure S1. Western blot detection of CaMKII$^{KD}$ phosphorylation by CaMKII$^{WT}$.**

A) Blots of pT286 detection on CaMKII$^{KD}$ (4 µM) after phosphorylation with CaMKII$^{WT}$ (10 nM) in the presence of ATP:$Mg^{2+}$ and low $Ca^{2+}$:CaM concentrations (10 nM). Each time point is done in triplicate.

B) Blots of pT286 detection on CaMKII$^{KD}$ (4 µM), after phosphorylation with CaMKII$^{WT}$ (10 nM) in the presence of ATP:$Mg^{2+}$ and high $Ca^{2+}$:CaM concentrations (2 µM). Each time point is done in triplicate.

C) Western blot showing there is no phosphorylation signal from pT286 antibody on CaMKII$^{KD}$ (4 µM) when CaMKII$^{WT}$ is omitted from the kinase reaction.

D) Western blot showing that 10 nM CaMKII$^{WT}$ (as included on the gels in panels A and) gives no phosphorylation signal from the pT286 antibody.

E) Raw densitometry data from blots in panels A and B fitted with Langmuir function. Half-maximum times for CaMKII$^{KD}$ phosphorylation by CaMKII$^{WT}$ under low $Ca^{2+}$:CaM conditions is $t_{½} = 0.9 \pm 0.4$ min, and under high $Ca^{2+}$:CaM conditions is $t_{½} = 1.1 \pm 0.2$ min

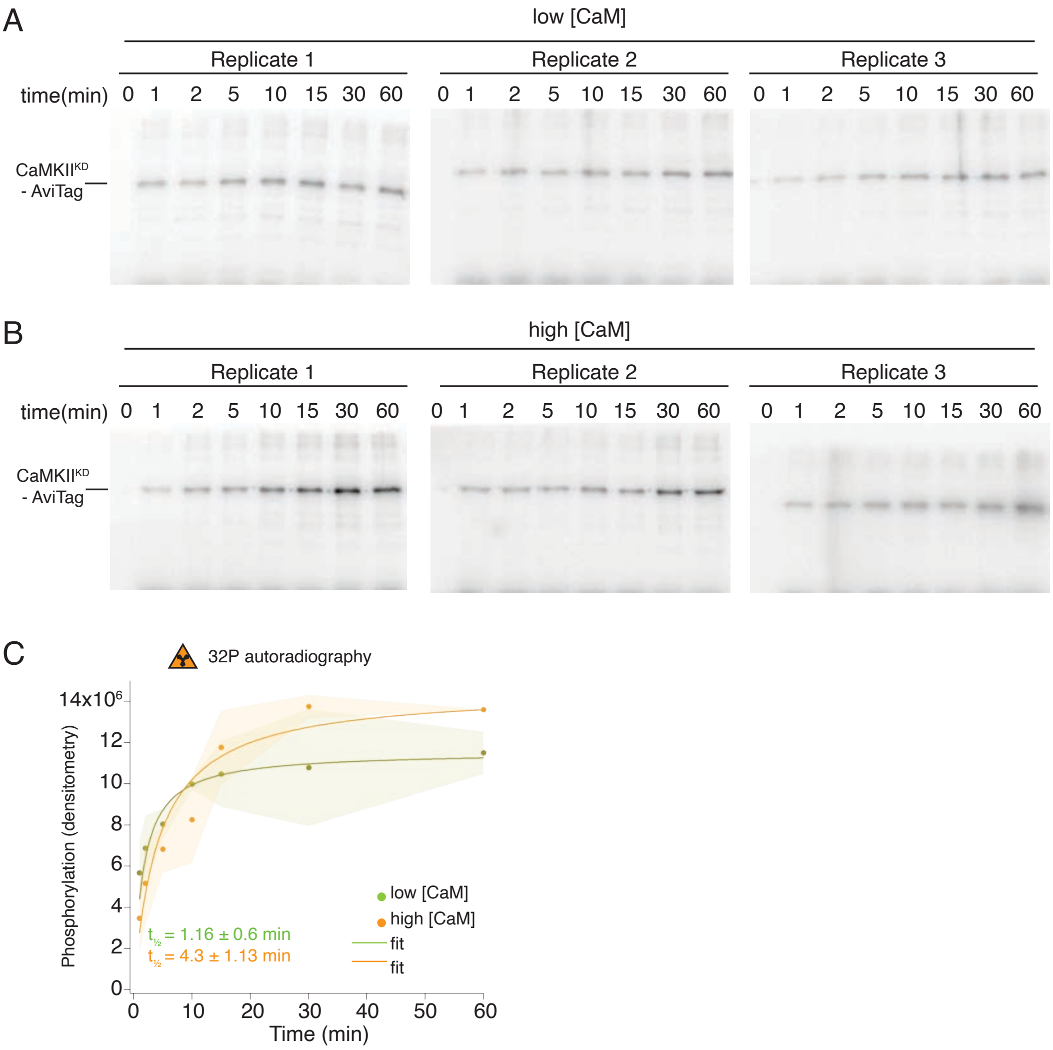


**Supplementary Figure S2. Radioactivity detection of CaMKII$^{KD}$ phosphorylation by CaMKII$^{WT}$.**

A) Gels of CaMKII$^{KD}$ phosphorylation detection in the presence of ATP:$Mg^{2+}$ and low $Ca^{2+}$:CaM concentrations (10 nM). Each time point is done in triplicates.

B) Gels of CaMKII$^{KD}$ phosphorylation detection in the presence of ATP:$Mg^{2+}$ and high $Ca^{2+}$:CaM concentrations (2 µM). Each time point is done in triplicates.

C) Raw densitometry data from gels in A and B fitted with Langmuir function. Half-maximum times for CaMKII$^{KD}$ phosphorylation by CaMKII$^{WT}$ under low $Ca^{2+}$:CaM conditions is $t_{½} = 1.16 \pm 0.6$ min, and under high $Ca^{2+}$:CaM conditions is $t_{½} = 4.3 \pm 1.13$ min.

Figure S3. Properties of CaMKII$^{F394BzF}$ : Kinase activity and size exclusion chromatography

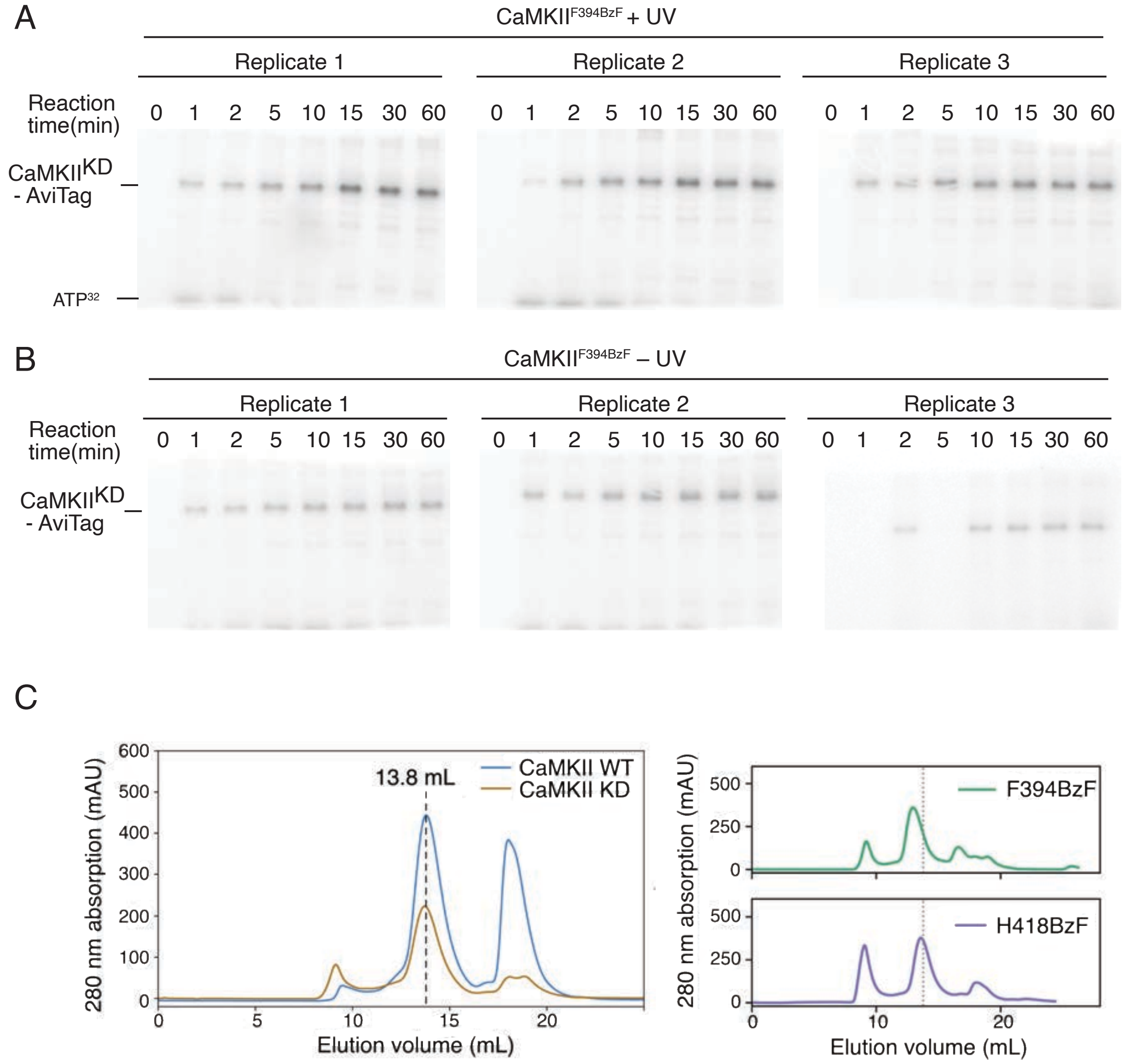


**Supplementary Figure S3. Radioactivity detection of CaMKII$^{KD}$ phosphorylation by CaMKII$^{F394BzF}$.**

A) Gels of CaMKII$^{KD}$ phosphorylation detection by UV treated CaMKII$^{F394BzF}$. Each time point is done in triplicates. These data are used for figure 2C.

B) Gels of CaMKII$^{KD}$ phosphorylation detection by untreated CaMKII$^{F394BzF}$. Each time point is done in triplicates. These data are used for figure 2C.

C) Size exclusion chromatography of CaMKII wild-type and mutants used in this study. Dashed lines indicate the peak elution volume for wild-type.

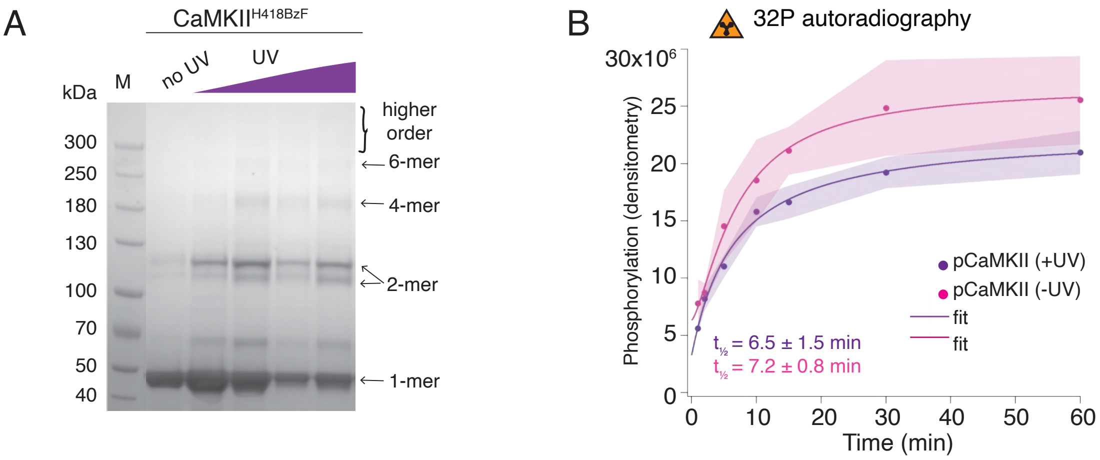


**Supplementary Figure S4. Hub domain mutant CaMKII$^{H418BzF}$ phosphorylates CaMKII$^{KD}$ irrespective of crosslinking.**

A) Coomassie stained gel showing UV-dependent oligomerisation of CaMKII$^{H418BzF}$.

B) Phosphorylation of CaMKII$^{KD}$ by UV-treated or -untreated CaMKII$^{H418BzF}$ . Langmuir fit determined half-maximum times of phosphorylation $t_{½}$ = 7.2 ± 0.8 min for untreated CaMKII$^{H418BzF}$ and $t_{½}$ = 6.5 ± 1.5 min for UV-treated CaMKII$^{H418BzF}$.

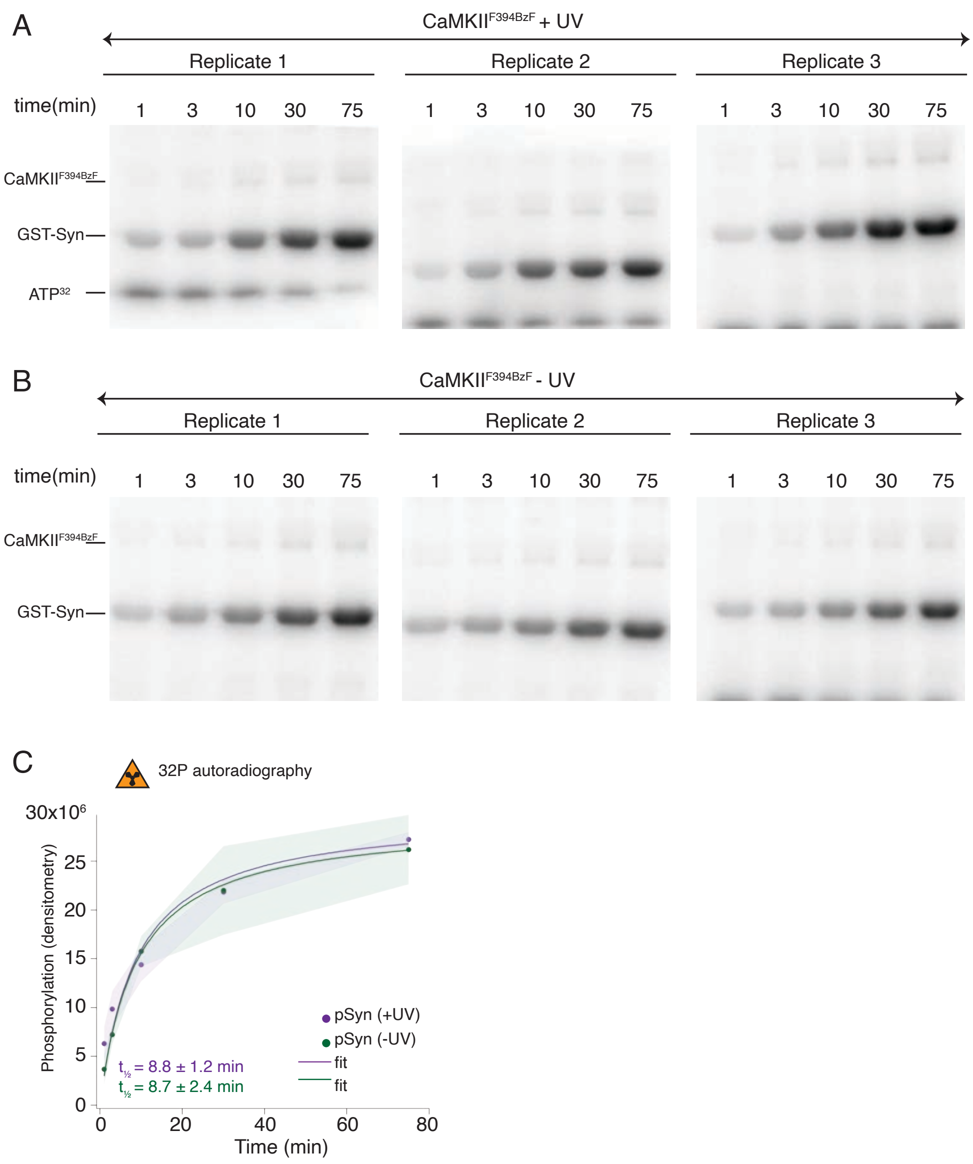


**Supplementary Figure S5. Radioactivity detection of substrate phosphorylation by CaMKII$^{F394BzF}$.**

A) Gels of CaMKII substrate (GST-Syn) phosphorylation by UV-treated CaMKII$^{F394BzF}$. Each time point is done in triplicate.

B) Gels of CaMKII substrate (GST-Syn) phosphorylation by control CaMKII$^{F394BzF}$ (no UV treatment). Each time point is done in triplicate.

C) Phosphorylation of GST-Syn by UV-treated or -untreated CaMKII$^{F394BzF}$. Langmuir fit determined half-maximum times of GST-Syn phosphorylation $t_{½}$ = 8.7 ± 2.4 min for untreated CaMKII$^{F394BzF}$ and $t_{½}$ = 8.8 ± 1.2 min for UV-treated CaMKII$^{F394BzF}$.

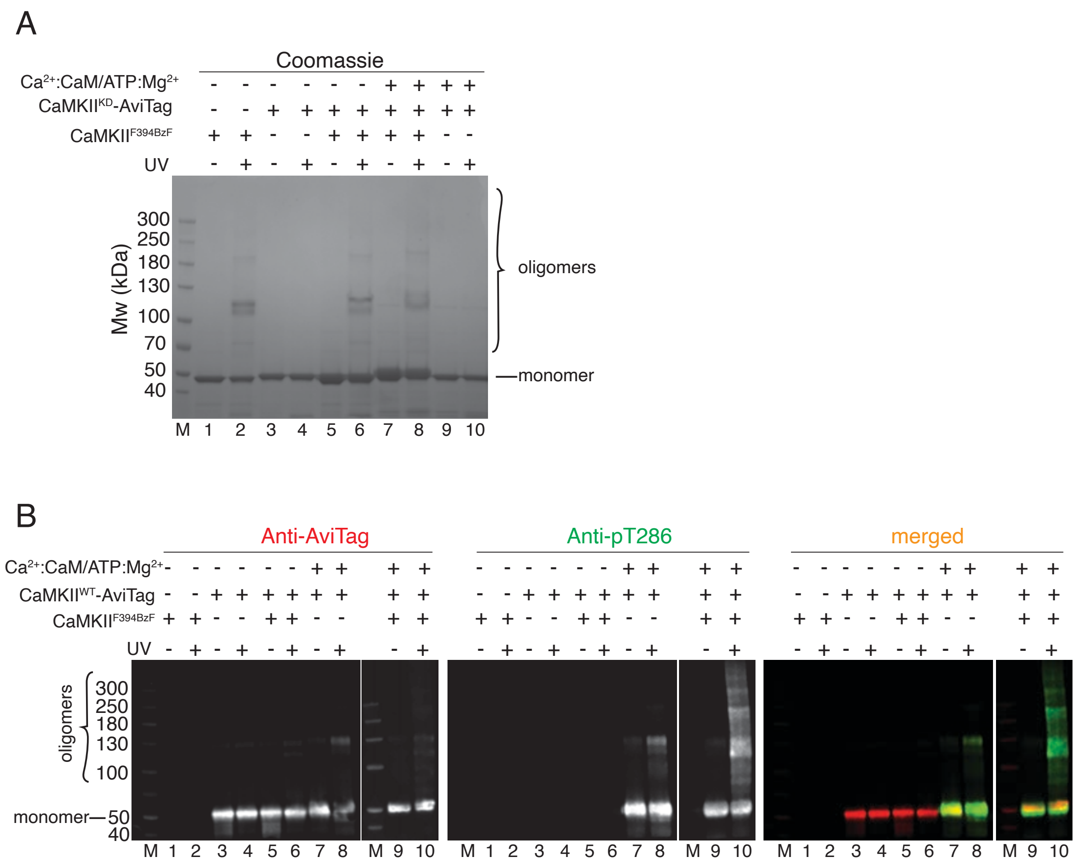


**Supplementary Figure S6. CaMKII holoenzymes do not mix during activation**

A) Coomassie stained gel of CaMKII$^{F394BzF}$ and CaMKII$^{KD}$ crosslinkning used in Figure 2E.

B) Western blot detection of possible CaMKII$^{WT}$-AviTag incorporation in CaMKII$^{F394BzF}$ holoenzymes.

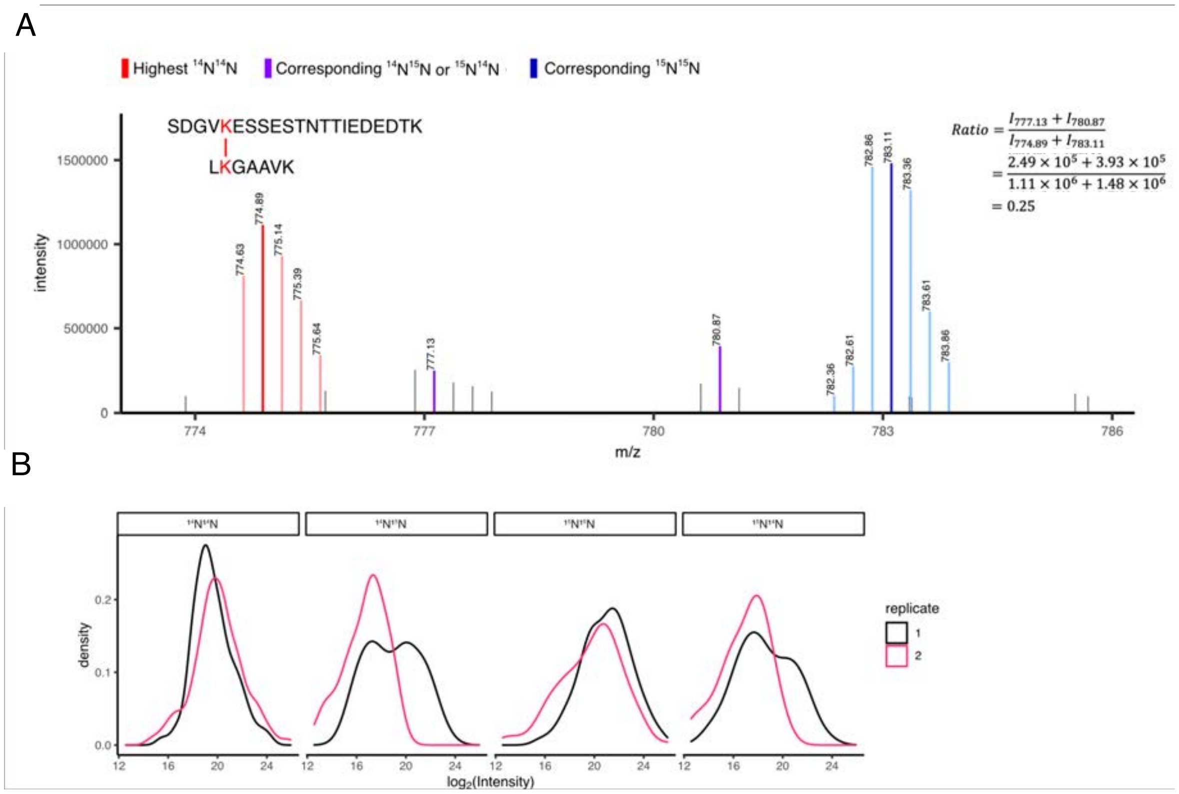


**Supplementary Figure S7. XL-MS identifies interactions of CaMKII holoenzymes**

A) Zoomed-in exemplary MS1 spectrum showing uni-isotopic (homotypic) and mixed-isotopic (heterotypic) crosslinks. The ratio of mixed-isotopic and uni-isotopic crosslinks is calculated using the intensities of the highest intense peak in each isotopic distribution.

B) The MS1 intensity distribution of the most intense peak of all four isotopic distributions (i.e., $^{14}N^{14}N$, $^{14}N^{15}N$, $^{15}N^{14}N$, $^{15}N^{15}N$) showing good reproducibility over replicates.

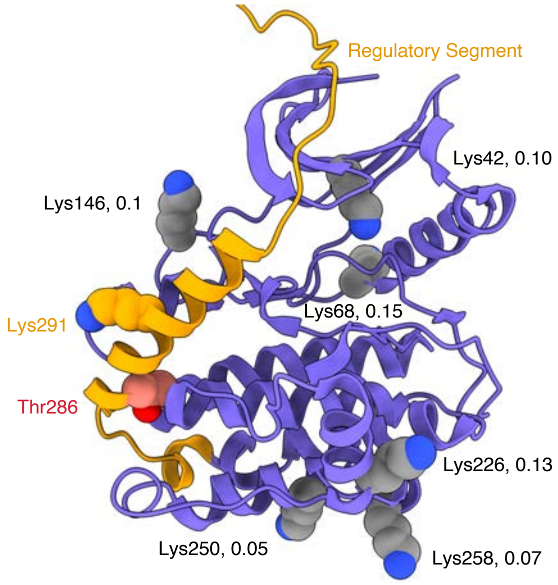


**Supplementary Figure S8. Crosslinking sites involving pT286 peptides from mixed isotypes**

Kinase domain (from the 5u6y PDB structure) with regulatory domain (orange) docked. 6 Lysine residues that gave heterotypic crosslinked peptides including P-Thr286 (DSS link to Lys291) are indicated as spheres. The detection ratio (heterotypic to homotypic mass spectra intensity) for each crosslinked interaction is indicated.

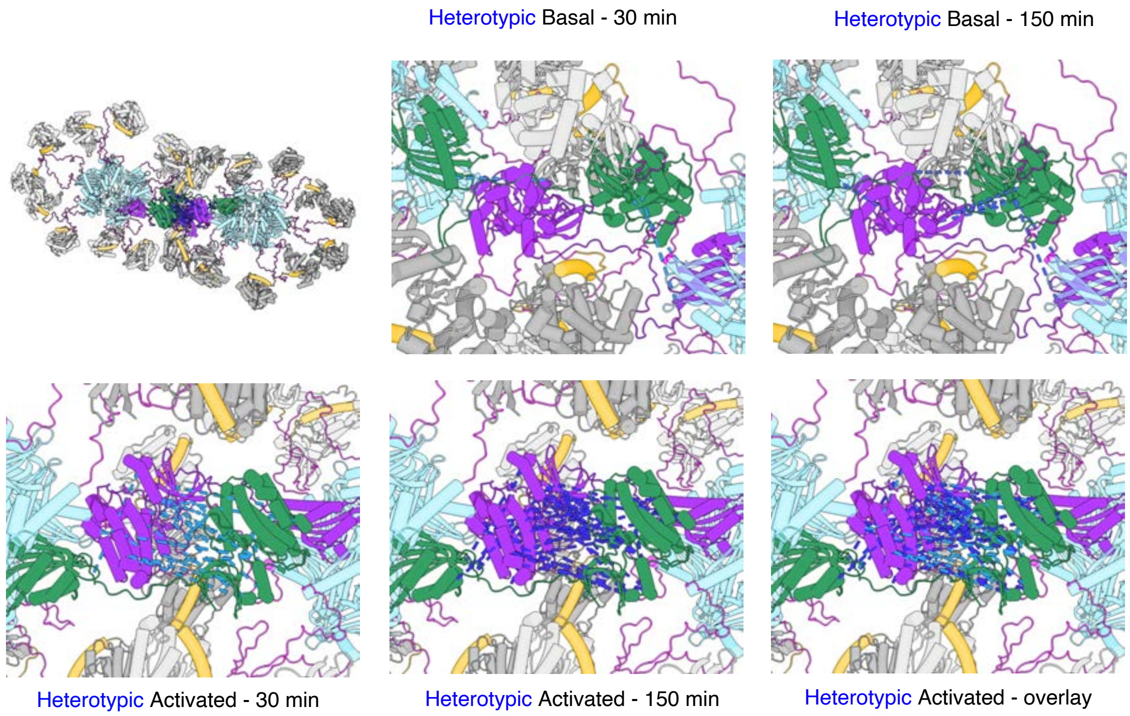


**Supplementary Figure S9. Close mode of holoenzyme interaction from MS crosslinks**

Heterotypic DSS crosslinks between kinase domains have plausible lengths (less than 20Å) when one kinase domain from a holoenzyme is interdigitated into the structure of another holoenzyme. Hub domains are in light blue, regulatory domains (docked) in orange. Docking was done by hand to minimize clashes (see text for details).

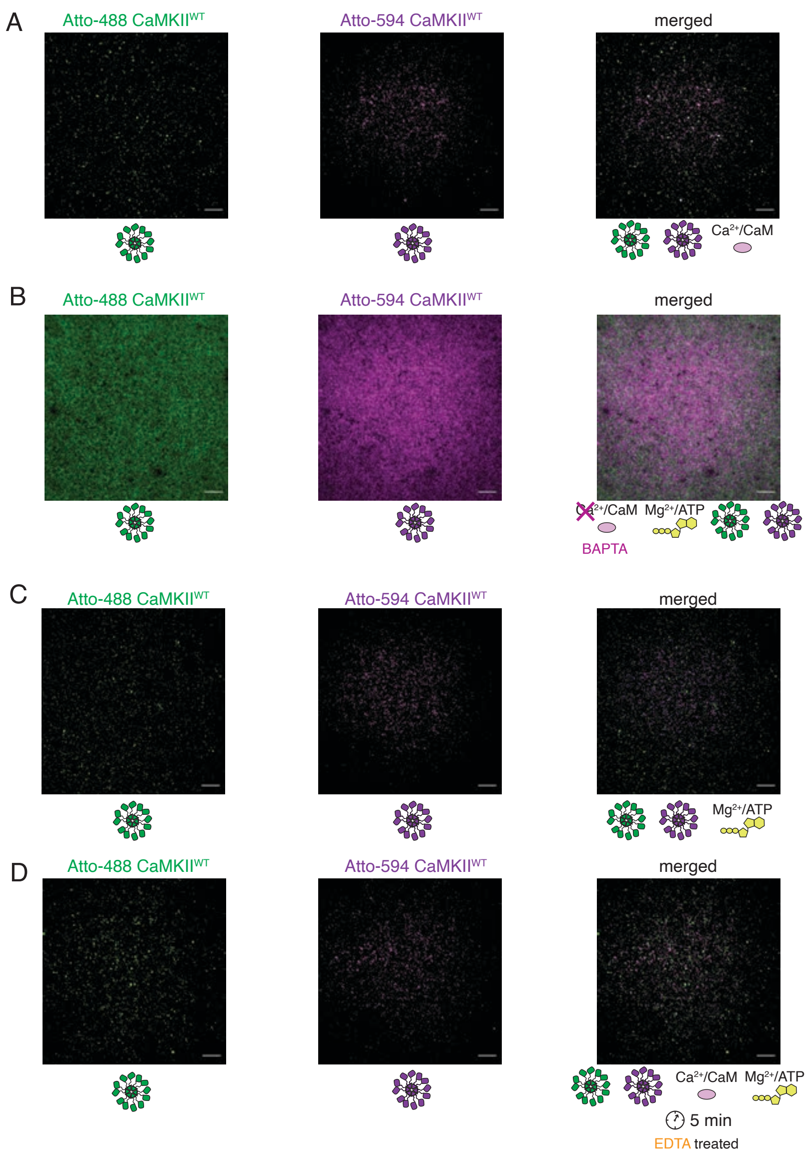


**Supplementary Figure S10. Activation stimuli are necessary for CaMKII$^{WT}$ holoenzyme colocalization**

A) Absence of colocalization between differently labeled CaMKII$^{WT}$ holoenzymes in the absence of $Mg^{2+}$:ATP

B) Absence of colocalization between differently labeled CaMKII$^{WT}$ holoenzymes upon chelation of $Ca^{2+}$ by BAPTA

C) Absence of colocalization between differently labeled CaMKII$^{WT}$ holoenzymes in the absence of CaM

D) Absence of colocalization between differently labeled CaMKII$^{WT}$ holoenzymes incubated first in activating conditions, and following later chelation of $Mg^{2+}$ by EDTA.

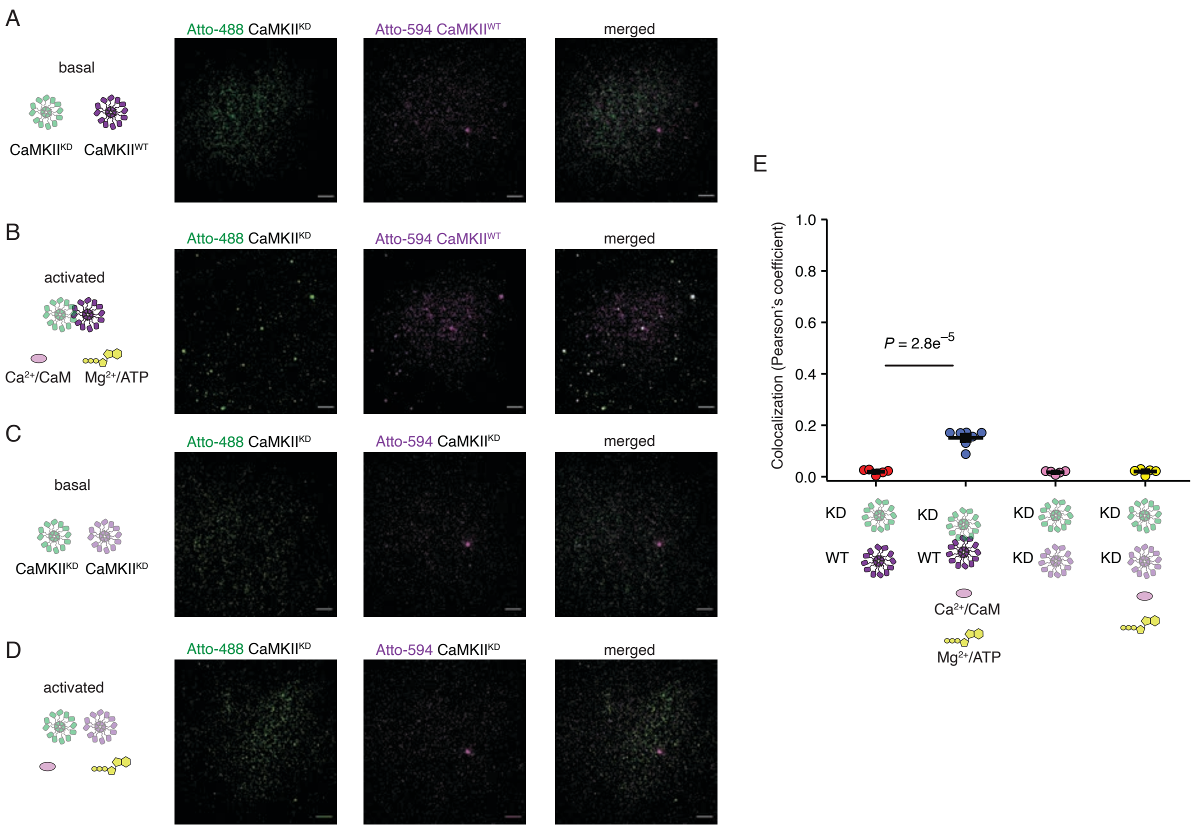


**Supplementary Figure S11. CaMKII$^{KD}$ holoenzymes cluster only in the presence of activated CaMKII$^{WT}$ holoenzymes**

A) Absence of colocalization between CaMKII$^{KD}$ and CaMKII$^{WT}$ in basal conditions

B) Colocalization of CaMKII$^{KD}$ and CaMKII$^{WT}$ upon activation

C) Absence of colocalization between differently labeled CaMKII$^{KD}$ in basal conditions

D) Absence of colocalization between differently labeled CaMKII$^{KD}$ in activating conditions

E) Summary of colocalization analysis. Probability of no difference from Dunnett's multiple comparison test.

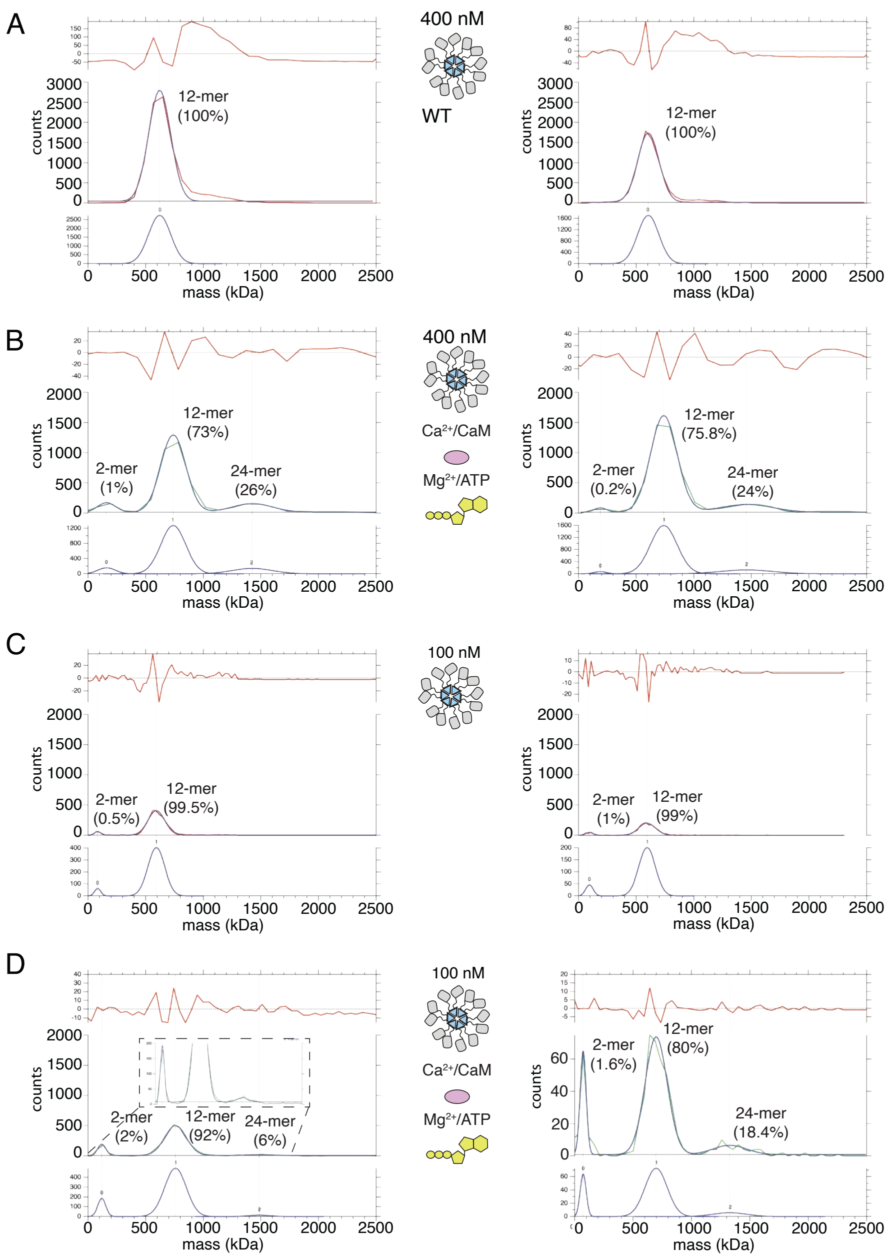


**Supplementary Figure S12. CaMKII$^{WT}$ forms higher order clusters during activation**

A) Two replicates of the particle mass distribution of 400nM CaMKII$^{WT}$ under basal conditions (red curve). Blue curve is multi-Gaussian fit (shown separately on lower graph). Red curve in upper graph is the fit residual.

B) Two replicates of the particle mass distribution of 400nM CaMKII$^{WT}$ under activating conditions (green curve).

C) Two replicates of the particle mass distribution of 400nM CaMKII$^{WT}$ under basal conditions (red curve).

D) Two replicates of the particle mass distribution of 100nM CaMKII$^{WT}$ under activating conditions (green curve).

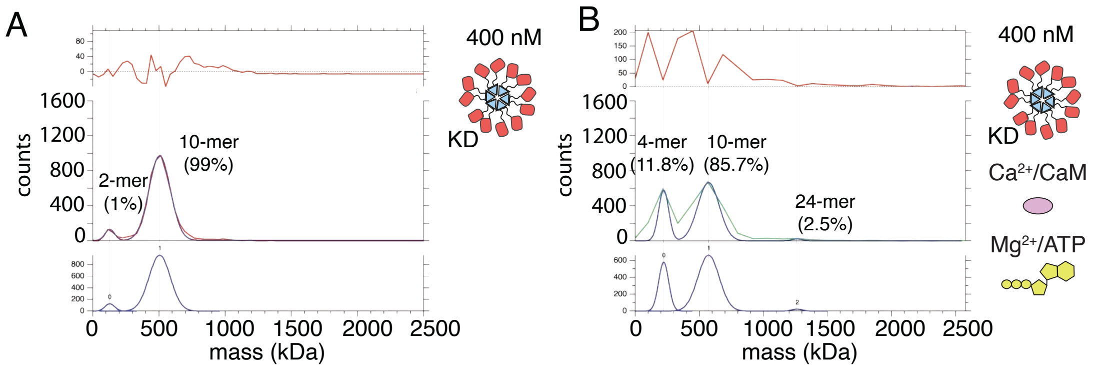


**Supplementary Figure S13. CaMKII$^{KD}$ fails to form higher order clusters during activation.**

A) Particle mass distribution of 400nM CaMKII$^{KD}$ under basal conditions.

B) Particle mass distribution of 400nM CaMKII$^{KD}$ under activating conditions.
