## Supplementary Tables for "CaMKII activity spreads by inter-holoenzyme phosphorylation"

Supplementary Table 1

| # | Peptide | domain_a | domain_b | crosslinked lysines |
|---|---|---|---|---|
| | **Homotypic crosslinks (basal, 30 min)** | | | |
| 1 | AGAYDFPSPEWDTVTPEAKDLINK(19)-NFSGGKSGGNK(6) | kinase | linker | 245-317 |
| 2 | CVKVLAGQEYAAK(3)-LLKHPNIVR(3) | kinase | kinase | 32-68 |
| 3 | DGKWQIVHFHR(3)-DHQKLER(4) | hub | kinase | 461-56 |
| 4 | DGKWQIVHFHR(3)-KQEIIK(1) | hub | hub | 461-347 |
| 5 | DGKWQIVHFHR(3)-KSDGVK(1) | hub | linker | 461-323 |
| 6 | DHQKLER(4)-DHQKLER(4) | kinase | kinase | 56-56 |
| 7 | DHQKLER(4)-IINTKK(5) | kinase | kinase | 56-47 |
| 8 | DHQKLER(4)-KSDGVK(1) | kinase | linker | 56-323 |
| 9 | DHQKLER(4)-LKGAAVK(2) | kinase | kinase | 56-148 |
| 10 | DLINKMLTINPSK(5)-DHQKLER(4) | kinase | kinase | 250-56 |
| 11 | DLINKMLTINPSK(5)-DLKPENLLLASK(3) | kinase | kinase | 250-137 |
| 12 | DLINKMLTINPSK(5)-KQEIIK(1) | kinase | hub | 250-347 |
| 13 | DLINKMLTINPSK(5)-LKGAAVK(2) | kinase | kinase | 250-148 |
| 14 | DLINKMLTINPSK(5)-MLTINPSKR(8) | kinase | kinase | 250-258 |
| 15 | DLKPENLLLASK(3)-DHQKLER(4) | kinase | kinase | 137-56 |
| 16 | DLKPENLLLASK(3)-DLKPENLLLASK(3) | kinase | kinase | 137-137 |
| 17 | DLKPENLLLASK(3)-KSDGVK(1) | kinase | linker | 137-323 |
| 18 | DLKPENLLLASK(3)-LKGAAVK(2) | kinase | kinase | 137-148 |
| 19 | DLKPENLLLASK(3)-LLKHPNIVR(3) | kinase | kinase | 137-68 |
| 20 | DLKPENLLLASK(3)-MLTINPSKR(8) | kinase | kinase | 137-258 |
| 21 | DLKPENLLLASK(3)-NFSGGKSGGNK(6) | kinase | linker | 137-317 |
| 22 | DLKPENLLLASKLK(12)-DHQKLER(4) | kinase | kinase | 146-56 |
| 23 | DLKPENLLLASKLK(12)-IINTKK(5) | kinase | kinase | 146-47 |
| 24 | DLKPENLLLASKLK(12)-KSDGVK(1) | kinase | linker | 146-323 |
| 25 | DLKPENLLLASKLK(12)-LKGAAVK(2) | kinase | kinase | 146-148 |
| 26 | DLKPENLLLASKLK(12)-LLKHPNIVR(3) | kinase | kinase | 146-68 |
| 27 | DLKPENLLLASKLK(12)-MLTINPSKR(8) | kinase | kinase | 146-258 |
| 28 | DLKPENLLLASKLK(12)-NFSGGKSGGNK(6) | kinase | linker | 146-317 |
| 29 | ESSESTNTTIEDEDTKVR(16)-DGKWQIVHFHR(3) | linker | hub | 344-461 |
| 30 | ESSESTNTTIEDEDTKVR(16)-DHQKLER(4) | hub | kinase | 344-56 |
| 31 | ESSESTNTTIEDEDTKVR(16)-ESSESTNTTIEDEDTKVR(16) | hub | hub | 344-344 |
| 32 | ESSESTNTTIEDEDTKVR(16)-KQEIIK(1) | hub | hub | 344-347 |
| 33 | ESSESTNTTIEDEDTKVR(16)-KSDGVK(1) | hub | linker | 344-323 |
| 34 | ESSESTNTTIEDEDTKVR(16)-LKGAAVK(2) | hub | kinase | 344-148 |
| 35 | ESSESTNTTIEDEDTKVR(16)-LKGAILTTMLATR(2) | hub | regulatory | 344-300 |

| | | | | |
|---|---|---|---|---|
| 36 | ESSESTNTTIEDEDTKVR(16)-LLKHPNIVR(3) | hub | kinase | 344-68 |
| 37 | ESSESTNTTIEDEDTKVR(16)-MLTINPSKR(8) | hub | kinase | 344-258 |
| 38 | ESSESTNTTIEDEDTKVR(16)-NFSGGKSGGNK(6) | hub | linker | 344-317 |
| 39 | ESSESTNTTIEDEDTKVRK(16)-KQEIIK(1) | hub | hub | 344-347 |
| 40 | FTEEYQLFEELGKGAFSVVR(13)-DGKWQIVHFHR(3) | kinase | hub | 21-461 |
| 41 | FTEEYQLFEELGKGAFSVVR(13)-DHQKLER(4) | kinase | kinase | 21-56 |
| 42 | FTEEYQLFEELGKGAFSVVR(13)-DLKPENLLLASK(3) | kinase | kinase | 21-137 |
| 43 | FTEEYQLFEELGKGAFSVVR(13)-DLKPENLLLASKLK(12) | kinase | kinase | 21-146 |
| 44 | FTEEYQLFEELGKGAFSVVR(13)-IINTKK(5) | kinase | kinase | 21-47 |
| 45 | FTEEYQLFEELGKGAFSVVR(13)-KSDGVK(1) | kinase | linker | 21-323 |
| 46 | FTEEYQLFEELGKGAFSVVR(13)-LKGAILTTMLATR(2) | kinase | regulatory | 21-300 |
| 47 | FTEEYQLFEELGKGAFSVVR(13)-MLTINPSKR(8) | kinase | kinase | 21-258 |
| 48 | FTEEYQLFEELGKGAFSVVR(13)-NFSGGKSGGNK(6) | kinase | linker | 21-317 |
| 49 | FTEEYQLFEELGKGAFSVVR(13)-VLAGQEYAAKIINTK(10) | kinase | kinase | 21-42 |
| 50 | FTEEYQLFEELGKGAFSVVRR(13)-IINTKKLSAR(5) | kinase | kinase | 21-47 |
| 51 | GAAVKLADFGLAIEVEGEQQAWFGFAGTPGYLSPEVLR(5)-LLKHPNIVR(3) | kinase | kinase | 153-68 |
| 52 | ITAAEALKHPWISHR(8)-DHQKLER(4) | kinase | kinase | 267-56 |
| 53 | ITAAEALKHPWISHR(8)-DLINKMLTINPSK(5) | kinase | kinase | 267-250 |
| 54 | ITAAEALKHPWISHR(8)-DLKPENLLLASK(3) | kinase | kinase | 267-137 |
| 55 | ITAAEALKHPWISHR(8)-KQEIIK(1) | kinase | hub | 267-347 |
| 56 | ITAAEALKHPWISHR(8)-KSDGVK(1) | kinase | linker | 267-323 |
| 57 | KLKGAILTTMLATR(3)-KLKGAILTTMLATR(1) | regulatory | regulatory | 300-298 |
| 58 | KQEIIK(1)-IINTKK(5) | hub | kinase | 347-47 |
| 59 | KQEIIK(1)-KQEIIK(1) | hub | hub | 347-347 |
| 60 | KQEIIK(1)-KSDGVK(1) | hub | linker | 347-323 |
| 61 | KQEIIK(1)-LKGAAVK(2) | hub | kinase | 347-148 |
| 62 | KSDGVKESSESTNTTIEDEDTK(1)-MLTINPSKR(8) | linker | kinase | 323-258 |
| 63 | KSDGVKESSESTNTTIEDEDTK(1)-NFSGGKSGGNK(6) | linker | linker | 323-317 |
| 64 | KSDGVKESSESTNTTIEDEDTK(6)-MLTINPSKR(8) | linker | kinase | 328-258 |
| 65 | KSDGVKESSESTNTTIEDEDTK(6)-NFSGGKSGGNK(6) | linker | linker | 328-317 |
| 66 | LKGAILTTMLATR(2)-DHQKLER(4) | regulatory | kinase | 300-56 |
| 67 | LKGAILTTMLATR(2)-DLKPENLLLASK(3) | regulatory | kinase | 300-137 |
| 68 | LKGAILTTMLATR(2)-KQEIIK(1) | regulatory | hub | 300-347 |
| 69 | LKGAILTTMLATR(2)-KSDGVK(1) | regulatory | linker | 300-323 |
| 70 | LKGAILTTMLATR(2)-LKGAAVK(2) | regulatory | kinase | 300-148 |
| 71 | LKGAILTTMLATR(2)-LKGAILTTMLATR(2) | regulatory | regulatory | 300-300 |
| 72 | LKGAILTTMLATR(2)-MLTINPSKR(8) | regulatory | kinase | 300-258 |

| | | | | |
|---|---|---|---|---|
| 73 | LKGAILTTMLATR(2)-NFSGGKSGGNK(6) | regulatory | linker | 300-317 |
| 74 | LKGAILTTMLATR(2)-QETVDCLKK(8) | regulatory | regulatory | 300-291 |
| 75 | LLKHPNIVR(3)-DHQKLER(4) | kinase | kinase | 68-56 |
| 76 | LLKHPNIVR(3)-LKGAAVK(2) | kinase | kinase | 68-148 |
| 77 | LYQQIKAGAYDFPSPEWDTVTPEAK(6)-DGKWQIVHFHR(3) | kinase | hub | 226-461 |
| 78 | LYQQIKAGAYDFPSPEWDTVTPEAK(6)-DHQKLER(4) | kinase | kinase | 226-56 |
| 79 | LYQQIKAGAYDFPSPEWDTVTPEAK(6)-DLKPENLLLASKLK(12) | kinase | kinase | 226-146 |
| 80 | LYQQIKAGAYDFPSPEWDTVTPEAK(6)-ITAAEALKHPWISHR(8) | kinase | kinase | 226-267 |
| 81 | LYQQIKAGAYDFPSPEWDTVTPEAK(6)-KQEIIK(1) | kinase | hub | 226-347 |
| 82 | LYQQIKAGAYDFPSPEWDTVTPEAK(6)-KSDGVK(1) | kinase | linker | 226-323 |
| 83 | LYQQIKAGAYDFPSPEWDTVTPEAK(6)-LKGAAVK(2) | kinase | kinase | 226-148 |
| 84 | LYQQIKAGAYDFPSPEWDTVTPEAK(6)-LKGAILTTMLATR(2) | kinase | regulatory | 226-300 |
| 85 | LYQQIKAGAYDFPSPEWDTVTPEAK(6)-LLKHPNIVR(3) | kinase | kinase | 226-68 |
| 86 | LYQQIKAGAYDFPSPEWDTVTPEAK(6)-MLTINPSKR(8) | kinase | kinase | 226-258 |
| 87 | LYQQIKAGAYDFPSPEWDTVTPEAK(6)-NFSGGKSGGNK(6) | kinase | linker | 226-317 |
| 88 | LYQQIKAGAYDFPSPEWDTVTPEAK(6)-SDGVKESSESTNTTIEDEDTK(5) | kinase | linker | 226-328 |
| 89 | LYQQIKAGAYDFPSPEWDTVTPEAK(6)-VLAGQEYAAKIINTK(10) | kinase | kinase | 226-42 |
| 90 | MLTINPSKR(8)-DHQKLER(4) | kinase | kinase | 258-56 |
| 91 | MLTINPSKR(8)-IINTKK(5) | kinase | kinase | 258-47 |
| 92 | MLTINPSKR(8)-KQEIIK(1) | kinase | hub | 258-347 |
| 93 | MLTINPSKR(8)-KSDGVK(1) | kinase | linker | 258-323 |
| 94 | MLTINPSKR(8)-LKGAAVK(2) | kinase | kinase | 258-148 |
| 95 | MLTINPSKR(8)-MLTINPSKR(8) | kinase | kinase | 258-258 |
| 96 | MLTINPSKR(8)-NFSGGKSGGNK(6) | kinase | linker | 258-317 |
| 97 | NFSGGKSGGNK(6)-KQEIIK(1) | linker | hub | 317-347 |
| 98 | NFSGGKSGGNK(6)-KSDGVK(1) | linker | linker | 317-323 |
| 99 | NFSGGKSGGNK(6)-LKGAAVK(2) | linker | kinase | 317-148 |
| 100 | NFSGGKSGGNK(6)-NFSGGKSGGNK(6) | linker | linker | 317-317 |
| 101 | NSKPVHTTILNPHIHLMGDESACIAYIR(3)-DHQKLER(4) | hub | kinase | 408-56 |
| 102 | QETVDCLKK(8)-DHQKLER(4) | regulatory | kinase | 291-56 |
| 103 | QETVDCLKK(8)-KQEIIK(1) | regulatory | hub | 291-347 |
| 104 | QETVDCLKK(8)-KSDGVK(1) | regulatory | linker | 291-323 |
| 105 | QETVDCLKK(8)-LLKHPNIVR(3) | regulatory | kinase | 291-68 |
| 106 | QETVDCLKK(8)-NFSGGKSGGNK(6) | regulatory | linker | 291-317 |
| 107 | RDGKWQIVHFHR(4)-SGGNKKSDGVK(5) | hub | linker | 461-322 |
| 108 | RITAAEALKHPWISHR(9)-DLINKMLTINPSKR(13) | kinase | kinase | 267-258 |
| 109 | RITAAEALKHPWISHR(9)-DLINKMLTINPSKR(5) | kinase | kinase | 267-250 |

| | | | | |
|---|---|---|---|---|
| 110 | RKLKGAILTTMLATR(2)-VLAGQEYAAKIINTK(10) | regulatory | kinase | 298-42 |
| 111 | RKLKGAILTTMLATR(4)-KSDGVK(1) | regulatory | linker | 300-323 |
| 112 | SDGVKESSESTNTTIEDEDTK(5)-DGKWQIVHFHR(3) | linker | hub | 328-461 |
| 113 | SDGVKESSESTNTTIEDEDTK(5)-DHQKLER(4) | linker | kinase | 328-56 |
| 114 | SDGVKESSESTNTTIEDEDTK(5)-DLKPENLLLASKLK(12) | linker | kinase | 328-146 |
| 115 | SDGVKESSESTNTTIEDEDTK(5)-ESSESTNTTIEDEDTKVR(16) | linker | linker | 328-344 |
| 116 | SDGVKESSESTNTTIEDEDTK(5)-IINTKK(5) | linker | kinase | 328-47 |
| 117 | SDGVKESSESTNTTIEDEDTK(5)-ITAAEALKHPWISHR(8) | linker | kinase | 328-267 |
| 118 | SDGVKESSESTNTTIEDEDTK(5)-KQEIIK(1) | linker | hub | 328-347 |
| 119 | SDGVKESSESTNTTIEDEDTK(5)-LKGAILTTMLATR(2) | linker | regulatory | 328-300 |
| 120 | SDGVKESSESTNTTIEDEDTK(5)-LLKHPNIVR(3) | linker | kinase | 328-68 |
| 121 | SDGVKESSESTNTTIEDEDTK(5)-MLTINPSKR(8) | linker | kinase | 328-258 |
| 122 | SDGVKESSESTNTTIEDEDTK(5)-NFSGGKSGGNK(6) | linker | linker | 328-317 |
| 123 | SDGVKESSESTNTTIEDEDTK(5)-RKLKGAILTTMLATR(2) | linker | regulatory | 328-298 |
| 124 | SDGVKESSESTNTTIEDEDTK(5)-VLAGQEYAAKIINTK(10) | linker | kinase | 328-42 |
| 125 | VLAGQEYAAKIINTK(10)-DHQKLER(4) | kinase | kinase | 42-56 |
| 126 | VLAGQEYAAKIINTK(10)-DLKPENLLLASK(3) | kinase | kinase | 42-137 |
| 127 | VLAGQEYAAKIINTK(10)-DLKPENLLLASKLK(12) | kinase | kinase | 42-146 |
| 128 | VLAGQEYAAKIINTK(10)-IINTKK(5) | kinase | kinase | 42-47 |
| 129 | VLAGQEYAAKIINTK(10)-KSDGVK(1) | kinase | linker | 42-323 |
| 130 | VLAGQEYAAKIINTK(10)-LKGAAVK(2) | kinase | kinase | 42-148 |
| 131 | VLAGQEYAAKIINTK(10)-LKGAILTTMLATR(2) | kinase | regulatory | 42-300 |
| 132 | VLAGQEYAAKIINTK(10)-LLKHPNIVR(3) | kinase | kinase | 42-68 |
| 133 | VLAGQEYAAKIINTK(10)-MLTINPSKR(8) | kinase | kinase | 42-258 |
| 134 | VLAGQEYAAKIINTK(10)-NFSGGKSGGNK(6) | kinase | linker | 42-317 |
| 135 | VLAGQEYAAKIINTK(10)-VLAGQEYAAKIINTK(10) | kinase | kinase | 42-42 |
| 136 | VLAGQEYAAKIINTKK(10)-DHQKLER(4) | kinase | kinase | 42-56 |

### Homotypic crosslinks (basal, 150 min)

| # | Peptide | domain_a | domain_b | crosslinked lysines |
|---|---|---|---|---|
| 1 | AGAYDFPSPEWDTVTPEAKDLINK(19)-DHQKLER(4) | kinase | kinase | 245-56 |
| 2 | AGAYDFPSPEWDTVTPEAKDLINK(19)-DLKPENLLLASK(3) | kinase | kinase | 245-137 |
| 3 | AGAYDFPSPEWDTVTPEAKDLINK(19)-IINTKK(5) | kinase | kinase | 245-47 |
| 4 | AGAYDFPSPEWDTVTPEAKDLINK(19)-ITAAEALKHPWISHR(8) | kinase | kinase | 245-267 |
| 5 | AGAYDFPSPEWDTVTPEAKDLINK(19)-KSDGVK(1) | kinase | linker | 245-323 |
| 6 | AGAYDFPSPEWDTVTPEAKDLINK(19)-LKGAAVK(2) | kinase | kinase | 245-148 |
| 7 | AGAYDFPSPEWDTVTPEAKDLINK(19)-LLKHPNIVR(3) | kinase | kinase | 245-68 |
| 8 | AGAYDFPSPEWDTVTPEAKDLINK(19)-MLTINPSKR(8) | kinase | kinase | 245-258 |
| 9 | AGAYDFPSPEWDTVTPEAKDLINK(19)-NFSGGKSGGNK(6) | kinase | linker | 245-317 |
| 10 | CVKVLAGQEYAAK(3)-DHQKLER(4) | kinase | kinase | 32-56 |
| 11 | DGKWQIVHFHR(3)-KQEIIK(1) | hub | hub | 461-347 |
| 12 | DGKWQIVHFHR(3)-KSDGVK(1) | hub | linker | 461-323 |
| 13 | DHQKLER(4)-DHQKLER(4) | kinase | kinase | 56-56 |
| 14 | DHQKLER(4)-IINTKK(5) | kinase | kinase | 56-47 |
| 15 | DHQKLER(4)-KQEIIK(1) | kinase | hub | 56-347 |
| 16 | DHQKLER(4)-KSDGVK(1) | kinase | linker | 56-323 |
| 17 | DHQKLER(4)-LKGAAVK(2) | kinase | kinase | 56-148 |
| 18 | DLINKMLTINPSK(5)-DHQKLER(4) | kinase | kinase | 250-56 |
| 19 | DLINKMLTINPSK(5)-IINTKK(5) | kinase | kinase | 250-47 |
| 20 | DLINKMLTINPSK(5)-KQEIIK(1) | kinase | hub | 250-347 |
| 21 | DLINKMLTINPSK(5)-KSDGVK(1) | kinase | linker | 250-323 |
| 22 | DLINKMLTINPSK(5)-LKGAAVK(2) | kinase | kinase | 250-148 |
| 23 | DLINKMLTINPSK(5)-MLTINPSKR(8) | kinase | kinase | 250-258 |
| 24 | DLINKMLTINPSK(5)-NFSGGKSGGNK(6) | kinase | linker | 250-317 |
| 25 | DLKPENLLLASK(3)-DHQKLER(4) | kinase | kinase | 137-56 |
| 26 | DLKPENLLLASK(3)-DLKPENLLLASK(3) | kinase | kinase | 137-137 |
| 27 | DLKPENLLLASK(3)-IINTKK(5) | kinase | kinase | 137-47 |
| 28 | DLKPENLLLASK(3)-KSDGVK(1) | kinase | linker | 137-323 |
| 29 | DLKPENLLLASK(3)-LKGAAVK(2) | kinase | kinase | 137-148 |
| 30 | DLKPENLLLASK(3)-MLTINPSKR(8) | kinase | kinase | 137-258 |
| 31 | DLKPENLLLASK(3)-NFSGGKSGGNK(6) | kinase | linker | 137-317 |
| 32 | DLKPENLLLASK(3)-QETVDCLKK(8) | kinase | regulatory | 137-291 |
| 33 | DLKPENLLLASKLK(12)-DHQKLER(4) | kinase | kinase | 146-56 |
| 34 | DLKPENLLLASKLK(12)-DLKPENLLLASKLK(12) | kinase | kinase | 146-146 |
| 35 | DLKPENLLLASKLK(12)-LKGAAVK(2) | kinase | kinase | 146-148 |

| | | | | |
|---|---|---|---|---|
| 36 | DLKPENLLLASKLK(12)-MLTINPSKR(8) | kinase | kinase | 146-258 |
| 37 | DLKPENLLLASKLK(12)-NFSGGKSGGNK(6) | kinase | linker | 146-317 |
| 38 | ESSESTNTTIEDEDTKVR(16)-DHQKLER(4) | hub | kinase | 344-56 |
| 39 | ESSESTNTTIEDEDTKVR(16)-DLKPENLLLASKLK(12) | hub | kinase | 344-146 |
| 40 | ESSESTNTTIEDEDTKVR(16)-ESSESTNTTIEDEDTKVR(16) | hub | hub | 344-344 |
| 41 | ESSESTNTTIEDEDTKVR(16)-ITAAEALKHPWISHR(8) | hub | kinase | 344-267 |
| 42 | ESSESTNTTIEDEDTKVR(16)-KQEIIK(1) | hub | hub | 344-347 |
| 43 | ESSESTNTTIEDEDTKVR(16)-KSDGVK(1) | hub | linker | 344-323 |
| 44 | ESSESTNTTIEDEDTKVR(16)-LKGAAVK(2) | hub | kinase | 344-148 |
| 45 | ESSESTNTTIEDEDTKVR(16)-LKGAILTTMLATR(2) | hub | regulatory | 344-300 |
| 46 | ESSESTNTTIEDEDTKVR(16)-MLTINPSKR(8) | hub | kinase | 344-258 |
| 47 | ESSESTNTTIEDEDTKVR(16)-NFSGGKSGGNK(6) | hub | linker | 344-317 |
| 48 | ESSESTNTTIEDEDTKVR(16)-RKLKGAILTTMLATR(2) | hub | regulatory | 344-298 |
| 49 | ESSESTNTTIEDEDTKVRK(16)-KQEIIK(1) | hub | hub | 344-347 |
| 50 | ESSESTNTTIEDEDTKVRK(16)-KSDGVK(1) | hub | linker | 344-323 |
| 51 | FTEEYQLFEELGKGAFSVVR(13)-DGKWQIVHFHR(3) | kinase | hub | 21-461 |
| 52 | FTEEYQLFEELGKGAFSVVR(13)-DHQKLER(4) | kinase | kinase | 21-56 |
| 53 | FTEEYQLFEELGKGAFSVVR(13)-DLKPENLLLASK(3) | kinase | kinase | 21-137 |
| 54 | FTEEYQLFEELGKGAFSVVR(13)-DLKPENLLLASKLK(12) | kinase | kinase | 21-146 |
| 55 | FTEEYQLFEELGKGAFSVVR(13)-IINTKK(5) | kinase | kinase | 21-47 |
| 56 | FTEEYQLFEELGKGAFSVVR(13)-ITAAEALKHPWISHR(8) | kinase | kinase | 21-267 |
| 57 | FTEEYQLFEELGKGAFSVVR(13)-KQEIIK(1) | kinase | hub | 21-347 |
| 58 | FTEEYQLFEELGKGAFSVVR(13)-KSDGVK(1) | kinase | linker | 21-323 |
| 59 | FTEEYQLFEELGKGAFSVVR(13)-LKGAAVK(2) | kinase | kinase | 21-148 |
| 60 | FTEEYQLFEELGKGAFSVVR(13)-LKGAILTTMLATR(2) | kinase | regulatory | 21-300 |
| 61 | FTEEYQLFEELGKGAFSVVR(13)-MLTINPSKR(8) | kinase | kinase | 21-258 |
| 62 | FTEEYQLFEELGKGAFSVVR(13)-NFSGGKSGGNK(6) | kinase | linker | 21-317 |
| 63 | FTEEYQLFEELGKGAFSVVR(13)-VLAGQEYAAKIINTK(10) | kinase | kinase | 21-42 |
| 64 | FTEEYQLFEELGKGAFSVVRR(13)-IINTKKLSAR(5) | kinase | kinase | 21-47 |
| 65 | FTEEYQLFEELGKGAFSVVRR(13)-IINTKKLSAR(6) | kinase | kinase | 21-48 |
| 66 | GAAVKLADFGLAIEVEGEQQAWFGFAGTPGYLSPEVLR(5)-LLKHPNIVR(3) | kinase | kinase | 153-68 |
| 67 | IINTKK(5)-IINTKK(5) | kinase | kinase | 47-47 |
| 68 | IINTKK(5)-LKGAAVK(2) | kinase | kinase | 47-148 |
| 69 | ITAAEALKHPWISHR(8)-DLINKMLTINPSK(5) | kinase | kinase | 267-250 |
| 70 | ITAAEALKHPWISHR(8)-DLKPENLLLASK(3) | kinase | kinase | 267-137 |
| 71 | ITAAEALKHPWISHR(8)-KQEIIK(1) | kinase | hub | 267-347 |
| 72 | ITAAEALKHPWISHR(8)-KSDGVK(1) | kinase | linker | 267-323 |

| | | | | |
|---|---|---|---|---|
| 73 | ITAAEALKHPWISHR(8)-MLTINPSKR(8) | kinase | kinase | 267-258 |
| 74 | KLKGAILTTMLATR(3)-KLKGAILTTMLATR(1) | regulatory | regulatory | 300-298 |
| 75 | KQEIIK(1)-IINTKK(5) | hub | kinase | 347-47 |
| 76 | KQEIIK(1)-KQEIIK(1) | hub | hub | 347-347 |
| 77 | KQEIIK(1)-KSDGVK(1) | hub | linker | 347-323 |
| 78 | KQEIIK(1)-LKGAAVK(2) | hub | kinase | 347-148 |
| 79 | KQEIIKVTEQLIEAISNGGFESYTK(1)-RDGKWQIVHFHR(4) | hub | hub | 347-461 |
| 80 | KQEIIKVTEQLIEAISNGGFESYTK(6)-DHQKLER(4) | hub | kinase | 352-56 |
| 81 | KQEIIKVTEQLIEAISNGGFESYTK(6)-NFSGGKSGGNK(6) | hub | linker | 352-317 |
| 82 | KSDGVKESSESTNTTIEDEDTK(6)-KQEIIK(1) | linker | hub | 328-347 |
| 83 | KSDGVKESSESTNTTIEDEDTK(6)-LKGAILTTMLATR(2) | linker | regulatory | 328-300 |
| 84 | KSDGVKESSESTNTTIEDEDTK(6)-NFSGGKSGGNK(6) | linker | linker | 328-317 |
| 85 | KSDGVKESSESTNTTIEDEDTKVR(22)-KQEIIK(1) | hub | hub | 344-347 |
| 86 | LKGAAVK(2)-LKGAAVK(2) | kinase | kinase | 148-148 |
| 87 | LKGAILTTMLATR(2)-DHQKLER(4) | regulatory | kinase | 300-56 |
| 88 | LKGAILTTMLATR(2)-DLKPENLLLASK(3) | regulatory | kinase | 300-137 |
| 89 | LKGAILTTMLATR(2)-KQEIIK(1) | regulatory | hub | 300-347 |
| 90 | LKGAILTTMLATR(2)-KSDGVK(1) | regulatory | linker | 300-323 |
| 91 | LKGAILTTMLATR(2)-MLTINPSKR(8) | regulatory | kinase | 300-258 |
| 92 | LKGAILTTMLATR(2)-NFSGGKSGGNK(6) | regulatory | linker | 300-317 |
| 93 | LKGAILTTMLATR(2)-QETVDCLKK(8) | regulatory | regulatory | 300-291 |
| 94 | LLKHPNIVR(3)-DHQKLER(4) | kinase | kinase | 68-56 |
| 95 | LLKHPNIVR(3)-KQEIIK(1) | kinase | hub | 68-347 |
| 96 | LLKHPNIVR(3)-LKGAAVK(2) | kinase | kinase | 68-148 |
| 97 | LLKHPNIVR(3)-MLTINPSKR(8) | kinase | kinase | 68-258 |
| 98 | LYQQIKAGAYDFPSPEWDTVTPEAK(6)-DGKWQIVHFHR(3) | kinase | hub | 226-461 |
| 99 | LYQQIKAGAYDFPSPEWDTVTPEAK(6)-DHQKLER(4) | kinase | kinase | 226-56 |
| 100 | LYQQIKAGAYDFPSPEWDTVTPEAK(6)-DLINKMLTINPSK(5) | kinase | kinase | 226-250 |
| 101 | LYQQIKAGAYDFPSPEWDTVTPEAK(6)-DLKPENLLLASK(3) | kinase | kinase | 226-137 |
| 102 | LYQQIKAGAYDFPSPEWDTVTPEAK(6)-DLKPENLLLASKLK(12) | kinase | kinase | 226-146 |
| 103 | LYQQIKAGAYDFPSPEWDTVTPEAK(6)-ESSESTNTTIEDEDTKVR(16) | kinase | hub | 226-344 |
| 104 | LYQQIKAGAYDFPSPEWDTVTPEAK(6)-IINTKK(5) | kinase | kinase | 226-47 |
| 105 | LYQQIKAGAYDFPSPEWDTVTPEAK(6)-ITAAEALKHPWISHR(8) | kinase | kinase | 226-267 |
| 106 | LYQQIKAGAYDFPSPEWDTVTPEAK(6)-KQEIIK(1) | kinase | hub | 226-347 |
| 107 | LYQQIKAGAYDFPSPEWDTVTPEAK(6)-KSDGVK(1) | kinase | linker | 226-323 |
| 108 | LYQQIKAGAYDFPSPEWDTVTPEAK(6)-LKGAAVK(2) | kinase | kinase | 226-148 |
| 109 | LYQQIKAGAYDFPSPEWDTVTPEAK(6)-LKGAILTTMLATR(2) | kinase | regulatory | 226-300 |

| | | | | |
|---|---|---|---|---|
| 110 | LYQQIKAGAYDFPSPEWDTVTPEAK(6)-LLKHPNIVR(3) | kinase | kinase | 226-68 |
| 111 | LYQQIKAGAYDFPSPEWDTVTPEAK(6)-LYQQIKAGAYDFPSPEWDTVTPEAK(6) | kinase | kinase | 226-226 |
| 112 | LYQQIKAGAYDFPSPEWDTVTPEAK(6)-MLTINPSKR(8) | kinase | kinase | 226-258 |
| 113 | LYQQIKAGAYDFPSPEWDTVTPEAK(6)-NFSGGKSGGNK(6) | kinase | linker | 226-317 |
| 114 | LYQQIKAGAYDFPSPEWDTVTPEAK(6)-SDGVKESSESTNTTIEDEDTK(5) | kinase | linker | 226-328 |
| 115 | LYQQIKAGAYDFPSPEWDTVTPEAK(6)-VLAGQEYAAKIINTK(10) | kinase | kinase | 226-42 |
| 116 | MLTINPSKR(8)-DHQKLER(4) | kinase | kinase | 258-56 |
| 117 | MLTINPSKR(8)-IINTKK(5) | kinase | kinase | 258-47 |
| 118 | MLTINPSKR(8)-KQEIIK(1) | kinase | hub | 258-347 |
| 119 | MLTINPSKR(8)-KSDGVK(1) | kinase | linker | 258-323 |
| 120 | MLTINPSKR(8)-LKGAAVK(2) | kinase | kinase | 258-148 |
| 121 | MLTINPSKR(8)-MLTINPSKR(8) | kinase | kinase | 258-258 |
| 122 | MLTINPSKR(8)-NFSGGKSGGNK(6) | kinase | linker | 258-317 |
| 123 | NFSGGKSGGNK(6)-DHQKLER(4) | linker | kinase | 317-56 |
| 124 | NFSGGKSGGNK(6)-KQEIIK(1) | linker | hub | 317-347 |
| 125 | NFSGGKSGGNK(6)-KSDGVK(1) | linker | linker | 317-323 |
| 126 | NFSGGKSGGNK(6)-LKGAAVK(2) | linker | kinase | 317-148 |
| 127 | NFSGGKSGGNK(6)-NFSGGKSGGNK(6) | linker | linker | 317-317 |
| 128 | QEIIKVTEQLIEAISNGGFESYTK(5)-DGKWQIVHFHR(3) | hub | hub | 352-461 |
| 129 | QEIIKVTEQLIEAISNGGFESYTK(5)-DHQKLER(4) | hub | kinase | 352-56 |
| 130 | QEIIKVTEQLIEAISNGGFESYTK(5)-MLTINPSKR(8) | hub | kinase | 352-258 |
| 131 | QEIIKVTEQLIEAISNGGFESYTK(5)-NFSGGKSGGNK(6) | hub | linker | 352-317 |
| 132 | QETVDCLKK(8)-DHQKLER(4) | regulatory | kinase | 291-56 |
| 133 | QETVDCLKK(8)-KQEIIK(1) | regulatory | hub | 291-347 |
| 134 | QETVDCLKK(8)-KSDGVK(1) | regulatory | linker | 291-323 |
| 135 | QETVDCLKK(8)-LLKHPNIVR(3) | regulatory | kinase | 291-68 |
| 136 | QETVDCLKK(8)-NFSGGKSGGNK(6) | regulatory | linker | 291-317 |
| 137 | RDGKWQIVHFHR(4)-SGGNKKSDGVK(5) | hub | linker | 461-322 |
| 138 | RKLKGAILTTMLATR(2)-DLKPENLLLASK(3) | regulatory | kinase | 298-137 |
| 139 | RKLKGAILTTMLATR(2)-RKLKGAILTTMLATR(2) | regulatory | regulatory | 298-298 |
| 140 | RKLKGAILTTMLATR(2)-VLAGQEYAAKIINTK(10) | regulatory | kinase | 298-42 |
| 141 | RKLKGAILTTMLATR(4)-DHQKLER(4) | regulatory | kinase | 300-56 |
| 142 | SDGVKESSESTNTTIEDEDTK(5)-DGKWQIVHFHR(3) | linker | hub | 328-461 |
| 143 | SDGVKESSESTNTTIEDEDTK(5)-DHQKLER(4) | linker | kinase | 328-56 |
| 144 | SDGVKESSESTNTTIEDEDTK(5)-DLINKMLTINPSK(5) | linker | kinase | 328-250 |
| 145 | SDGVKESSESTNTTIEDEDTK(5)-DLKPENLLLASK(3) | linker | kinase | 328-137 |
| 146 | SDGVKESSESTNTTIEDEDTK(5)-DLKPENLLLASKLK(12) | linker | kinase | 328-146 |

| | | | | |
|---|---|---|---|---|
| 147 | SDGVKESSESTNTTIEDEDTK(5)-ESSESTNTTIEDEDTKVR(16) | linker | hub | 328-344 |
| 148 | SDGVKESSESTNTTIEDEDTK(5)-ITAAEALKHPWISHR(8) | linker | kinase | 328-267 |
| 149 | SDGVKESSESTNTTIEDEDTK(5)-KQEIIK(1) | linker | hub | 328-347 |
| 150 | SDGVKESSESTNTTIEDEDTK(5)-KSDGVK(1) | linker | linker | 328-323 |
| 151 | SDGVKESSESTNTTIEDEDTK(5)-LKGAAVK(2) | linker | kinase | 328-148 |
| 152 | SDGVKESSESTNTTIEDEDTK(5)-LKGAILTTMLATR(2) | linker | regulatory | 328-300 |
| 153 | SDGVKESSESTNTTIEDEDTK(5)-LLKHPNIVR(3) | linker | kinase | 328-68 |
| 154 | SDGVKESSESTNTTIEDEDTK(5)-MLTINPSKR(8) | linker | kinase | 328-258 |
| 155 | SDGVKESSESTNTTIEDEDTK(5)-NFSGGKSGGNK(6) | linker | linker | 328-317 |
| 156 | SDGVKESSESTNTTIEDEDTK(5)-NFSGGKSGGNKK(6) | linker | linker | 328-317 |
| 157 | SDGVKESSESTNTTIEDEDTK(5)-RKLKGAILTTMLATR(2) | linker | regulatory | 328-298 |
| 158 | SDGVKESSESTNTTIEDEDTK(5)-VLAGQEYAAKIINTK(10) | linker | kinase | 328-42 |
| 159 | VLAGQEYAAKIINTK(10)-DHQKLER(4) | kinase | kinase | 42-56 |
| 160 | VLAGQEYAAKIINTK(10)-DLKPENLLLASK(3) | kinase | kinase | 42-137 |
| 161 | VLAGQEYAAKIINTK(10)-DLKPENLLLASKLK(12) | kinase | kinase | 42-146 |
| 162 | VLAGQEYAAKIINTK(10)-KSDGVK(1) | kinase | linker | 42-323 |
| 163 | VLAGQEYAAKIINTK(10)-LKGAAVK(2) | kinase | kinase | 42-148 |
| 164 | VLAGQEYAAKIINTK(10)-LKGAILTTMLATR(2) | kinase | regulatory | 42-300 |
| 165 | VLAGQEYAAKIINTK(10)-LLKHPNIVR(3) | kinase | kinase | 42-68 |
| 166 | VLAGQEYAAKIINTK(10)-MLTINPSKR(8) | kinase | kinase | 42-258 |
| 167 | VLAGQEYAAKIINTK(10)-NFSGGKSGGNK(6) | kinase | linker | 42-317 |
| 168 | VLAGQEYAAKIINTKK(10)-DHQKLER(4) | kinase | kinase | 42-56 |

Supplementary Table 3

| Heterotypic crosslinks (basal, 30 min) | | | | | | |
|---|---|---|---|---|---|---|
| # | Peptides | domain_a | domain_b | crosslinked lysines | R sample 1 | R sample 2 |
| 1 | DGKWQIVHFHR(3)-DLKPENLLLASK(3) | hub | kinase | 461-137 | 0,15 | 0,12 |
| 2 | LLKHPNIVR(3)-KQEIIK(1) | kinase | hub | 68-347 | 0,08 | 0,13 |
| 3 | ESSESTNTTIEDEDTKVR(16)-IINTKK(5) | hub | kinase | 344-47 | 0,02 | 0,05 |

| Heterotypic crosslinks (basal, 150 min) | | | | | | |
|---|---|---|---|---|---|---|
| **#** | **Peptides** | **domain_a** | **domain_b** | **crosslinked lysines** | **R sample 1** | **R sample 2** |
| 1 | DGKWQIVHFHR(3)-DLKPENLLLASK(3) | hub | kinase | 461-137 | 0,12 | 0,09 |
| 2 | DLKPENLLLASK(3)-KQEIIK(1) | kinase | hub | 137-347 | 0,09 | 0,08 |
| 3 | DLKPENLLLASKLK(12)-DLINKMLTINPSK(5) | kinase | kinase | 146-250 | 0,05 | 0,08 |
| 4 | DLKPENLLLASKLK(12)-DLKPENLLLASK(3) | kinase | kinase | 146-137 | 0,10 | 0,08 |
| 5 | DLKPENLLLASKLK(12)-IINTKK(5) | kinase | kinase | 146-47 | 0,06 | 0,07 |
| 6 | ESSESTNTTIEDEDTKVR(16)-DLKPENLLLASK(3) | hub | kinase | 344-137 | 0,09 | 0,13 |
| 7 | ITAAEALKHPWISHR(8)-DLKPENLLLASKLK(12) | kinase | kinase | 267-146 | 0,07 | 0,28 |
| 8 | VLAGQEYAAKIINTK(10)-DLINKMLTINPSK(5) | kinase | kinase | 42-250 | 0,06 | 0,06 |

Supplementary Table 5

| Homotypic crosslinks (activated, 30 min) | | | | |
|---|---|---|---|---|
| # | Peptide | domain_a | domain_b | crosslinked lysines |
| 1 | AGAYDFPSPEWDTVTPEAKDLINK(19)-DHQKLER(4) | kinase | kinase | 245-56 |
| 2 | AGAYDFPSPEWDTVTPEAKDLINK(19)-DLKPENLLLASK(3) | kinase | kinase | 245-137 |
| 3 | AGAYDFPSPEWDTVTPEAKDLINK(19)-ITAAEALKHPWISHR(8) | kinase | kinase | 245-267 |
| 4 | AGAYDFPSPEWDTVTPEAKDLINK(19)-KQEIIK(1) | kinase | hub | 245-347 |
| 5 | CVKVLAGQEYAAK(3)-IINTKK(5) | kinase | kinase | 32-47 |
| 6 | CVKVLAGQEYAAK(3)-LLKHPNIVR(3) | kinase | kinase | 32-68 |
| 7 | DGKWQIVHFHR(3)-KQEIIK(1) | hub | hub | 461-347 |
| 8 | DGKWQIVHFHR(3)-KSDGVK(1) | hub | linker | 461-323 |
| 9 | DGKWQIVHFHR(3)-NFSGGKSGGNK(6) | hub | linker | 461-317 |
| 10 | DHQKLER(4)-DHQKLER(4) | kinase | kinase | 56-56 |
| 11 | DHQKLER(4)-IINTKK(5) | kinase | kinase | 56-47 |
| 12 | DLINKMLTINPSK(5)-DLKPENLLLASK(3) | kinase | kinase | 250-137 |
| 13 | DLKPENLLLASK(3)-DHQKLER(4) | kinase | kinase | 137-56 |
| 14 | DLKPENLLLASK(3)-DLKPENLLLASK(3) | kinase | kinase | 137-137 |
| 15 | DLKPENLLLASK(3)-KSDGVK(1) | kinase | linker | 137-323 |
| 16 | DLKPENLLLASK(3)-LKGAAVK(2) | kinase | kinase | 137-148 |
| 17 | DLKPENLLLASK(3)-MLTINPSKR(8) | kinase | kinase | 137-258 |
| 18 | DLKPENLLLASK(3)-NFSGGKSGGNK(6) | kinase | linker | 137-317 |
| 19 | DLKPENLLLASKLK(12)-DHQKLER(4) | kinase | kinase | 146-56 |
| 20 | DLKPENLLLASKLK(12)-KSDGVK(1) | kinase | linker | 146-323 |
| 21 | DLKPENLLLASKLK(12)-LLKHPNIVR(3) | kinase | kinase | 146-68 |
| 22 | ESSESTNTTIEDEDTKVR(16)-DGKWQIVHFHR(3) | hub | hub | 344-461 |
| 23 | ESSESTNTTIEDEDTKVR(16)-ESSESTNTTIEDEDTKVR(16) | hub | hub | 344-344 |
| 24 | ESSESTNTTIEDEDTKVR(16)-KQEIIK(1) | hub | hub | 344-347 |
| 25 | ESSESTNTTIEDEDTKVR(16)-KSDGVK(1) | hub | linker | 344-323 |
| 26 | ESSESTNTTIEDEDTKVR(16)-NFSGGKSGGNK(6) | hub | linker | 344-317 |
| 27 | ESSESTNTTIEDEDTKVRK(16)-ESSESTNTTIEDEDTKVR(16) | hub | hub | 344-344 |
| 28 | ESSESTNTTIEDEDTKVRK(16)-KQEIIK(1) | hub | hub | 344-347 |
| 29 | IINTKK(5)-IINTKK(5) | kinase | kinase | 47-47 |
| 30 | IINTKK(5)-LKGAAVK(2) | kinase | kinase | 47-148 |
| 31 | ITAAEALKHPWISHR(8)-DLINKMLTINPSK(5) | kinase | kinase | 267-250 |
| 32 | ITAAEALKHPWISHR(8)-DLKPENLLLASKLK(12) | kinase | kinase | 267-146 |
| 33 | ITAAEALKHPWISHR(8)-ITAAEALKHPWISHR(8) | kinase | kinase | 267-267 |
| 34 | ITAAEALKHPWISHR(8)-KQEIIK(1) | kinase | hub | 267-347 |
| 35 | ITAAEALKHPWISHR(8)-KSDGVK(1) | kinase | linker | 267-323 |

| | | | | |
|---|---|---|---|---|
| 36 | ITAAEALKHPWISHR(8)-LKGAAVK(2) | kinase | kinase | 267-148 |
| 37 | ITAAEALKHPWISHR(8)-MLTINPSKR(8) | kinase | kinase | 267-258 |
| 38 | ITAAEALKHPWISHR(8)-NFSGGKSGGNK(6) | kinase | linker | 267-317 |
| 39 | KQEIIK(1)-IINTKK(5) | hub | kinase | 347-47 |
| 40 | KQEIIK(1)-KSDGVK(1) | hub | linker | 347-323 |
| 41 | KQEIIK(1)-LKGAAVK(2) | hub | kinase | 347-148 |
| 42 | KSDGVKESSESTNTTIEDEDTK(6)-KQEIIK(1) | linker | hub | 328-347 |
| 43 | KSDGVKESSESTNTTIEDEDTK(6)-NFSGGKSGGNK(6) | linker | linker | 328-317 |
| 44 | LKGAAVK(2)-LKGAAVK(2) | kinase | kinase | 148-148 |
| 45 | LKGAILTTMLATR(2)-DHQKLER(4) | regulatory | kinase | 300-56 |
| 46 | LKGAILTTMLATR(2)-KSDGVK(1) | regulatory | linker | 300-323 |
| 47 | LKGAILTTMLATR(2)-NFSGGKSGGNK(6) | regulatory | linker | 300-317 |
| 48 | LLKHPNIVR(3)-DHQKLER(4) | kinase | kinase | 68-56 |
| 49 | LLKHPNIVR(3)-KQEIIK(1) | kinase | hub | 68-347 |
| 50 | LLKHPNIVR(3)-LLKHPNIVR(3) | kinase | kinase | 68-68 |
| 51 | LYQQIKAGAYDFPSPEWDTVTPEAK(6)-DHQKLER(4) | kinase | kinase | 226-56 |
| 52 | LYQQIKAGAYDFPSPEWDTVTPEAK(6)-DLKPENLLLASKLK(12) | kinase | kinase | 226-146 |
| 53 | LYQQIKAGAYDFPSPEWDTVTPEAK(6)-IINTKK(5) | kinase | kinase | 226-47 |
| 54 | LYQQIKAGAYDFPSPEWDTVTPEAK(6)-KSDGVK(1) | kinase | linker | 226-323 |
| 55 | LYQQIKAGAYDFPSPEWDTVTPEAK(6)-LYQQIKAGAYDFPSPEWDTVTPEAK(6) | kinase | kinase | 226-226 |
| 56 | LYQQIKAGAYDFPSPEWDTVTPEAK(6)-MLTINPSKR(8) | kinase | kinase | 226-258 |
| 57 | LYQQIKAGAYDFPSPEWDTVTPEAK(6)-NFSGGKSGGNK(6) | kinase | linker | 226-317 |
| 58 | MLTINPSKR(8)-DHQKLER(4) | kinase | kinase | 258-56 |
| 59 | MLTINPSKR(8)-IINTKK(5) | kinase | kinase | 258-47 |
| 60 | MLTINPSKR(8)-KQEIIK(1) | kinase | hub | 258-347 |
| 61 | MLTINPSKR(8)-KSDGVK(1) | kinase | linker | 258-323 |
| 62 | MLTINPSKR(8)-LKGAAVK(2) | kinase | kinase | 258-148 |
| 63 | MLTINPSKR(8)-MLTINPSKR(8) | kinase | kinase | 258-258 |
| 64 | MLTINPSKR(8)-NFSGGKSGGNK(6) | kinase | linker | 258-317 |
| 65 | NFSGGKSGGNK(6)-KQEIIK(1) | linker | hub | 317-347 |
| 66 | NFSGGKSGGNK(6)-KSDGVK(1) | linker | linker | 317-323 |
| 67 | NFSGGKSGGNK(6)-NFSGGKSGGNK(6) | linker | linker | 317-317 |
| 68 | QETVDCLKK(8)-LLKHPNIVR(3) | regulatory | kinase | 291-68 |
| 69 | RITAAEALKHPWISHR(9)-DLINKMLTINPSKR(13) | kinase | kinase | 267-258 |
| 70 | RITAAEALKHPWISHR(9)-DLINKMLTINPSKR(5) | kinase | kinase | 267-250 |
| 71 | SDGVKESSESTNTTIEDEDTK(5)-DGKWQIVHFHR(3) | linker | hub | 328-461 |
| 72 | SDGVKESSESTNTTIEDEDTK(5)-ESSESTNTTIEDEDTKVR(16) | linker | HUB | 328-344 |

| | | | | |
|---|---|---|---|---|
| 73 | SDGVKESSESTNTTIEDEDTK(5)-KQEIIK(1) | linker | hub | 328-347 |
| 74 | SDGVKESSESTNTTIEDEDTK(5)-KSDGVK(1) | linker | linker | 328-323 |
| 75 | SDGVKESSESTNTTIEDEDTK(5)-LKGAAVK(2) | linker | kinase | 328-148 |
| 76 | SDGVKESSESTNTTIEDEDTK(5)-LLKHPNIVR(3) | linker | kinase | 328-68 |
| 77 | SDGVKESSESTNTTIEDEDTK(5)-MLTINPSKR(8) | linker | kinase | 328-258 |
| 78 | SDGVKESSESTNTTIEDEDTK(5)-NFSGGKSGGNK(6) | linker | linker | 328-317 |
| 79 | SDGVKESSESTNTTIEDEDTK(5)-NFSGGKSGGNKK(6) | linker | linker | 328-317 |
| 80 | SDGVKESSESTNTTIEDEDTK(5)-SDGVKESSESTNTTIEDEDTK(5) | linker | linker | 328-328 |
| 81 | SDGVKESSESTNTTIEDEDTKVR(21)-KQEIIK(1) | hub | hub | 344-347 |
| 82 | VLAGQEYAAKIINTK(10)-DHQKLER(4) | kinase | kinase | 42-56 |
| 83 | VLAGQEYAAKIINTK(10)-LLKHPNIVR(3) | kinase | kinase | 42-68 |
| 84 | VLAGQEYAAKIINTK(10)-MLTINPSKR(8) | kinase | kinase | 42-258 |
| 85 | VLAGQEYAAKIINTK(10)-VLAGQEYAAKIINTK(10) | kinase | kinase | 42-42 |

Supplementary Table 6

| Homotypic crosslinks (activated, 150 min) | | | | |
|---|---|---|---|---|
| # | Peptide | domain_a | domain_b | crosslinked lysines |
| 1 | AGAYDFPSPEWDTVTPEAKDLINK(19)-DGKWQIVHFHR(3) | kinase | hub | 245-461 |
| 2 | AGAYDFPSPEWDTVTPEAKDLINK(19)-ITAAEALKHPWISHR(8) | kinase | kinase | 245-267 |
| 3 | AGAYDFPSPEWDTVTPEAKDLINK(19)-MLTINPSKR(8) | kinase | kinase | 245-258 |
| 4 | AGAYDFPSPEWDTVTPEAKDLINK(19)-NFSGGKSGGNK(6) | kinase | linker | 245-317 |
| 5 | CVKVLAGQEYAAK(3)-IINTKK(5) | kinase | kinase | 32-47 |
| 6 | DGKWQIVHFHR(3)-KQEIIK(1) | hub | hub | 461-347 |
| 7 | DGKWQIVHFHR(3)-KSDGVK(1) | hub | linker | 461-323 |
| 8 | DGKWQIVHFHR(3)-NFSGGKSGGNK(6) | hub | linker | 461-317 |
| 9 | DHQKLER(4)-DHQKLER(4) | kinase | kinase | 56-56 |
| 10 | DHQKLER(4)-IINTKK(5) | kinase | kinase | 56-47 |
| 11 | DLINKMLTINPSK(5)-DHQKLER(4) | kinase | kinase | 250-56 |
| 12 | DLINKMLTINPSK(5)-DLINKMLTINPSK(5) | kinase | kinase | 250-250 |
| 13 | DLINKMLTINPSK(5)-NFSGGKSGGNK(6) | kinase | linker | 250-317 |
| 14 | DLINKMLTINPSKR(5)-DLINKMLTINPSKR(13) | kinase | kinase | 250-258 |
| 15 | DLKPENLLLASK(3)-DLKPENLLLASK(3) | kinase | kinase | 137-137 |
| 16 | DLKPENLLLASK(3)-LKGAAVK(2) | kinase | kinase | 137-148 |
| 17 | DLKPENLLLASK(3)-NFSGGKSGGNK(6) | kinase | linker | 137-317 |
| 18 | DLKPENLLLASKLK(12)-DHQKLER(4) | kinase | kinase | 146-56 |
| 19 | DLKPENLLLASKLK(12)-DLKPENLLLASKLK(12) | kinase | kinase | 146-146 |
| 20 | ESSESTNTTIEDEDTKVR(16)-DGKWQIVHFHR(3) | hub | hub | 344-461 |
| 21 | ESSESTNTTIEDEDTKVR(16)-ESSESTNTTIEDEDTKVR(16) | hub | hub | 344-344 |
| 22 | ESSESTNTTIEDEDTKVR(16)-ITAAEALKHPWISHR(8) | hub | kinase | 344-267 |
| 23 | ESSESTNTTIEDEDTKVR(16)-KQEIIK(1) | hub | hub | 344-347 |
| 24 | ESSESTNTTIEDEDTKVR(16)-KSDGVK(1) | hub | linker | 344-323 |
| 25 | ESSESTNTTIEDEDTKVR(16)-LLKHPNIVR(3) | hub | kinase | 344-68 |
| 26 | ESSESTNTTIEDEDTKVR(16)-MLTINPSKR(8) | hub | kinase | 344-258 |
| 27 | ESSESTNTTIEDEDTKVR(16)-NFSGGKSGGNK(6) | hub | linker | 344-317 |
| 28 | ESSESTNTTIEDEDTKVRK(16)-KQEIIK(1) | hub | hub | 344-347 |
| 29 | IINTKK(5)-IINTKK(5) | kinase | kinase | 47-47 |
| 30 | IINTKK(5)-LKGAAVK(2) | kinase | kinase | 47-148 |
| 31 | ITAAEALKHPWISHR(8)-DHQKLER(4) | kinase | kinase | 267-56 |
| 32 | ITAAEALKHPWISHR(8)-DLINKMLTINPSK(5) | kinase | kinase | 267-250 |
| 33 | ITAAEALKHPWISHR(8)-DLKPENLLLASK(3) | kinase | kinase | 267-137 |
| 34 | ITAAEALKHPWISHR(8)-KQEIIK(1) | kinase | hub | 267-347 |
| 35 | ITAAEALKHPWISHR(8)-KSDGVK(1) | kinase | linker | 267-323 |

| | | | | |
|---|---|---|---|---|
| 36 | ITAAEALKHPWISHR(8)-MLTINPSKR(8) | kinase | kinase | 267-258 |
| 37 | ITAAEALKHPWISHR(8)-NFSGGKSGGNK(6) | kinase | linker | 267-317 |
| 38 | KQEIIK(1)-KQEIIK(1) | hub | hub | 347-347 |
| 39 | KQEIIK(1)-KSDGVK(1) | hub | linker | 347-323 |
| 40 | KQEIIK(1)-LKGAAVK(2) | hub | kinase | 347-148 |
| 41 | KQEIIKVTEQLIEAISNGGFESYTK(1)-RDGKWQIVHFHR(4) | hub | hub | 347-461 |
| 42 | KQEIIKVTEQLIEAISNGGFESYTK(6)-ESSESTNTTIEDEDTKVR(16) | hub | hub | 352-344 |
| 43 | KQEIIKVTEQLIEAISNGGFESYTK(6)-KSDGVK(1) | hub | linker | 352-323 |
| 44 | KSDGVKESSESTNTTIEDEDTK(6)-NFSGGKSGGNK(6) | linker | linker | 328-317 |
| 45 | LKGAILTTMLATR(2)-KQEIIK(1) | regulatory | hub | 300-347 |
| 46 | LKGAILTTMLATR(2)-KSDGVK(1) | regulatory | linker | 300-323 |
| 47 | LKGAILTTMLATR(2)-NFSGGKSGGNK(6) | regulatory | linker | 300-317 |
| 48 | LKGAILTTMLATR(2)-QETVDCLKK(8) | regulatory | regulatory | 300-291 |
| 49 | LLKHPNIVR(3)-DHQKLER(4) | kinase | kinase | 68-56 |
| 50 | LLKHPNIVR(3)-NFSGGKSGGNK(6) | kinase | linker | 68-317 |
| 51 | LYQQIKAGAYDFPSPEWDTVTPEAK(6)-DGKWQIVHFHR(3) | kinase | hub | 226-461 |
| 52 | LYQQIKAGAYDFPSPEWDTVTPEAK(6)-DHQKLER(4) | kinase | kinase | 226-56 |
| 53 | LYQQIKAGAYDFPSPEWDTVTPEAK(6)-DLINKMLTINPSK(5) | kinase | kinase | 226-250 |
| 54 | LYQQIKAGAYDFPSPEWDTVTPEAK(6)-ITAAEALKHPWISHR(8) | kinase | kinase | 226-267 |
| 55 | LYQQIKAGAYDFPSPEWDTVTPEAK(6)-LYQQIKAGAYDFPSPEWDTVTPEAK(6) | kinase | kinase | 226-226 |
| 56 | LYQQIKAGAYDFPSPEWDTVTPEAK(6)-MLTINPSKR(8) | kinase | kinase | 226-258 |
| 57 | LYQQIKAGAYDFPSPEWDTVTPEAK(6)-NFSGGKSGGNK(6) | kinase | linker | 226-317 |
| 58 | MLTINPSKR(8)-DHQKLER(4) | kinase | kinase | 258-56 |
| 59 | MLTINPSKR(8)-KQEIIK(1) | kinase | hub | 258-347 |
| 60 | MLTINPSKR(8)-KSDGVK(1) | kinase | linker | 258-323 |
| 61 | MLTINPSKR(8)-LKGAAVK(2) | kinase | kinase | 258-148 |
| 62 | MLTINPSKR(8)-MLTINPSKR(8) | kinase | kinase | 258-258 |
| 63 | MLTINPSKR(8)-NFSGGKSGGNK(6) | kinase | linker | 258-317 |
| 64 | NFSGGKSGGNK(6)-KQEIIK(1) | linker | hub | 317-347 |
| 65 | NFSGGKSGGNK(6)-KSDGVK(1) | linker | linker | 317-323 |
| 66 | NFSGGKSGGNK(6)-LKGAAVK(2) | linker | kinase | 317-148 |
| 67 | NFSGGKSGGNK(6)-NFSGGKSGGNK(6) | linker | linker | 317-317 |
| 68 | QEIIKVTEQLIEAISNGGFESYTK(5)-DGKWQIVHFHR(3) | hub | hub | 352-461 |
| 69 | RITAAEALKHPWISHR(9)-DLINKMLTINPSKR(13) | kinase | kinase | 267-258 |
| 70 | RITAAEALKHPWISHR(9)-DLINKMLTINPSKR(5) | kinase | kinase | 267-250 |
| 71 | SDGVKESSESTNTTIEDEDTK(5)-DGKWQIVHFHR(3) | linker | hub | 328-461 |
| 72 | SDGVKESSESTNTTIEDEDTK(5)-DHQKLER(4) | linker | kinase | 328-56 |

| | | | | |
|---|---|---|---|---|
| 73 | SDGVKESSESTNTTIEDEDTK(5)-DLINKMLTINPSK(5) | linker | kinase | 328-250 |
| 74 | SDGVKESSESTNTTIEDEDTK(5)-ESSESTNTTIEDEDTKVR(16) | linker | hub | 328-344 |
| 75 | SDGVKESSESTNTTIEDEDTK(5)-ITAAEALKHPWISHR(8) | linker | kinase | 328-267 |
| 76 | SDGVKESSESTNTTIEDEDTK(5)-KQEIIK(1) | linker | hub | 328-347 |
| 77 | SDGVKESSESTNTTIEDEDTK(5)-KSDGVK(1) | linker | linker | 328-323 |
| 78 | SDGVKESSESTNTTIEDEDTK(5)-LKGAILTTMLATR(2) | linker | regulatory | 328-300 |
| 79 | SDGVKESSESTNTTIEDEDTK(5)-LLKHPNIVR(3) | linker | kinase | 328-68 |
| 80 | SDGVKESSESTNTTIEDEDTK(5)-MLTINPSKR(8) | linker | kinase | 328-258 |
| 81 | SDGVKESSESTNTTIEDEDTK(5)-NFSGGKSGGNK(6) | linker | linker | 328-317 |
| 82 | SDGVKESSESTNTTIEDEDTK(5)-NFSGGKSGGNKK(6) | linker | linker | 328-317 |
| 83 | SDGVKESSESTNTTIEDEDTK(5)-RKLKGAILTTMLATR(2) | linker | regulatory | 328-298 |
| 84 | SDGVKESSESTNTTIEDEDTK(5)-SDGVKESSESTNTTIEDEDTK(5) | linker | linker | 328-328 |
| 85 | VLAGQEYAAKIINTK(10)-DHQKLER(4) | kinase | kinase | 42-56 |
| 86 | VLAGQEYAAKIINTK(10)-VLAGQEYAAKIINTK(10) | kinase | kinase | 42-42 |

Supplementary Table 7

| Heterotypic crosslinks (activated, 30 min) | | | | | | |
|---|---|---|---|---|---|---|
| # | Peptides | domain_a | domain_b | crosslinked lysines | R sample 1 | R sample 2 |
| 1 | DLKPENLLLASK(3)-IINTKK(5) | kinase | kinase | 137-47 | 1,63 | 0,22 |
| 2 | FTEEYQLFEELGKGAFSVVR(13)-MLTINPSKR(8) | kinase | kinase | 21-258 | 0,55 | 0,15 |
| 3 | VLAGQEYAAKIINTK(10)-DLINKMLTINPSK(5) | kinase | kinase | 42-250 | 0,35 | 0,34 |
| 4 | DLKPENLLLASKLK(12)-DLKPENLLLASK(3) | kinase | kinase | 146-137 | 0,34 | 0,22 |
| 5 | ESSESTNTTIEDEDTKVR(16)-DLKPENLLLASK(3) | hub | kinase | 344-137 | 0,33 | 0,19 |
| 6 | ITAAEALKHPWISHR(8)-VLAGQEYAAKIINTK(10) | kinase | kinase | 267-42 | 0,29 | 0,24 |
| 7 | NFSGGKSGGNK(6)-DHQKLER(4) | linker | kinase | 317-56 | 0,27 | 0,21 |
| 8 | LKGAILTTMLATR(2)-DLKPENLLLASK(3) | regulatory | kinase | 300-137 | 0,26 | 0,12 |
| 9 | MLTINPSKR(8)-LLKHPNIVR(3) | kinase | kinase | 258-68 | 0,25 | 0,19 |
| 10 | LYQQIKAGAYDFPSPEWDTVTPEAK(6)-VLAGQEYAAKIINTK(10) | kinase | kinase | 226-42 | 0,24 | 0,15 |
| 11 | FTEEYQLFEELGKGAFSVVR(13)-LLKHPNIVR(3) | kinase | kinase | 21-68 | 0,23 | 0,17 |
| 12 | FTEEYQLFEELGKGAFSVVR(13)-LKGAAVK(2) | kinase | kinase | 21-148 | 0,23 | 0,20 |
| 13 | ITAAEALKHPWISHR(8)-IINTKK(5) | kinase | kinase | 267-47 | 0,22 | 0,18 |
| 14 | DLKPENLLLASKLK(12)-LKGAAVK(2) | kinase | kinase | 146-148 | 0,22 | 0,27 |
| 15 | LLKHPNIVR(3)-IINTKK(5) | kinase | kinase | 68-47 | 0,21 | 0,22 |
| 16 | AGAYDFPSPEWDTVTPEAKDLINK(19)-LLKHPNIVR(3) | kinase | kinase | 245-68 | 0,19 | 0,35 |
| 17 | DHQKLER(4)-KSDGVK(1) | kinase | linker | 56-323 | 0,16 | 0,17 |
| 18 | FTEEYQLFEELGKGAFSVVR(13)-DHQKLER(4) | kinase | kinase | 21-56 | 0,15 | 0,12 |
| 19 | VLAGQEYAAKIINTK(10)-DLKPENLLLASK(3) | kinase | kinase | 42-137 | 0,15 | 0,21 |
| 20 | ESSESTNTTIEDEDTKVR(16)-ITAAEALKHPWISHR(8) | hub | kinase | 344-267 | 0,05 | 0,12 |

Supplementary Table 8

| Heterotypic crosslinks (activated, 150 min) | | | | | | |
|---|---|---|---|---|---|---|
| **#** | **Peptides** | **domain_a** | **domain_b** | **crosslinked lysines** | **R sample 1** | **R sample 2** |
| 1 | VLAGQEYAAKIINTK(10)-DLKPENLLLASKLK(12) | kinase | kinase | 42-146 | 0,56 | 0,40 |
| 2 | FTEEYQLFEELGKGAFSVVR(13)-ITAAEALKHPWISHR(8) | kinase | kinase | 21-267 | 0,48 | 0,45 |
| 3 | DLKPENLLLASKLK(12)-DLINKMLTINPSK(5) | kinase | kinase | 146-250 | 0,41 | 0,34 |
| 4 | DLKPENLLLASK(3)-LLKHPNIVR(3) | kinase | kinase | 137-68 | 0,37 | 0,30 |
| 5 | DLKPENLLLASKLK(12)-DLKPENLLLASK(3) | kinase | kinase | 146-137 | 0,37 | 0,29 |
| 6 | FTEEYQLFEELGKGAFSVVR(13)-DLINKMLTINPSK(5) | kinase | kinase | 21-250 | 0,36 | 0,24 |
| 7 | FTEEYQLFEELGKGAFSVVR(13)-VLAGQEYAAKIINTK(10) | kinase | kinase | 21-42 | 0,33 | 0,24 |
| 8 | MLTINPSKR(8)-GSGSGMTR(1) | kinase | kinase | 258-1 | 0,33 | 0,20 |
| 9 | NFSGGKSGGNK(6)-DHQKLER(4) | linker | kinase | 317-56 | 0,31 | 0,27 |
| 10 | DLINKMLTINPSK(5)-IINTKK(5) | kinase | kinase | 250-47 | 0,30 | 0,19 |
| 11 | ITAAEALKHPWISHR(8)-VLAGQEYAAKIINTK(10) | kinase | kinase | 267-42 | 0,30 | 0,25 |
| 12 | FTEEYQLFEELGKGAFSVVR(13)-MLTINPSKR(8) | kinase | kinase | 21-258 | 0,30 | 0,24 |
| 13 | VLAGQEYAAKIINTK(10)-IINTKK(5) | kinase | kinase | 42-47 | 0,29 | 0,23 |
| 14 | FTEEYQLFEELGKGAFSVVR(13)-LKGAAVK(2) | kinase | kinase | 21-148 | 0,29 | 0,21 |
| 15 | DHQKLER(4)-KQEIIK(1) | kinase | hub | 56-347 | 0,28 | 0,15 |
| 16 | LLKHPNIVR(3)-KQEIIK(1) | kinase | hub | 68-347 | 0,28 | 0,12 |
| 17 | FTEEYQLFEELGKGAFSVVR(13)-DLKPENLLLASK(3) | kinase | kinase | 21-137 | 0,27 | 0,23 |
| 18 | AGAYDFPSPEWDTVTPEAKDLINK(19)-DHQKLER(4) | kinase | kinase | 245-56 | 0,26 | 0,17 |
| 19 | ESSESTNTTIEDEDTKVR(16)-DLKPENLLLASK(3) | hub | kinase | 344-137 | 0,26 | 0,18 |
| 20 | MLTINPSKR(8)-IINTKK(5) | kinase | kinase | 258-47 | 0,25 | 0,18 |
| 21 | FTEEYQLFEELGKGAFSVVR(13)-DLKPENLLLASKLK(12) | kinase | kinase | 21-146 | 0,25 | 0,15 |
| 22 | AGAYDFPSPEWDTVTPEAKDLINK(19)-GSGSGMTR(1) | kinase | kinase | 245-1 | 0,25 | 0,26 |
| 23 | LKGAILTTMLATR(2)-DHQKLER(4) | regulatory | kinase | 300-56 | 0,25 | 0,16 |
| 24 | VLAGQEYAAKIINTK(10)-MLTINPSKR(8) | kinase | kinase | 42-258 | 0,24 | 0,20 |
| 25 | LYQQIKAGAYDFPSPEWDTVTPEAK(6)-VLAGQEYAAKIINTK(10) | kinase | kinase | 226-42 | 0,24 | 0,25 |
| 26 | GSGSGMTR(1)-LKGAAVK(2) | kinase | kinase | 1-148 | 0,24 | 0,25 |
| 27 | AGAYDFPSPEWDTVTPEAKDLINK(19)-DLKPENLLLASK(3) | kinase | kinase | 245-137 | 0,23 | 0,19 |
| 28 | FTEEYQLFEELGKGAFSVVR(13)-KSDGVK(1) | kinase | linker | 21-323 | 0,23 | 0,20 |
| 29 | FTEEYQLFEELGKGAFSVVR(13)-LLKHPNIVR(3) | kinase | kinase | 21-68 | 0,23 | 0,21 |
| 30 | DLINKMLTINPSK(5)-LKGAAVK(2) | kinase | kinase | 250-148 | 0,23 | 0,15 |
| 31 | VLAGQEYAAKIINTK(10)-LKGAAVK(2) | kinase | kinase | 42-148 | 0,23 | 0,18 |
| 32 | VLAGQEYAAKIINTK(10)-NFSGGKSGGNK(6) | kinase | linker | 42-317 | 0,23 | 0,14 |
| 33 | VLAGQEYAAKIINTK(10)-DLKPENLLLASK(3) | kinase | kinase | 42-137 | 0,23 | 0,22 |
| 34 | ITAAEALKHPWISHR(8)-IINTKK(5) | kinase | kinase | 267-47 | 0,22 | 0,23 |
| 35 | LYQQIKAGAYDFPSPEWDTVTPEAK(6)-DLKPENLLLASK(3) | kinase | kinase | 226-137 | 0,22 | 0,17 |
| 36 | DLINKMLTINPSK(5)-DLKPENLLLASK(3) | kinase | kinase | 250-137 | 0,21 | 0,25 |
| 37 | LKGAILTTMLATR(2)-DLKPENLLLASK(3) | regulatory | kinase | 300-137 | 0,21 | 0,19 |
| 38 | AGAYDFPSPEWDTVTPEAKDLINK(19)-DLKPENLLLASKLK(12) | kinase | kinase | 245-146 | 0,20 | 0,16 |
| 39 | LYQQIKAGAYDFPSPEWDTVTPEAK(6)-DLKPENLLLASKLK(12) | kinase | kinase | 226-146 | 0,19 | 0,15 |
| 40 | LYQQIKAGAYDFPSPEWDTVTPEAK(6)-LKGAAVK(2) | kinase | kinase | 226-148 | 0,19 | 0,12 |
| 41 | LLKHPNIVR(3)-LKGAAVK(2) | kinase | kinase | 68-148 | 0,18 | 0,20 |

| | | | | | | |
|---|---|---|---|---|---|---|
| 42 | VLAGQEYAAKIINTK(10)-LLKHPNIVR(3) | kinase | kinase | 42-68 | 0,17 | 0,15 |
| 43 | DLKPENLLLASKLK(12)-MLTINPSKR(8) | kinase | kinase | 146-258 | 0,16 | 0,16 |
| 44 | VLAGQEYAAKIINTK(10)-KQEIIK(1) | kinase | hub | 42-347 | 0,16 | 0,15 |
| 45 | DHQKLER(4)-KSDGVK(1) | kinase | linker | 56-323 | 0,16 | 0,12 |
| 46 | VLAGQEYAAKIINTK(10)-KSDGVK(1) | kinase | linker | 42-323 | 0,14 | 0,15 |

| pT286 Heterotypic peptides (30 min) | | | | | |
|---|---|---|---|---|---|
| **#** | **Peptides** | **domain_a** | **domain_b** | **link** | **R** |
| 1 | QETVDCLKK(8)-LLKHPNIVR(3) | regulatory | kinase | 291-68 | 0,15 |
| 2 | LYQQIKAGAYDFPSPEWDTVTPEAK(6)-QETVDCLKK(8) | kinase | regulatory | 226-291 | 0,13 |
| 3 | VLAGQEYAAKIINTK(10)-QETVDCLKK(8) | kinase | regulatory | 42-291 | 0,10 |
| 4 | DLKPENLLLASKLK(12)-QETVDCLKK(8) | kinase | regulatory | 146-291 | 0,10 |
| 5 | QETVDCLKK(8)-MLTINPSKR(8) | regulatory | kinase | 291-258 | 0,07 |
| 6 | DLINKMLTINPSK(5)-QETVDCLKK(8) | kinase | regulatory | 250-291 | 0,05 |

Supplementary Table 10

| pT286 Heterotypic peptides (150 min) | | | | | |
|---|---|---|---|---|---|
| # | Peptides | domain_a | domain_b | crosslinked lysines | R |
| 1 | VLAGQEYAAKIINTK(10)-QETVDCLKK(8) | kinase | regulatory | 42-291 | 0,39 |
| 2 | QETVDCLKK(8)-LLKHPNIVR(3) | regulatory | kinase | 291-68 | 0,35 |
| 3 | QETVDCLKK(8)-DHQKLER(4) | regulatory | kinase | 291-56 | 0,26 |
| 4 | ITAAEALKHPWISHR(8)-QETVDCLKK(8) | kinase | regulatory | 267-291 | 0,18 |
| 5 | DLINKMLTINPSK(5)-QETVDCLKK(8) | kinase | regulatory | 250-291 | 0,15 |
| 6 | QETVDCLKK(8)-LKGAAVK(2) | regulatory | kinase | 291-148 | 0,11 |
